# Estimation of the propensity for sexual selection in a cyclical parthenogen

**DOI:** 10.1101/2020.02.05.935148

**Authors:** David Duneau, Florian Altermatt, Jean-Baptiste Ferdy, Frida Ben-Ami, Dieter Ebert

## Abstract

Cyclical parthenogenesis is a widespread reproductive strategy in which organisms go through one or multiple rounds of clonal reproduction before sexual reproduction. Because sexual reproduction is typically less common than parthenogenesis in populations of the planktonic cladoceran *Daphnia magna,* it is not frequently studied. Here we examine the sexual process of *D. magna* and its relation to sexual selection in *Daphnia* rockpool populations by observing natural mating in these shallow habitats where sex generally occurs throughout the summer. Although microsatellite markers were found to reveal no evidence of disassortative mating or, thus, of inbreeding avoidance, body length and infection status did reveal assortative mating, suggesting sexual selection to act. When two males mated with a single female, the larger male was observed to remain longer, possibly giving it an advantage in sperm competition. Indirect evidence points at the brood pouch as the likely site of fertilization and thus, sperm competition. Sperm length was as variable within ejaculates as it was among males from different populations. Our data provide firm evidence that sexual selection is present in this species, most likely manifesting itself through a combination of female choice and male–male competition.

## Introduction

Whenever sexual reproduction occurs, sexual selection operates to varying degrees on the ability to fertilize (Andersson 1994; Shuker 2010; Clutton-Brock 2017). Sexual selection generally acts on traits that evolve jointly but separately in each sex: For females this means female choice (for males or sperm), and for males this means traits that allow them to gain a competitive advantage either directly against each other or in the face of female choice. Females tend to choose for characteristics that maximize the success of their progeny, such as male vigor, size, ornament, and infection status. Males, on the other hand, tend to increase their mating success with condition-dependent traits, such as mate searching intensity, fighting ability, sperm quality or some types of exaggerated morphological characters (Andersson 1994; Morehouse 2014; Kaldun and Otti 2016; Houslay et al. 2017). Sexual selection thus has the potential to reduce mutation load—one of the frequently cited reasons why, despite its cost, it is maintained (Whitlock and Agrawal 2009; Lumley et al. 2015). In fact, the potential evolutionary benefits of sexual selection probably explain why organisms with sole asexual reproduction are extremely rare.

Organisms that alternate between sexual and asexual reproduction are said to reproduce by cyclical parthenogenesis. Examples are found in many taxa: aphids, stick insects, rotifers, parasitic nematodes of human, and even vertebrates (Hand 1991; Lampert 2009). In freshwater cladocerans from the genus *Daphnia*, periods of asexual reproduction are punctuated by sexual reproduction. These model crustaceans have played an important role in helping us understand fundamental biological concepts such as the capacity of immune cells to engulf foreign antigens (Metschnikoff 1884), the definition of germ line and soma (Weismann 1893), phenotypic plasticity (Wolterek 1909), the capacity of natural populations to genetically adapt to anthropogenic stressors (Jansen et al. 2011), parasite local adaptation (Reger et al. 2018), host–parasite coevolution (Decaestecker et al. 2007; Ebert et al. 2016), phenotypically plastic response to biotic and abiotic stressors (Tollrian and Heibl 2004; Cavalheri et al. 2019), and even the consequences of climate change on animal populations (George et al. 1990; Carter et al. 2017).

The overwhelming majority of studies that use *Daphnia*, however, focus on their asexual mode of reproduction. The biology of sexual reproduction in the genus *Daphnia*, including mating and fertilization, has largely been unexplored, and evidence for sexual selection in this genus has, to our knowledge, never been presented. The few studies that address sexual reproduction in *Daphnia* have been limited to laboratory conditions and have used different species with seemingly different ecologies (Brewer 1998; Winsor and Innes 2002; Wuerz et al. 2017). We argue that understanding sexual reproduction in cyclical parthenogenetic species such as *Daphnia* offers the opportunity to study the evolution of traits that are presumably under sexual selection after periods of clonal reproduction.

In *Daphnia*, sexual reproduction is linked to dormancy, as the two sexual eggs are protected by a hard melanized case known as an ephippium that allows embryos in developmental arrest to survive summer draughts and winter freezes. Sexual reproduction is key to the long-term persistence of *Daphnia* populations in unstable environments, creating genetically diverse egg-banks from which future populations are established. The frequency of sexual reproduction correlates with habitat instability at a continental scale (Roulin et al. 2013).

Sex is environmentally determined in *Daphnia* (Hobæk and Larsson 1990). Males and females are believed to be genetically identical (Hebert and Ward 1972), although their morphologies and swimming behaviors differ corresponding to their respective reproductive roles (Brewer 1998; Ebert 2005; Wuerz et al. 2017). Differences between the sexes is based on Darwin’s theory of sexual selection (Darwin 1871; Shine 1979; Clutton-Brock 2017) or on selection by intraspecific niche divergence (Cox and Calsbeek 2010; Law and Mehta 2018). As there is no indication that niches diverge between sexes in *Daphnia*, sexual selection may be the main factor that acts on sexually dimorphic traits; however, little is known about mating in *Daphnia*. Unlike copepods, (Lonsdale et al. 1998), there is no evidence that males use sex pheromones to find their mates (Crease and Hebert 1983; Winsor and Innes 2002), even though *Daphnia* respond behaviorally and phenotypically to several chemical cues such as fish kairomones (Hahn et al. 2019). Still, male swimming behavior seems to be optimized for finding an appropriate mate (Brewer 1998), and mating is not random, as males capture sexually active females (i.e. females carrying ephippial egg cases) more often than they capture other males or asexual females (Brewer 1998; Winsor and Innes 2002). Thus, it seems plausible that couples form in response to certain criteria, possibly reflecting individual quality. There is tremendous variation among *Daphnia* species in mating duration (from a few seconds up to a day (Forró 1997)), probably because the genus is believed to be older than 140 million years (Cornetti et al. 2019), predating the placental mammal diversification (Springer et al. 2003). Hence, it is difficult to generalize from one species to another. Here, we looked at the sexual interactions of males and females—from mate finding to the release of the fertilized resting egg—in natural populations of *Daphnia magna*. Our data provides evidence that sexual selection is present in this species and that it likely manifests itself in a combination of female choice and male–male competition.

## Materials and methods

### Study area

We studied a metapopulation of *Daphnia magna* on the coast of the Baltic Sea in southwestern Finland, near Tvärminne Zoological Station (59°50’ N, 23°15’E). The rockpools in this metapopulation are small (average volume about 300 L) and shallow (10 to 60 cm deep) (Altermatt and Ebert 2010), allowing easy access to every part of the habitat. About 40 % of them are inhabited with at least one *Daphnia* species (Pajunen 1986). Our fieldwork was performed over the course of four summers (2003, 2009, 2010 and 2011) and included 33 rockpool populations (See supplementary material 1 for further information).

### Data collection

To estimate the sex ratio in populations, we randomly collected planktonic *D. magna* with handheld nets (mesh size 0.3 mm) or from 1 L samples of water. The shallow pools allowed us to search and collect mating pairs with wide-mouth pipettes. Mating pairs were then kept separately in 25-mL jars and observed in 1-minute intervals. We recorded the number of males concomitantly attached to the female, the time period until male detachment from the female and the order of detachment when there was more than one male. Using a field dissecting scope, we observed the females and recorded the time post mating until they laid the sexual eggs in the brood pouch, which by this time had already assumed the typical shape of a resting egg case. Females were then kept in the jars until they dropped the resting egg case. Only males with an open chest (*i.e.* sign of being adult) were considered in the study. Females are more difficult to classify as adults but we discarded very small individuals to avoid none reproductive individuals. We measured body- and spine-length before checking for parasites under a microscope or storing them in ethanol at −20 °C.

The parasite we looked for was *Hamiltosporidium tvarminnensis*, a microsporidium commonly, but not exclusively, found in the studied metapopulation (Haag et al. 2011; Goren and Ben-Ami 2013). Although it infects several *Daphnia* species, its success and pathogenicity are very host specific (Vizoso and Ebert 2005; Sheikh-Jabbari et al. 2014; Urca and Ben-Ami 2018; Orlansky and Ben-Ami 2019). It has a mixed mode of transmission—either vertically or horizontally, when spores are released from the decaying cadaver (Lass and Ebert 2006).

To measure sperm length, we exposed males to a 1% nicotine solution, which stimulates muscle contractions, thereby causing a release of sperm (as in Duneau et al. 2012). As only mature sperm are in the testicular lumen (p. 11 in Wingstrand, 1978; p 277 in Zaffagnini, 1987), this method is better than crushing males, which would yield immature sperm of various length. Using a camera mounted on a microscope (magnification 200x), we then photographed and measured the longest length of several sperm in the sample with ImageJ (version 1.50i).

### Genotyping

To genotype individuals, we homogenized them individually in 100 μl of Chelex solution and followed a Chelex DNA extraction protocol (Walsh et al. 1991) before performing a PCR on four *Daphnia* microsatellite markers (see details in supplementary material 1 – section 5). PCR reactions of 5 μL were set up with the following cycling conditions: 95 °C for 15 min, followed by 30 cycles of 94 °C for 30 s, 60 °C for 1.5 min, 72 °C for 1.5 min, and 10 cycles of 94 °C for 30 s, 47 °C for 1.5 min, 72 °C for 1.5 min, and a final elongation step of 72 °C for 10 min. Genotyping was done on an *AB 3130xl Genetic Analyzer* (Applied Biosystems) using *genescan 500 LIZ* size standard (Applied Biosystems). Microsatellite alleles were scored using *genemapper* Software version 4.0 (ABI Prism).

### Coefficient of relatedness

Prior to the relatedness analysis, a simulation was performed to assess different estimators of relatedness coefficients (*r_xy_*) and determine the most appropriate estimator for our dataset. Given the allelic frequencies within the population (from the sample sizes n_Sp1-5_ = 264 and n_SP1-6_ = 262), 2,000 individual genotypes were simulated. From the simulated genotypes, 1,000 pairs (or comparisons between two simulated individuals) were drawn for four relationship categories: unrelated, half-siblings, full-siblings and parent-offspring. Then, *r_xy_* was calculated for each pair within those relationship categories using six different estimators (Lynch 1988; Queller and Goodnight 1989; Li et al. 1993; Ritland 1996; Lynch and Ritland 1999; Wang 2002; Milligan 2003) as described in (Wang 2011). All simulations and calculations of *r_xy_* for the empirical dataset were conducted using the package *related v0.8* (Pew et al. 2015) and the software Coancestry (Wang 2011) implemented in R. We used the Triadic Likelihood method (TrioML) to describe the coefficient of relatedness between mating males and females, as it was the most appropriate for describing the known relatedness in our simulated data. Using it, we compared the coefficient of relatedness of mating individuals to the coefficient of randomly associated male and female pairs from the same population.

### Paternity assessment

Based on the genotypes obtained for the coefficient of relatedness, we selected polyandrous matings to assess the paternity of each egg in the ephippia from microsatellites. We performed the same genotyping as above, but on the oocyte.

### Statistical analysis

All analyses were performed using R and Rstudio (RStudio Team 2016; R Core Team 2019). Supplementary material 1 was generated by Rmarkdown, a component of RStudio, and provided a summary data table, with all scripts and their associated analyses and plots, including supplementary figures. All the analysis and illustrations were done with the tidyverse R package suite (Wickham 2016; Wickham and Henry 2019; Wickham et al. 2019). We used the *Viridis* color palette to assure that plots would print well in grey scale and be readable for those with colorblindness (Garnier 2018). To illustrate the difference between factors in paired analysis, we also used the estimation graphic methods followed in Ho *et al.* (2019) with the package *dabestr v0.2.2*. This method uses bootstrap to estimate the difference between means and a 95 % confidence interval. Although not perfect to illustrate complex mixed models, it nonetheless helps represent the effect of paired comparison and indicates the level of confidence we can have in it. Odds ratios quantify the relation between two factors and typically quantify the effect of a variable.

Generalized mixed models were fitted using the function *fitme* from the package *spaMM v2.6.1* (Rousset and Ferdy 2014). This function allowed us to include random effect in the mixed model whenever necessary, notably to pair the variable by mating or pool it by population (with the argument “1|”), to nest variables (with the argument “/”), to specify the family of the random effect (with the argument “rand.family”) and to consider heteroscedasticity (with the argument “resid.model”). The significance of the factors in the model was tested using a likelihood ratio test, which compares the model with and without the variable of interest.

#### Sperm length

Based on AIC criteria, the sperm length was best fitted with a Gamma distribution. In males from the same mating, we tested for difference in sperm length between the first and second male to detach using the model: Sperm_length ~ Position_detached + (1|ID_mating), family=Gamma(link=“log”), rand.family= gaussian(“identity”).

In males from several lineages raised in laboratory conditions (AM-AR initially sampled from Armenia, CY-PA-1 from Cyprus, DE-Iinb1 from Germany and RU-KOR-1 from Russia), the model included the differences in variances among clones as follows: Sperm_length ~ Clone + (1|Clone/ID), resid.model= ~ Clone, family= Gamma(link=, “log“), rand.family= Gamma(link= “log”).

As sperm length varied considerably within an ejaculate, we explored this by fitting a gamma or a normal distribution on sperm length data for each individual male using the function *fitdist* from the package *fitdistrplus v1.0.14* (Delignette-Muller and Dutang 2015). We then tested the goodness of fit of this distribution with the function *gofstat* from the same package. The final AIC for each distribution was obtained by summing the AICs for each male. To test whether there might potentially be two sub-populations of sperms inside one ejaculate, we compared the AIC of the best model to that of a mixed model considering two gaussian distributions. The fit was performed with the function *densityMclust* from the package *mclust v5.4.5* (Scrucca et al. 2016), and the AIC was calculated.

#### Body length

The body length of mating males was best fit with a Gaussian distribution and by considering the difference in variance among populations. To test if there was a difference between males in the same mating, we used the following full model: Body_length ~ as.factor(Nbr_of_males) + (1|Population), resid.model= ~ Population, family= gaussian(link=identity), rand.family= gaussian(link=identity).

The relation between spine and body length of males and females was also best fit with a Gaussian distribution and by considering the difference in variance among populations. To test for a difference between mating and single individuals in the population, we used the following model: Spine_length ~ Body_length + Sex + Mating_status + (1|Population), resid.model= ~ Population, family=gaussian(link=identity). The significance of the factors in the model was tested with a likelihood ratio test, which compares the model with and without “Mating status” as a variable.

#### Infectious status

The sex-ratio of the 27 populations in relation to *H. tvaerminennsis* prevalence in single females was best fitted with a Gaussian distribution. The full model to test for correlation with prevalence in single females was as follows: Sex_ratio ~ Population_size + Prevalence_Female, family= gaussian(link=identity). The significance of the factors in the model was tested using a likelihood ratio test, which compares the model with and without “Prevalence_Female” as a variable.

The prevalence in males and females, both single or sexually active, was best fitted with a binomial distribution, noting the presence/absence (1 vs 0) of each individual. “Population” was considered as a random effect in order to pair the analysis. The full model to test for prevalence differences between sexes was as follows: Infectious_status ~ Sex + (1|Population), family=binomial(link=“logit”), either with single individuals only or with mating individuals only.

We used the same approach to test for difference in prevalence between mating and non mating individuals, using the full model Infectious_status ~ Mating_status + (1|Population), family=binomial(link=“logit”), rand.family= gaussian(link= “identity”), with both sexes analyzed separately.

The connection between assortative mating and infection status was tested by evaluating prevalence in males when attached to an infected vs an uninfected female. Population was considered as a random effect to account for prevalence differences among populations, and the ID of the mating pair was nested in the population in order to pair the analysis. The full model to test for assortative mating was as follows: cbind(Nbr_inf_M,Nbr_uninf_M) ~ Infection_Female + (1|Pop/ID_mating), family= binomial), rand.family= gaussian (link= identity), where “cbind(Nbr_inf_M,Nbr_uninf_M)” accounts for the number of infected males given the total number of males in the mating.

## Results

### Mating formation

Most of these results were obtained from studies in natural rockpool populations of *Daphnia magna*. During the summers when we sampled, the average proportion of males was around 30 %, ranging from 5 to 60 % across populations (Figure 1A). The shallow rockpools of this metapopulation allowed us to capture pairs of mating *Daphnia* and observe in glass vials the pair’s separation, the laying of eggs, and the release of the resting egg cases. Most matings involved one male (i.e. monandrous mating), although matings with two males were also frequent (i.e. polyandrous mating, Figure 1B). In rare cases (seven times out of the 968 matings in the study), we found three males in the same mating. Because our sampling our not totally random, our design did not allow us to estimate the frequency of polyandrous mating per population and consequently, we could not determine what parameters may influence it.

**Figure 1:**
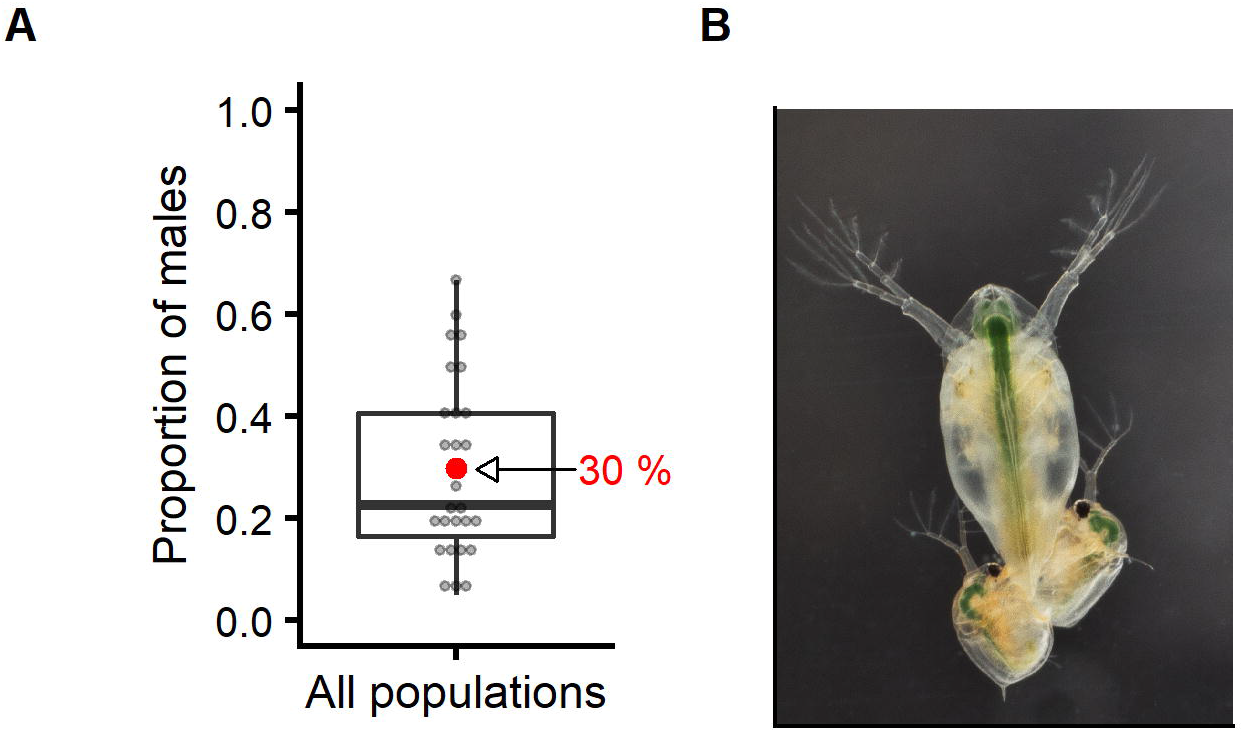
Mating in *Daphnia magna*. A/ Proportion of adult males in *Daphnia magna* rockpool populations. Based on random sampling, the proportion of males ranged from 5 to 60 % with an average about 30 %. Boxplot divides the dataset into quartiles, representing the minimum and maximum, as well as the first quartile (25 % of the dataset lies below it), the median and the third quartile (75 % of the dataset lies below it). Each grey dot represents one population, with the red dot showing the arithmetic mean. B/ Photograph of a female (large individual) mating with two males (small individuals). Females can mate with one (most common), two, or even three males simultaneously.

In 80 % of the cases (382/477), we found that mating females showed the typical morphological changes of the brood pouch associated with the formation of a resting egg case (ephippium), suggesting that they were ready to mate. Since relatively few adult females in a population are typically in this transient stage, this finding indicates that mating pairs do not form randomly. However, since *D. magna* males seem to search for their mates randomly—as opposed to males of the cladoceran *Moina brachiate* that apparently detect the reproductive status of the females (Forró 1997)—it is therefore likely that, in *D. magna* as in *D. pulicaria* (Brewer 1998), females make the choice to accept or escape a mating attempt, and that they are more likely to accept if they are in the appropriate stage of the sexual process. The other 20 % of matings were with non-reproductive females (23/477) or with females reproducing asexually (73/477). If females who are not in sexual reproduction stage try to escape males, these matings may represent cases where males forced matings, not realizing they would not lead to fertilization success. In the 17 matings with asexually reproducing females for which we recorded mating duration, data, 13 of the males stayed on the females for over 10 minutes (i.e. less than when copulating with appropriate females but still long enough to be considered as more than a simple contact), suggesting that they did not realize that the females were asexual.

#### Role of body and spina length

To test if formation of mating pairs is mediated by body length, we compared the body length of mating males to the average body length of randomly caught males and females in the population (Figure 2A, supplementary material 1 – section 1.3.2). Mating females were on average 9.5 % larger than those randomly caught in the population, and mating males were on average 2.3 % larger than those randomly caught in the population. For females, this means that older females produce resting eggs. For males, this suggests that larger males are more successful in securing mating. We also found assortative mating regarding body length: larger than average males paired with larger than average females. The strength of this homogamy (15 %, as described by the estimate of the Pearson correlation) is lower than the average strength for size-related homogamy across animal taxa (31 % according to Jiang *et al.* (2013)) and depends on the population (See supplementary material 1 – section 1.3.3). As larger males may have better access to females, the strength of the homogamy could potentially be lowered by large males also catching small females. Controlling for the average body length of mating males in each population, we also tested whether, males in polyandrous matings tended to be smaller than a single male in a monandrous mating. Males in polyandrous mating were on average10 μm smaller than males in monandrous mating, a tiny and not significant difference (Figure 2B). However, polyandrous matings include males that arrive and attach first to the female, and so could be expected to be about the same length as males in monandrous matings. We thus tested and found that males from polyandrous matings differed in body length (Figure 2C), with the males departing second being on average 1.3% larger than the first males to detach. This suggests that a larger body length could help the male remain longer on the female, potentially giving it an advantage in competition for egg fertilization. We further tested whether the length of the tail spine (spina) might influence access to females, specifically whether individuals with longer spines mated more frequently. To do so, we used relative spine length and subtracted the mean value of individuals of the same sex caught randomly in the population. We found that relative spine length was generally shorter for mating individuals (Supplementary material 1 – section 1.3.6). Altogether, these results suggest that body length plays a role in *D. magna* sexual selection in the rockpool metapopulation.

**Figure 2:**
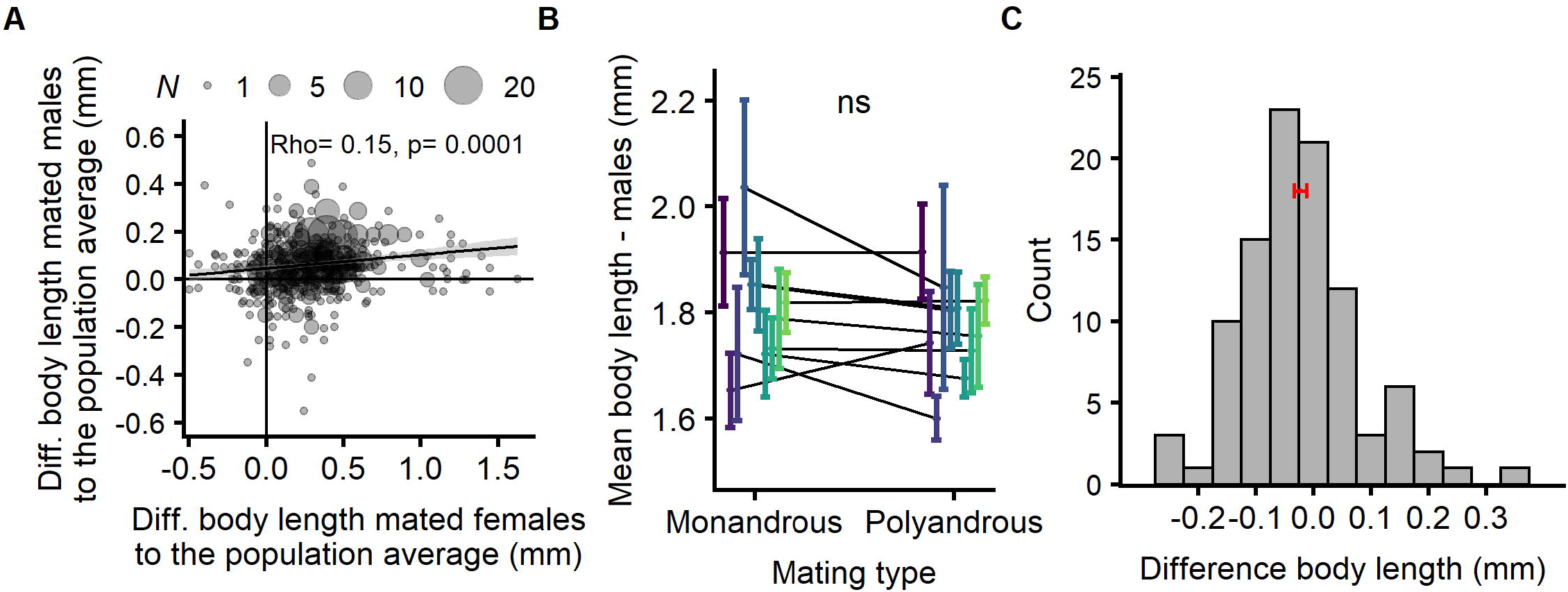
A/ Role of body length in the sexual process. Mating females were 9.5 % larger than randomly caught females in their population. Mating males were on average 2.3 % larger than randomly caught males in their population. Populations showed positive assortative mating for body length: males larger than the average of the population mated with females larger than the average of the population. B/ Body length and mating types. Males in polyandrous mating were 0.01 mm smaller on average than males in monandrous mating, a difference that was not statistically significant. C/ Body length difference of males in polyandrous matings. Males in the same mating are more different than expected by chance (Paired t-test: df=97, t= −0.02, p= 0.03), with the longer-remaining males being, on average, larger than the first males to detach (mean of the differences= 0.02 mm or 1.3 %), suggesting that body length contributes to a competitive advantage in egg fertilization. The red line displays zero difference.

#### Inbreeding avoidance

The fact that females can potentially choose a particular male offers up the possibility that the species can avoid inbreeding (Duthie and Reid 2016). As inbreeding depression and heterosis have been documented in Da*phnia magna*, the ability to avoid mating with relatives provides a selective advantage (De Meester 1993; Ebert et al. 2002; Haag et al. 2002). We investigated this inbreeding question in two natural populations by sequencing four polymorphic microsatellites loci (see details in tables in supplementary material 1 – section 4) and testing to see if females mated with more distantly related males less often than chance would expect. The individuals were captured either in the process of mating (Pop SP1-5: 85 females – 147 males; Pop SP1-6: 92 females – 138 males) or single (16 males for each population). Our result suggests that individuals that formed mating pairs were not less related than random mating simulated in *silico (W*ilcoxon test: Pop SP1-5: W=17085, p=0.6; Pop SP1-6; W=16814, p=0.17. See supplementary material 1 – section 1.4) suggesting, therefore, that inbreeding avoidance is not part of mate formation.

#### Parasite infections

The prevalence of *Hamiltosporidium tvarminnensis*, a common microsporidian that infects this metapopulation, averaged around 40 %, affecting from 0 to 100 % of individuals of both sexes (Supplementary material 1 – section 1.5.1). Thus, *H. tvarminnensis*, a parasite that mediates selection in this metapopulation (Cabalzar et al. 2019), was frequent during our study (Ebert et al. 2001; Lass and Ebert 2006). Laboratory experiments conducted by Roth *et al.* (2008) have shown that infected females produce more sons, implying that the parasite’s prevalence in a population should correlate positively with the number of males relative the number of females. Contrary to this finding, however, *H. tvarminnensis* prevalence in single females in our study did not correlate with the sex-ratio in the population (Figure 3A), suggesting that even if individually-infected females produced more infected sons, the production of males was compensated at the population level, in accordance with Booksmythe *et al.* (2018) and that *D. magna* populations are able to adjust the production of males depending on the current sex-ratio.

**Figure 3:**
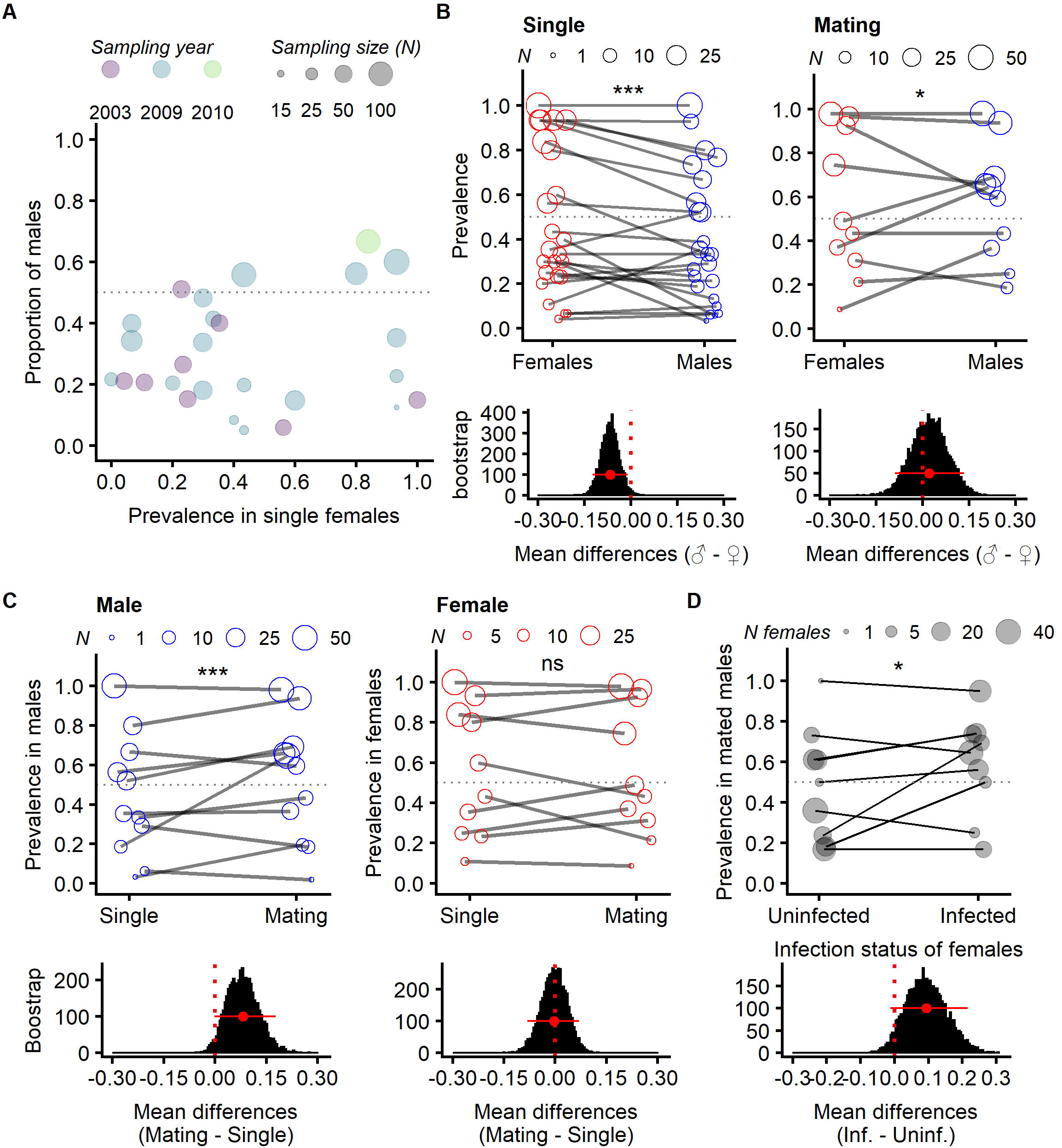
Role of the parasite *H. tvarminnensis* in the sexual process. A/ Correlation between the infection prevalence in single females in the population (i.e., proportion of infected females/total of single females) and in males in the population (df= 1, Chi^2 LRT= 0.064, p= 1). B/ Sexual dimorphism in infection prevalence of single vs mating individuals. Prevalence differences between sexes depended on the mating status (Sex x Mating status: df=1, Chi^2 LRT = 17.84, p= 0.00002). On average, prevalence was higher in females than in males in the populations (left panel, df=1, Chi^2 LRT = 17.84, p= 0.00012, odds ratio: 0.64), but this tendency was reversed in mating (right panel, df=1, Chi^2 LRT = 4.7, p= 0.03, odds ratio: 1.4). C/ Prevalence in single vs mating individuals. On average, the prevalence in mating males was higher than in single males (left panel, df=1, Chi^2 LRT = 17, p= 4.8e-5, odds ratio: 1.8). This difference was not found in females (right panel, df=1, Chi^2 LRT = 0.49, p= 0.49, odds ratio: 0.89). D/ Assortative mating regarding infection. In mating pairs, males were infected on average 1.8 times more when the female was infected (df=1, Chi^2 LRT = 5.2, p= 0.023, odds ratio: 1.8). Histograms beneath the graphs are an estimation graphic as in Ho *et al.* (2019) using bootstrap to estimate the difference between means and its 95 % confidence interval. Population size (N) is indicated by the size of the circles, which is considered in the statistical analysis. We illustrated the significance of the mixed models when p-value >0.05, with * when <0.5, and *** when <0.0001.

Prevalence differed between sexes, but ultimately depended on population and mating status (binomial glm, interaction Sex x Mating status: χ^2^ LRT = 17.9, df=1, p= 0.00002). On average, prevalence was lower in males than females (Figure 3B left panel, odds ratio: 0.64), as suggested in (Roth et al. 2008), but this was reversed in mating pairs (Figure 3B right panel, odds ratio: 1.41), and this result depended strongly on the population.

Parasitism is thought to be a major factor in sexual selection, either because of its direct cost (i.e. females want to avoid becoming infected), or its direct benefits (i.e. healthy males in an infected population might carry good genes). In our study, mating males were infected more often than single males (Figure 3C left panel, odds ratio: 1.85), whereas mating females were infected about equally as often as single females (Figure 3C right panel, odds ratio= 0.89). This could suggest that females choose infected males, as these males may be particularly strong if they can afford to mate while infected. There was also assortative mating based on infection status: Taking the infection rate of the population and the size of our samples into account, infected males were significantly more likely to mate with infected females than with uninfected ones (Figure 3D, odds ratio: 1.81).

### Mating behavior

After moving mating pairs from the pond to the glass jars, we observed them and recorded the time from the moment of their capture to the moment each male detached (Figure 4). The mean time to detachment was 24 min (± 1.9 se). Assuming that males and females in the process of mating were randomly caught and that mating time is normally distributed, this suggests that the total mating time may be twice as long, i.e. about 50 min. This estimate corroborates with the data we collected on the time until detachment—which ranged from a few to 60 minutes (excluding a unique outlier of 242 min). Polyandrous matings lasted longer than monandrous matings, as the second male remained attached longer, but the first male detached as fast as single mating males. Following male detachment, 96 % of females laid their eggs into the brood pouch (Figure 5A), with 86 % (45/52) of them doing so within 10 minutes (Figure 5B). Out of 107 egg cases produced, 93 % contained two eggs, 5 % had one egg, and 3 % were empty (Figure 5C).

**Figure 4:**
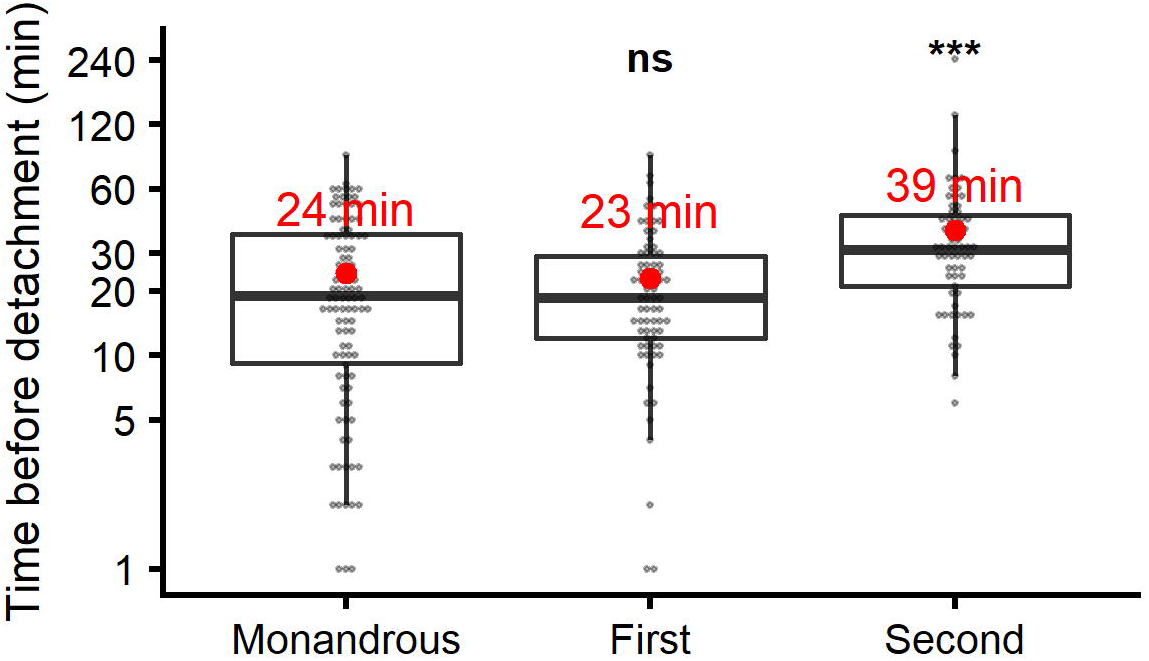
Duration of the mating process. In monandrous mating, males detached on average 24 min after they were caught, similar to the first males in polyandrous mating, which detached on average 23 min after they were caught (Wilcoxon test: W= 3676, p= 0.99). Second males detached on averaged 39 min post capture (Wilcoxon test: W= 4192, p= 0.0002). Those results suggest that mating lasted on average around 50 min. Each dot represents a male in a mating. The y-axis is on a log scale for better illustration.

**Figure 5:**
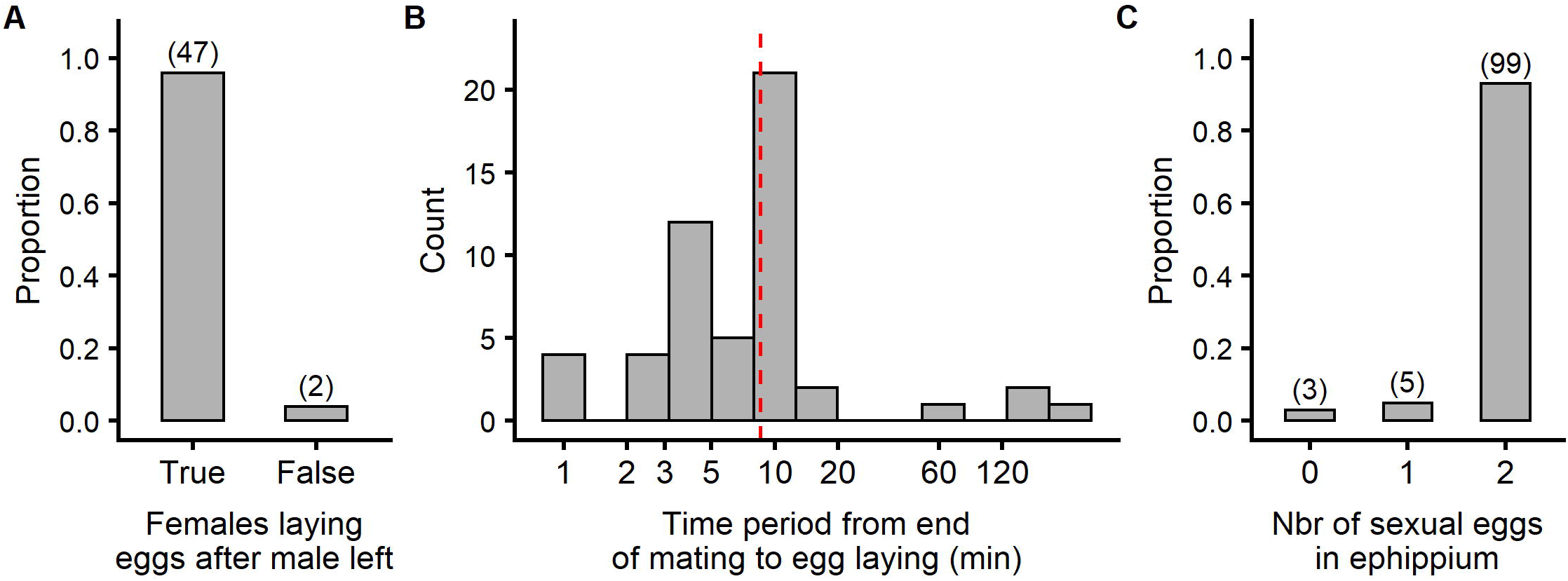
A/ Number of females laying eggs after male departure. B/Time between detachment of the last male and the female depositing sexual eggs; within 10 minutes for 86 % (45/52) of females, and within less than 5 minutes for ~40 % of females. The red line represents the median. C/ Distribution of egg quantities in the ephippium from females caught in the process of mating. Numbers within brackets represent sample sizes.

### Sperm and paternity

#### Fatherhood analysis

The simultaneous presence of two or more males mounted on the same female is not only a strong indicator for direct male–male competition, but also sperm competition. To explore this facet of the mating, we genotyped the mother, the two attached males, and the two embryos from eight egg cases (ephippia), and found that six of the embryo pairs were fertilized by only one of the males (full-sibs), while the other two embryo pairs were half-sibs.

#### Sperm length

We next investigated whether sperm length contributed to male–male competition (Godwin et al. 2017) and whether differences in sperm morphology and/or quality between males of the same mating could be the substrate for selection upon sperm competition. Looking at 46 polyandrous matings from natural populations, we found differing sperm lengths in males from the same mating (Figure 6A). Although it is difficult to tell whether this difference is biologically relevant, more than 50 % of the simultaneously attached male pairs had a mean difference greater than 0.77 μm (i.e. 8.6 % larger than the averaged sperm length). The average sperm length of the second male to detach was 0.96 times the average of the first, a difference that is not statistically significant (df= 1, Chi^2 LRT= 2.9, p= 0. 086).

**Figure 6:**
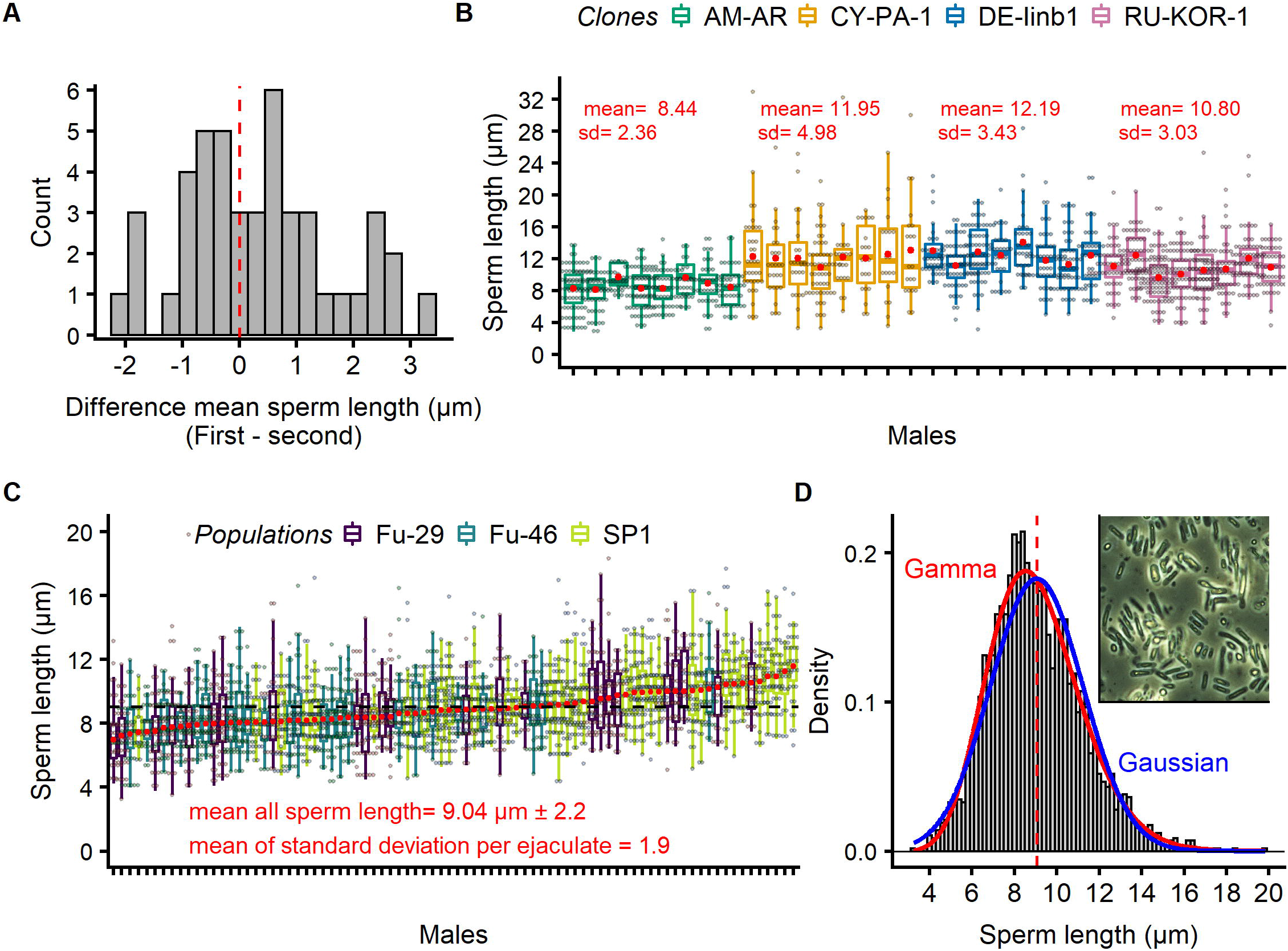
Sperm length of *Daphnia magna*. A/ Difference between mean ejaculates from males in the same mating (first male to detach minus the second). Difference in sperm length was larger than 0.77 μm (i.e. 8.6 % larger than the averaged sperm length) in over 50% of the cases; however, the position to detach did not predict the direction of the difference, as the second male to detach was 0.96 of the average of the first, a difference that was not statistically significant (df= 1, Chi^2 LRT= 2.9, p= 0. 086). B/ Sperm length in ejaculate of males of four clones under laboratory conditions revealed more variation between than within clones (df= 3, Chi^2 LRT= 17, p= 6e-4), indicating a genetic component of sperm length. C/ Sperm length in ejaculates of males captured from three natural populations while mating ranked by median length within their ejaculate. Average standard deviation within an ejaculate was 1.9 μm, about as large as the standard deviation of all the measured sperm, 2.2 μm. D/ Distribution of sperm length in all males represented in C. Distributions within ejaculates were generally better described by a Gamma than by a Gaussian distribution (or by a mixture of two Gaussian distributions), excluding the hypothesis of two different morphs, with possibly different functions. The inlet photograph shows sperm length variation in a typical ejaculate.

To test for genetic variation in sperm length, we studied the ejaculates of four laboratory-raised *D. magna* clones and found more variation in sperm length between clones than between individuals from the same genetic background; we also found that some clones had a higher mean sperm length than others (Figure 6B). Considering single ejaculates of males from the rockpools, we found the variation in sperm length within ejaculate to be strikingly large (Figure 6C inlet), ranging from 3 to 20 μm with an average of 9 μm (Figure 6C). The average standard deviation in sperm length for an ejaculate (i.e. 1.9 μm) was close to the standard deviation calculated for ejaculates across males from three different populations (i.e. 2.2 μm). As large variation in sperm length is often attributed to different sperm subpopulations, we looked at the distribution in sperm length within each ejaculate (not shown in Figure 6D, which only illustrates the distribution of the pooled sperm length to provide a general idea of sperm length variation) and found that it was better described by a Gamma distribution (Combined AIC per model for each ejaculate was 12096.67) than a Gaussian distribution (Combined AIC per model for each ejaculate was 12218.56) or by a mixture of two Gaussian distributions (Combined AIC per model for each ejaculate was 12613.04). Thus, it is less parsimonious to suggest that ejaculates are composed of a mixture of two morphologies, each with different roles in sperm competition.

## Discussion

Because it is often easier to focus on the asexual part of the life cycle of cyclical parthenogenetic species, the study of the evolution of these species generally concentrates on survival and reproduction (natural selection), without considering the possible role of sexual selection. By working with *D. magna* populations in shallow rockpools, where sexual reproduction is rather frequent, we were able to focus on the biology of this crustacean’s sexual reproduction in a natural setting and to gain an understanding of the role of sexual selection. We were able to conclude that clear evidence exists for sexual selection in cyclical parthenogenetic species. Below we discuss how sexual selection unfolds in the sequential steps of the sexual reproduction process, pointing out the possible mechanisms at work in each step. The entire process is summarized in Table 1.

**Table 1:**
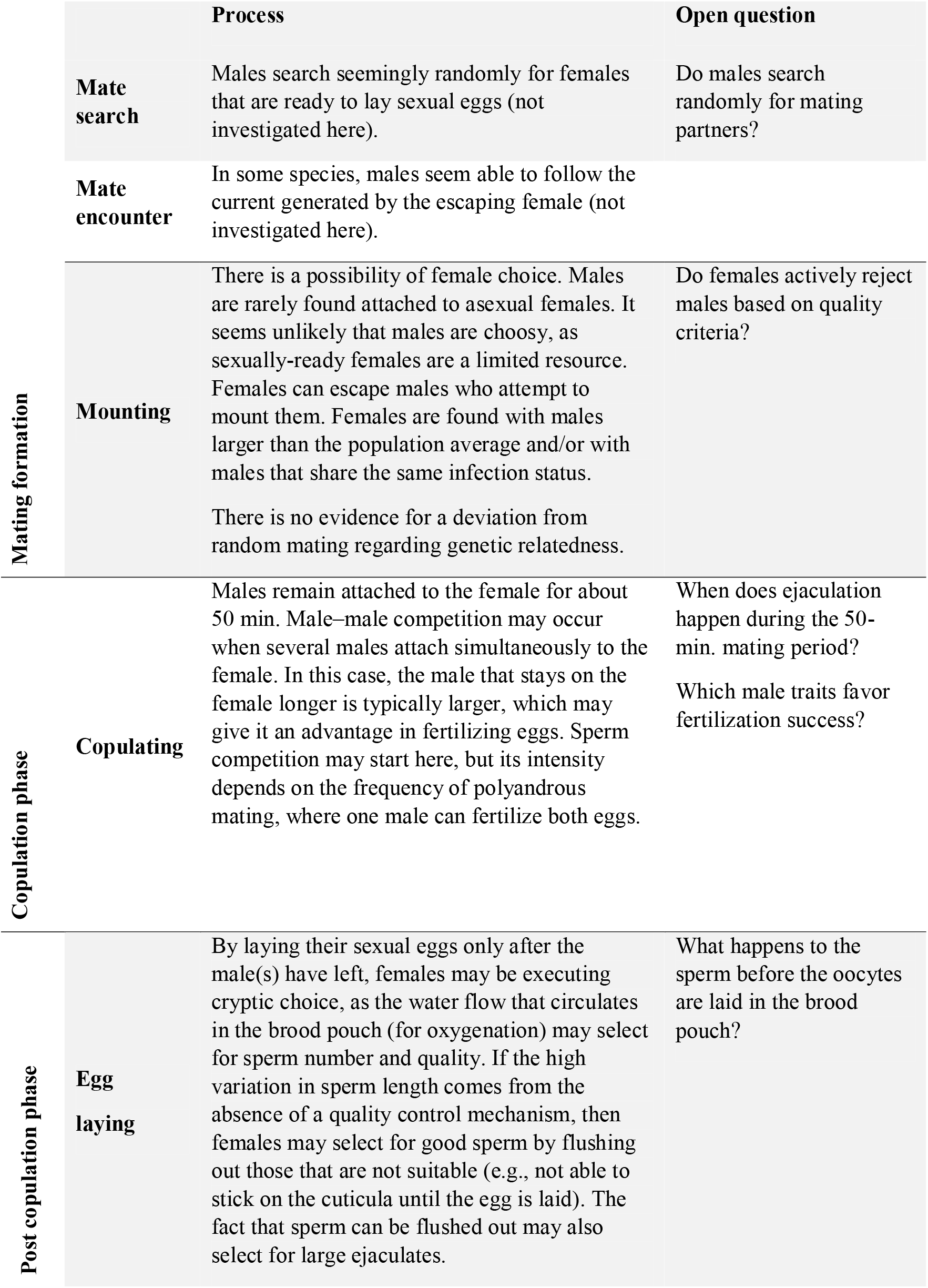
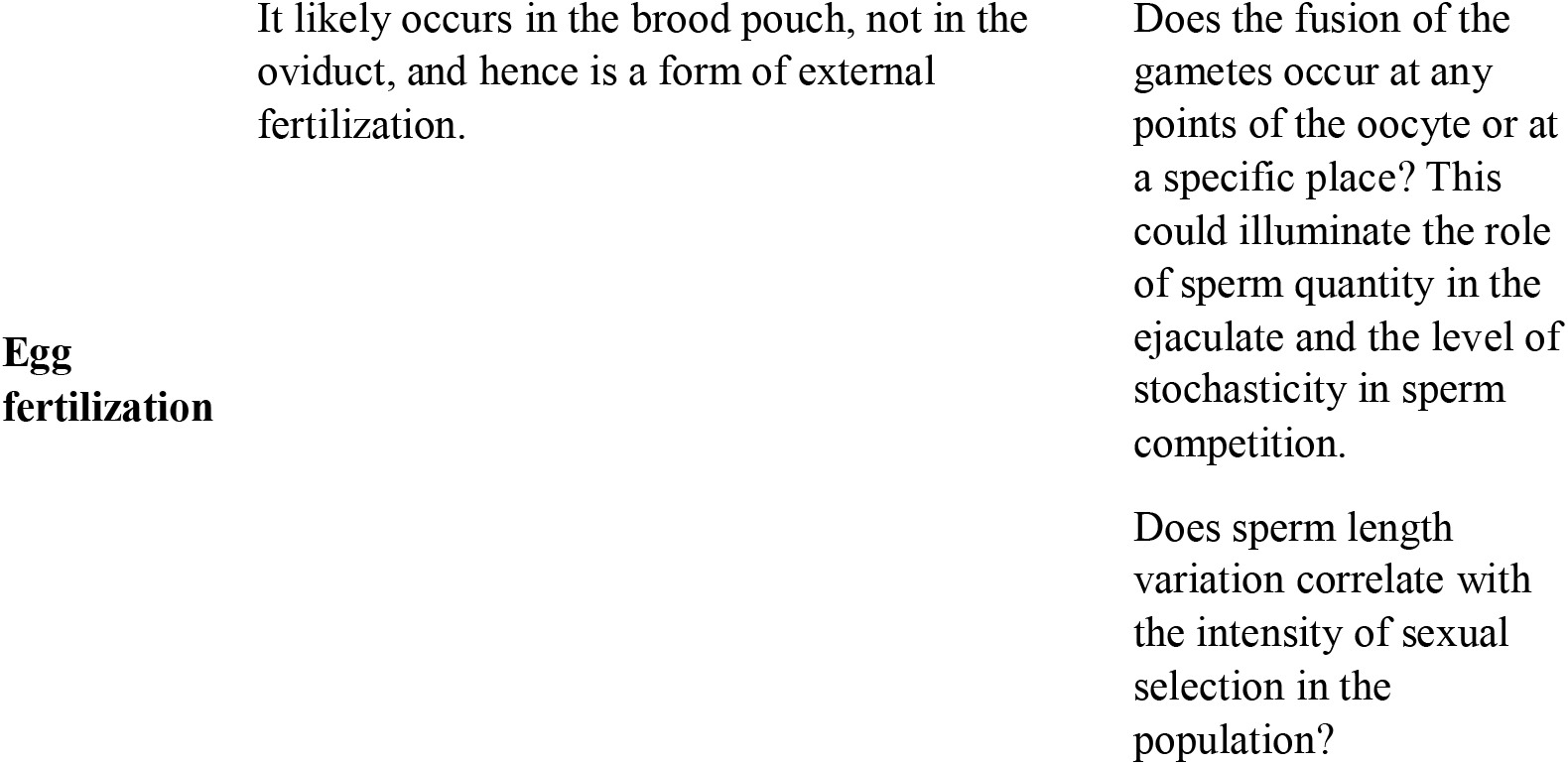
Description of the sexual process in *Daphnia magna*.

### Mating formation

As with other cyclical parthenogenetic species (Dixon 1977; Ward et al. 1984; Snell and Hoff 1985; Hand 1991), environmental conditions trigger some female *Daphnia* to switch from producing asexual daughters to producing asexual sons (Ebert 2005). Other females switch from producing asexual daughters to producing sexual eggs that need male fertilization. Males search for appropriate females by swimming quickly in pursuit. Although, male–female encounters appear to be random, some form of sorting must take place, as mating males are primarily found on females in the late phase of the sexual process, when the structure of the egg case is already visible and oocyte release is imminent. Furthermore, we found evidence of assortative mating for body length and infection status, suggesting that, after an initial, presumably random encounter, males either abandon females that do not meet their expectations, or females reject males when they are not at the right stage of reproduction, or when the males do not fit their quality assessment. Typically, even if it may be less strong in cyclical parthenogenetic species than in obligate sexual species, there are more males than females ready to mate in a population at any given time, so males are generally not a limited resource for females, whereas females in the right stage of fertility are more likely to be limited for males. Hence, females are more likely to be choosy, while males are more likely to accept any female, they encounter who is in the right stage of fertility. Consistent with this, we frequently observed females violently shaking off males that attempted to mount them in the rockpool populations (D. Duneau, D. Ebert, personal observation).

Our results also showed that males in mating pairs were more frequently infected than single males in the population. This result was unexpected, as infected males may be weaker and have been shown to have reduced sperm counts (Roth et al. 2008). Females may choose infected males because if they are infected and still able to catch a female, they are likely to be strong. In this case, infection would be an honest signal, a handicap (Zahavi 1975). Alternatively, males may counteract the females’ attempts to push them away, with stronger males being more able to resist these rebuffs. This would be consistent with our finding that males in the process of mating are, on average, larger than single males. It is also possible that infected males who are still alive and able to catch females are the strongest males. Since the parasite cannot spread to the female or her offspring during mating, there is no risk associated with mating. Thus, in the mating formation phase of the sexual process, sexual selection, notably for body length, may be driven by female choice.

### Copulation

As the density and proportion of sexual animals in a population increase, so too do multiple encounters, with polyandrous matings becoming more common. We observed many cases of polyandrous matings involving two males hooked to one female, and even some cases with three or four males. These matings open the door for two types of male–male competition: First, males may compete directly with each other for the best position and the longest stay-time on a female. We found that the male who stayed on the female longest was, on average, slightly larger than the male who left earlier, supporting the theory that stronger males have more control over the situation. Second, males may compete via their sperm. If multiple males deposit their ejaculates around the same time into the female’s brood chamber, competition may favor males with more and/or better sperm. Indeed, we found that one male fertilized both eggs in six out of eight cases studied in detail. As *D. magna* sperm seem non-motile, swimming speed in the brood pouch is likely not a factor in the competition.

### Fertilization

We observed the following sequence regarding fertilization: Within ten minutes after a male left, most females released one or two eggs into their brood pouches, which, by this time, had developed into the future egg case (ephippium), specialized as a resting structure. The oviducts open into this brood pouch at its caudal end, close to where the male attaches. Fertilization occurs in either the brood pouch (external fertilization) or in the two oviducts (internal fertilization). Although fertilization has never before been studied in *D. magna*, research on *D. pulex* has suggested that fertilization occurred internally, before deposition in the brood pouch (Ojima 1958; Hobæk and Larsson 1990). Here we argue, on the contrary, that fertilization in *D. magna* is most likely external, and that the most plausible mechanism of fertilization is that the male ejaculates a large number of sperm into the brood pouch, and that the oocytes come into contact with this sperm when they enter the brood pouch.

Although future studies are needed to fully test this hypothesis, several arguments support the brood pouch fertilization hypothesis over the internal fertilization hypothesis. First, the genital papilla on the male’s abdomen is large and conical shaped, and as such, has rather limited access to the female oviducts; it could not insert itself into the oviduct, which is closed until the eggs are laid (Lee et al. 2019). Second, as each sexual egg is released from one oviduct, males could probably only access one of the oviducts—the one closest to the side they are attached to. This would strongly reduce the possibility that both eggs could be fertilized by only one attached male, and, as both eggs typically hatch, they must both be fertilized. Also, we found that in double matings, the two eggs are often fertilized by the same male, which would not be feasible if the male needed to insert its papilla into each ovary. Third, at least from our observation, the sperm is apparently not motile. It is therefore unclear how the sperm could find the oviduct and travel within it. Fourth, the shape of the sperm is not streamlined to move in one direction. It ranges from oval to short rod-shaped with two blunt ends (Figure 6D). These facts, taken together, convince us that fertilization inside the oviduct is unlikely in *D. magna* and that, more likely, the males release their sperm into the brood pouch where they wait for the unfertilized eggs to arrive.

How does the sperm meet the oocyte? The brood pouch is part of the outside environment. It is open to the outside, and water can circulate freely through it. Sperm may cover the inner lining of the brood pouch, so that the eggs touch them as soon as they are released from the ovary. Sperm may also preferentially attach to the area surrounding the opening of the oviduct, so they can fertilize the eggs in the moment of their release, although there is no obvious structure supporting this speculation. If this were the case, the first male to attach would have an advantage, and mating may not need to last for 50 min on average. Another option is that sperm do not attach to any tissue before the eggs arrive but stay suspended in the brood pouch and risk getting flushed out by the stream of water that oxygenates the brood pouch. For fertilization to occur, the female may temporarily halt this water flow until the eggs are laid, so as to avoid washing out all the sperm, or, alternatively, the flushing out may select for males that produce large ejaculates with high quality sperm. Either way, sperm competition could take place in this selection arena, with ejaculate quality, sperm quantity and time of ejaculate deposition being crucial aspects of fertilization success.

Sex-ratio can be highly variable among *D. magna* populations and is known to vary strongly over the season (Booksmythe et al. 2018). Consequently, the intensity of sperm competition may also vary among populations and over time. When males are numerous, they must compete to fertilize the oocytes in polyandrous matings. Those with the highest number of sperms remaining in the brood pouch after they depart and the female lays her eggs are likely to have the highest fertilization success. This would favor males who produce more sperm and who stay longer on the females. When polyandrous mating is less common, it is still likely that ejaculate may evolve for higher sperm counts, as sperm released in the female brood pouch can still be flushed out. In this case, the reason for the selection would be sperm limitation (as defined in Liao *et al.*, 2018), not sperm competition. In either case, the difference in optima for sperm count upon selection by sperm limitation or sperm competition is unknown.

As most sperm are produced early in a male’s life (Wuerz et al. 2017), the total sperm count is limited by the size of the spermiduct. In *Daphnia*, vacuoles compact the mature sperm in an extracellular process before they are released into the spermiduct to maximize the amount of stored sperm (Wingstrand 1978). Even though the sperm we observed in *Daphnia* medium had remained intact, it is expected that the vacuole opens in the brood pouch, eventually upon contact with the oocyte. Hence, sperm number is tightly correlated with sperm length, or at least with its compaction. When sperm count is under strong selection, directional selection is expected to select for an optimal length versus optimal quantity. However, if it is costly to precisely control sperm length, the variation in sperm length is predicted to correlate negatively with the intensity of sperm competition (Bauer and Breed 2006; Fitzpatrick and Baer 2011; Varea-Sanchez et al. 2014; Rowley et al. 2019). The high variance in sperm length we observed here could, therefore, indicate weak sperm competition overall. In such a case, the rod-like sperm cell does not require a strict morphology, so its shape can be more or less condensed.

### Sexual selection gradient in each sex

A. J. Bateman articulated several principles explaining the undiscriminating drive of males to obtain mates versus the choosiness of females (Bateman 1948). In *Daphnia*, the relationship between mating success and reproductive success in females is what he calls single-mate saturation (Bateman 1948). This means that a single mating suffices to fertilize the entire clutch, so the female has no increase in reproductive success once she obtains a single mating partner. Whether or not females benefit from the male–male competition of polyandrous mating is not clear. Their clutch size is unaffected; only the paternity is potentially divided between the inseminating males. The sexual selection gradient in female *Daphnia* should therefore be weak (See figure 2 in Arnold, 1994). In males, however, the expected reproductive success increases linearly with the number of matings in the absence of sperm competition. For males, when polyandrous mating is common, the relationship between mating success and reproductive success is what Bateman calls diminishing returns (Bateman 1948). Here, as ejaculate size diminishes with the number of matings, the males’ competitiveness mechanically decreases too, as does the number of potential offspring gained with each additional mate (Arnold 1994). Hence, male mating success correlates linearly with reproductive success when males are proportionally less numerous than fertile females, while the correlation saturates in presence of male-male competition.

#### Sexual selection in other cyclical parthenogenetic species

Although only little is known about sexual selection in other cyclical parthenogenetic species, there are reasons to believe that it plays an equally important role in these species as it does in *Daphnia*. In aphids, for example, the best-studied cyclical parthenogenetic species, we know that some aphid species can recognize specific mates (Guldemond et al. 1994). Also, females have been shown to release species-specific sex pheromones daily and at specific times to call males (Guldemond and Dixon 1994). These two mechanisms are used to reduce interspecific insemination and may have played a role in speciation by reinforcement in aphids (Guldemond and Dixon 1994). They could also be used by females to choose mates within the same species. Although it is known that aphids can mate for more than once and display some form of pre-copulatory stroking behavior, the occurrence of sperm competition seems unknown (Doherty and Hales 2002). Female choice is certainly possible, as females can avoid inbreeding and refuse to mate with certain males (Huang and Caillaud 2012). Together, these traits in aphids along with the example of *Daphnia* suggest that sexual selection probably plays a role in the evolution of cyclical parthenogenetic species. Since sexual reproduction is generally associated with the capacity to respond to environmental changes or dispersal, this role may be more important than previously thought.

## Conclusion

Cyclical parthenogenesis, a reproductive strategy where organisms go through several rounds of clonal reproduction before reproducing sexually, is widespread in many taxa, including crustaceans, rotifers, aphids, and in human parasitic nematodes. While often understudied in these species, sexual selection is an important form of selection even when it is not the dominant reproductive strategy. Because the sexual event occurs only after several generations of clonal reproduction, the intensity of sexual selection on sexual traits is overall reduced and the temporary absence of sexual reproduction in cyclical parthenogens may give a non-negligible role to drift in the evolution of sexually dimorphic traits. The reduced intensity of sexual selection may allow genetic variation to accumulate and thus, enable selection to occur on multiple alleles at once. This principle is reminiscent of the first two phases of Wright’s shifting-balance theory (Wright 1982). Furthermore, it is likely that during clonal reproduction, sexual traits (e.g. body size) may deviate from their optimal for mating/fertilization success because they have different optimum under natural selection.

## Acknowledgements

We thank Michelle Krebs and Urs Stiefel for help in the laboratory and Juergen Hottinger, Pepijn Luickjx, Marjo Saastamoinen, Patrícia Beldade, Karen Haag, Andrea Hoffman, Gleb Georg Ebert and Katharina Ida Ebert for help in the field. We also thank the staff at Tvärminne Zoological Station for logistic support, as well as Axel Wiberg, Jeremias Brand, Vitor Faria and Eric Dexter for helpful discussions. The study was supported by the Swiss National Science Foundation. DD was supported by the French Laboratory of Excellence project ‘TULIP’ (ANR-10-LABX-41; ANR-11-IDEX-0002– 02).

## References

Altermatt, F., and D. Ebert. 2010. Populations in small, ephemeral habitat patches may drive dynamics in a *Daphnia magna* metapopulation. Ecology 91:2975–2982.

Andersson, M. 1994. Sexual selection. Princeton University Press, Princeton, NJ, USA.

Arnold, S. J. 1994. Bateman’s principles and the measurement of sexual selection in plants and animals. The American Naturalist 144:S126–S149.

Bateman, A. 1948. Intra-sexual selection in *Drosophila*. Heredity 2:349–368.

Bauer, M., and W. G. Breed. 2006. Variation of sperm head shape and tail length in a species of Australian hydromyine rodent: The spinifex hopping mouse, *Notomys alexis*. Reproduction, Fertility and Development 18:797–805.

Booksmythe, I., N. Gerber, D. Ebert, and H. Kokko. 2018. *Daphnia* females adjust sex allocation in response to current sex ratio and density. Ecology Letters 21:629–637.

Brewer, M. C. 1998. Mating behaviours of *Daphnia pulicaria*, a cyclic parthenogenD: comparisons with copepods. Philosophical Transactions of the Royal Society of London. Series B: Biological Sciences 353:805–815.

Cabalzar, A. P., P. D. Fields, Y. Kato, H. Watanabe, and D. Ebert. 2019. Parasite-mediated selection in a natural metapopulation of *Daphnia magna*. Molecular Ecology 28:4770–4785.

Carter, J. L., D. E. Schindler, and T. B. Francis. 2017. Effects of climate change on zooplankton community interactions in an Alaskan lake. Climate Change Responses 4:1–12.

Cavalheri, H. B., C. C. Symons, M. Schulhof, N. T. Jones, and J. B. Shurin. 2019. Rapid evolution of thermal plasticity in mountain lake *Daphnia* populations. Oikos 128:692–700.

Clutton-Brock, T. 2017. Reproductive competition and sexual selection. Philosophical Transactions of the Royal Society B: Biological Sciences 372:20160310.

Cornetti, L., P. D. Fields, K. Van Damme, and D. Ebert. 2019. A fossil-calibrated phylogenomic analysis of *Daphnia* and the Daphniidae. Molecular Phylogenetics and Evolution 137:250–262.

Cox, R. M., and R. Calsbeek. 2010. Sex-specific selection and intraspecific variation in sexual size dimorphism. Evolution 64:798–809.

Crease, T. J., and P. D. N. Hebert. 1983. A test for the production of sexual pheromones by *Daphnia magna* (Crustacea: Cladocera). Freshwater Biology 13:491–496.

Darwin, C. 1871. The descent of man, and selection in relation to sex. J. Murray, London.

De Meester, L. 1993. Inbreeding and outbreeding depression in *Daphnia*. Oecologia 96:80–84.

Decaestecker, E., S. Gaba, J. a M. Raeymaekers, R. Stoks, L. Van Kerckhoven, D. Ebert, and L. De Meester. 2007. Host-parasite “Red Queen” dynamics archived in pond sediment. Nature 450:870–873.

Delignette-Muller, M. L., and C. Dutang. 2015. *fitdistrplus*: An R package for fitting distributions. Journal of Statistical Software 64:1–34.

Dixon, A. F. G. 1977. Aphid ecology: Life cycles, polymorphism, and population regulation. Annual Review of Ecology and Systematics 8:329–353.

Doherty, H. M., and D. F. Hales. 2002. Mating success and mating behaviour of the aphid, *Myzus persicae* (Hemiptera: Aphididae). European Journal of Entomology 99:23–27.

Duneau, D., P. Luijckx, L. F. Ruder, and D. Ebert. 2012. Sex-specific effects of a parasite evolving in a female-biased host population. BMC Biology 10:104.

Duthie, A. B., and J. M. Reid. 2016. Evolution of inbreeding avoidance and inbreeding preference through mate choice among interacting relatives. The American Naturalist 188:651–667.

Ebert, D. 2005. Ecology, epidemiology, and evolution of parasitism in *Daphnia* [internet]. National library of medicine (US), national center for biotechnology information, Bethesda (MD).

Ebert, D., D. Duneau, M. D. Hall, P. Luijckx, J. P. Andras, L. Du Pasquier, and F. Ben-Ami. 2016. A population biology perspective on the stepwise infection process of the bacterial pathogen *Pasteuria ramosa* in *Daphnia*. Advances in Parasitology 91:265–310.

Ebert, D., C. Haag, M. Kirkpatrick, M. Riek, J. W. J. W. Hottinger, and V. I. Pajunen. 2002. A selective advantage to immigrant genes in a *Daphnia* metapopulation. Science 295:485–488.

Ebert, D., J. W. Hottinger, and V. I. Pajunen. 2001. Temporal and spatial dynamics of parasite richness in a *Daphnia* metapopulation. Ecology 82:3417–3434.

Fitzpatrick, J. L., and B. Baer. 2011. Polyandry reduces sperm length variation in social insects. Evolution 65:3006–3012.

Forró, L. 1997. Mating behaviour in *Moina brachiata* (Jurine, 1820) (Crustacea, Anomopoda). Hydrobiologia 360:153–159.

Garnier, S. 2018. *viridis*: Default color maps from “matplotlib.”

George, D. G., D. P. Hewitt, J. W. G. Lund, and W. J. P. Smyly. 1990. The relative effects of enrichment and climate change on the long-term dynamics of *Daphnia* in Esthwaite Water, Cumbria. Freshwater Biology 23:55–70.

Godwin, J. L., R. Vasudeva, Ł. Michalczyk, O. Y. Martin, A. J. Lumley, T. Chapman, and M. J. G. Gage. 2017. Experimental evolution reveals that sperm competition intensity selects for longer, more costly sperm. Evolution Letters 1:102–113.

Goren, L., and F. Ben-Ami. 2013. Ecological correlates between cladocerans and their endoparasites from permanent and rain pools: Patterns in community composition and diversity. Hydrobiologia 701:13–23.

Guldemond, J. A., and A. F. G. Dixon. 1994. Specificity and daily cycle of release of sex pheromones in aphids: a case of reinforcement? Biological Journal of the Linnean Society 52:287–303.

Guldemond, J. A., A. F. G. Dixon, and W. T. Tigges. 1994. Mate recognition in *Cryptomyzus* aphids: copulation and insemination. Entomologia Experimentalis et Applicata 73:67–75.

Haag, C. R., J. W. Hottinger, W. rgen, M. Riek, D. Ebert, M. Riex, and D. Ebert. 2002. Strong inbreeding depression in a *Daphnia* metapopulation. Evolution 56:518–526.

Haag, K. L., J. I. R. Larsson, D. Refardt, and D. Ebert. 2011. Cytological and molecular description of *Hamiltosporidium tvaerminnensis gen. et sp. nov.*, a microsporidian parasite of *Daphnia magna*, and establishment of *Hamiltosporidium magnivora comb. nov*. Parasitology 138:447–462.

Hahn, M. A., C. Effertz, L. Bigler, and E. von Elert. 2019. 5α-cyprinol sulfate, a bile salt from fish, induces diel vertical migration in *Daphnia*. eLife 8.

Hand, S. C. 1991. Metabolic dormancy in aquatic invertebrates. Pages 1–50 in Advances in Comparative and Environmental Physiology: Volume 8. Springer Berlin Heidelberg, Berlin, Heidelberg.

Hebert, P. D. N., and R. D. Ward. 1972. Inheritance during parthenogenesis in *Daphnia magna*. Genetics 71:639–642.

Ho, J., T. Tumkaya, S. Aryal, H. Choi, and A. Claridge-Chang. 2019. Moving beyond P values: data analysis with estimation graphics. Nature Methods 16:565–566.

Hobæk, A., and P. Larsson. 1990. Sex determination in *Daphnia magna*. Ecology 71:2255–2268.

Houslay, T. M., K. F. Houslay, J. Rapkin, J. Hunt, and L. F. Bussière. 2017. Mating opportunities and energetic constraints drive variation in age-dependent sexual signalling. Functional Ecology 31:728–741.

Huang, M. H., and M. C. Caillaud. 2012. Inbreeding avoidance by recognition of close kin in the pea aphid, *Acyrthosiphon pisum*. Journal of Insect Science 12:1–13.

Jansen, M., A. Coors, R. Stoks, and L. De Meester. 2011. Evolutionary ecotoxicology of pesticide resistance: A case study in *Daphnia*. Ecotoxicology 20:543–551.

Jiang, Y., D. I. Bolnick, and M. Kirkpatrick. 2013. Assortative mating in animals. The American Naturalist 181:E125–E138.

Kaldun, B., and O. Otti. 2016. Condition-dependent ejaculate production affects male mating behavior in the common bedbug *Cimex lectularius*. Ecology and Evolution 6:2548–2558.

Lampert, K. P. 2009. Facultative parthenogenesis in vertebrates: Reproductive error or chance? Sexual Development 2:290–301.

Lass, S., and D. Ebert. 2006. Apparent seasonality of parasite dynamics: Analysis of cyclic prevalence patterns. Proceedings of the Royal Society B: Biological Sciences 273:199–206.

Law, C. J., and R. S. Mehta. 2018. Carnivory maintains cranial dimorphism between males and females: Evidence for niche divergence in extant Musteloidea. Evolution 72:1950–1961.

Lee, D., J. S. Nah, J. Yoon, W. Kim, and K. Rhee. 2019. Live observation of the oviposition process in *Daphnia magna*. PLoS ONE 14:1–9.

Li, C. C., B. E. Weeks, and A. Chakravarti. 1993. Similarity of DNA fingerprints due to chance and relatedness. Human Heredity 43:45–52.

Liao, W. B., Y. Huang, Y. Zeng, M. J. Zhong, Y. Luo, and S. Lüpold. 2018. Ejaculate evolution in external fertilizers: Influenced by sperm competition or sperm limitation? Evolution 72:4–17.

Lonsdale, D. J., M. A. Frey, and T. W. Snell. 1998. The role of chemical signals in copepod reproduction. Journal of Marine Systems 15:1–12.

Lumley, A. J., Ł. Michalczyk, J. J. N. Kitson, L. G. Spurgin, C. A. Morrison, J. L. Godwin, M. E. Dickinson, et al. 2015. Sexual selection protects against extinction. Nature 522:470–473.

Lynch, M. 1988. Estimation of relatedness by DNA fingerprinting. Molecular Biology and Evolution 5:584–599.

Lynch, M., and K. Ritland. 1999. Estimation of pairwise relatedness with molecular markers. Genetics 152:1753–1766.

Metschnikoff, É. 1884. Über eine Sprosspilzkrank heit der Daphnien. Beitrag zur Lehre iiber den Kampf der E hagocyten gegen Krankheitserreger. Virchows Archiv 96:177–195.

Milligan, B. G. 2003. Maximum-likelihood estimation of relatedness. Genetics 163:1153–1167.

Morehouse, N. I. 2014. Condition-dependent ornaments, life histories, and the evolving architecture of resource-use. Integrative and Comparative Biology 54:591–600.

Ojima, Y. A. 1958. A cytological study on the development and maturation of the parthenogenetic and sexual eggs of *Daphnia pulex* (Crustacea–Cladocera). Kwansei Gakuen Univ Ann Stud. 6:123–176.

Orlansky, S., and F. Ben-Ami. 2019. Genetic resistance and specificity in sister taxa of *Daphnia*: Insights from the range of host susceptibilities. Parasites and Vectors 12:1–10.

Pajunen, V. I. 1986. Distributional dynamics of *Daphnia* species in a rock-pool environment. Annales Zoologici Fennici 23:131–140.

Pew, J., P. H. Muir, J. Wang, and T. R. Frasier. 2015. Related: An R package for analysing pairwise relatedness from codominant molecular markers. Molecular Ecology Resources 15:557–561.

Queller, D. C., and K. F. Goodnight. 1989. Estimating relatedness using genetic markers. Evolution 43:258–275.

R Core Team. 2019. R: A language and environment for statistical computing. Vienna, Austria.

Reger, J., M. I. Lind, M. R. Robinson, and A. P. Beckerman. 2018. Predation drives local adaptation of phenotypic plasticity. Nature Ecology and Evolution 2:100–107.

Ritland, K. 1996. Estimators for pairwise relatedness and individual inbreeding coefficients. Genetical Research 67:175–185.

Roth, O., D. Ebert, D. B. Vizoso, A. Bieger, S. Lass, B. Annette, and S. Lass. 2008. Male-biased sex-ratio distortion caused by *Octosporea bayeri*, a vertically and horizontally-transmitted parasite of *Daphnia magna*. International Journal for Parasitology 38:969–979.

Roulin, A. C., J. Routtu, M. D. Hall, T. Janicke, I. Colson, C. R. Haag, and D. Ebert. 2013. Local adaptation of sex induction in a facultative sexual crustacean: insights from QTL mapping and natural populations of *Daphnia magna*. Molecular Ecology 22:3567–3579.

Rousset, F., and J.-B. Ferdy. 2014. Testing environmental and genetic effects in the presence of spatial autocorrelation. Ecography 37:781–790.

Rowley, A., L. Locatello, A. Kahrl, M. Rego, A. Boussard, E. Garza-Gisholt, R. M. Kempster, et al. 2019. Sexual selection and the evolution of sperm morphology in sharks. Journal of Evolutionary Biology 32:1027–1035.

RStudio Team. 2016. RStudio: Integrated Development Environment for R. Boston, MA.

Scrucca, L., M. Fop, B. Murphy, T., and E. Raftery, Adrian. 2016. mclust 5: Clustering, classification and density estimation using Gaussian finite mixture models. The R Journal 8:289–317.

Sheikh-Jabbari, E., M. D. Hall, F. Ben-Ami, and D. Ebert. 2014. The expression of virulence for a mixed-mode transmitted parasite in a diapausing host. Parasitology 141:1097–1107.

Shine, R. 1979. Sexual selection and sexual dimorphism in the Amphibia. Copeia 1979:297–306.

Shuker, D. M. 2010. Sexual selection: endless forms or tangled bank? Animal Behaviour 79:e11–e17.

Snell, T. W., and F. H. Hoff. 1985. The effect of environmental factors on resting egg production in the rotifer *Brachionus plicatilis*. Journal of the World Mariculture Society 16:484–497.

Springer, M. S., W. J. Murphy, E. Eizirik, and S. J. O’Brien. 2003. Placental mammal diversification and the Cretaceous-Tertiary boundary. Proceedings of the National Academy of Sciences of the United States of America 100:1056–1061.

Tollrian, R., and C. Heibl. 2004. Phenotypic plasticity in pigmentation in *Daphnia* induced by UV radiation and fish kairomones. Functional Ecology 18:497–502.

Urca, H., and F. Ben-Ami. 2018. The role of spore morphology in horizontal transmission of a microsporidium of *Daphnia*. Parasitology 145:1452–1457.

Varea-Sanchez, M., L. Gomez Montoto, M. Tourmente, and E. R. S. Roldan. 2014. Postcopulatory sexual selection results in spermatozoa with more uniform head and flagellum sizes in rodents. PLoS One 9:e108148.

Vizoso, D. B., and D. Ebert. 2005. Mixed inoculations of a microsporidian parasite with horizontal and vertical infections. Oecologia 143:157–166.

Walsh, P. S., D. A. Metzger, and R. Higuchi. 1991. Chelex 100 as a medium for simple extraction of DNA for PCR-based typing from forensic material. BioTechniques 10:506–513.

Wang, J. 2002. An estimator for pairwise relatedness using molecular markers. Genetics 160:1203–1215.

Wang, J.. 2011. Coancestry: A program for simulating, estimating and analysing relatedness and inbreeding coefficients. Molecular Ecology Resources 11:141–145.

Ward, S. A., S. R. Leather, and A. F. G. Dixon. 1984. Temperature prediction and the timing of sex in aphids. Oecologia 62:230–233.

Weismann, A. 1893. The germ-plasm: A theory of heredity. Charles Scribner’s Sons, New York.

Whitlock, M. C., and A. F. Agrawal. 2009. Purging the genome with sexual selection: Reducing mutation load through selection on males. Evolution 63:569–582.

Wickham, H. 2016. ggplot2: Elegant graphics for data analysis. Springer-Verlag New York.

Wickham, H., R. François, L. Henry, and K. Müller. 2019. dplyr: A Grammar of data manipulation.

Wickham, H., and L. Henry. 2019. tidyr: Easily Tidy Data with “spread()” and “gather()” functions.

Wingstrand, K. G. 1978. Comparative spermatology of the Crustacea Entomostraca; 1, Subclass Branchiopoda. Biologiske Skrifter 22:1–67.

Winsor, G. L., and D. J. Innes. 2002. Sexual reproduction in *Daphnia pulex* (Crustacea: Cladocera): Observations on male mating behaviour and avoidance of inbreeding. Freshwater Biology 47:441–450.

Wolterek, R. 1909. Weitere experimentelle Untersuchungen über Artveränderung, speziel über das Wesen quantitativer Artunterschiede bei Daphniden. Verhandlungen der deutschen zoologischen Gesellschaft 9:110–173.

Wright, S. 1982. The shifting balance theory and macroevolution. Annual review of genetics 16:01–19.

Wuerz, M., E. Huebner, and J. Huebner. 2017. The morphology of the male reproductive system, spermatogenesis and the spermatozoon of *Daphnia magna* (Crustacea: Branchiopoda). Journal of Morphology 278:1536–1550.

Zaffagnini, F. 1987. Reproduction in *Daphnia*. Page 280 in R. H. Peters and R. De Bernadi, eds. Daphnia (Vol. 45). Istituto Italiano di Idrobiologia, Pallanza.

Zahavi, A. 1975. Mate selection-A selection for a handicap. Journal of Theoretical Biology 53:205–214.

