## Supplementary material for "Estimation of the propensity for sexual selection in a cyclical parthenogen": Daphnia_sexual_selection_SupMat.html

Sexual selection in Daphnia


Code 

- Show All Code
- Hide All Code

### Sexual selection in Daphnia

###### by David Duneau

#### in 2019

### Table of contents

### Library, Data import and reformatting

#### Library

```
setwd("F:/Dropbox/0_work/001_Res/0_Proj/02_Dros")

library(fitdistrplus)
library(dabestr)
library(DT)
library(scales)
library(kableExtra)
library(multcomp)
library(lmtest)
library(multcomp)
library(spaMM)
library("survminer")
library(mclust)
library(plotly)
library(dplyr)
library(tidyr)
library(spaMM)
library(EBImage)
library(gridExtra)
library(grid)
library(gtable)
library(sjPlot)
library(lme4)
library(car)
library(moments)
library(vcd)
library(visreg)
library(related)
library(cowplot)
library(ggplotify)
library(gridGraphics)

grab_grob <- function(){
  grid.echo()
  grid.grab()
}

RIGHT = function(x,n){
  substring(x,nchar(x)-n+1)
}

LEFT = function(x,n){
  substring(x,1,nchar(x)-n+1)
}

 r2.corr.mer <- function(m) {
   lmfit <-  lm(model.response(model.frame(m)) ~ fitted(m))
   summary(lmfit)$r.squared
 }

 logit2prob <- function(logit){
  odds <- exp(logit)
  prob <- odds / (1 + odds)
  return(prob)
 }
```

```
SuperSmallfont= 6
Smallfont= 8
Mediumfont= 10
Largefont= 14
verylargefont = 16
pointsize= 0.7
linesize=0.35
meansize = 1.5
Margin=c(0,0,0,0)

fontsizeaxes = 12
fontsizeaxes2 = 10
```

#### Import

```
path.to.data <- "F:/Dropbox/0_work/001_Res/0_Proj/006_Sex_select_daphnia_projects/03_paper/Report_sup_mat/"

rm(d,path,model)
d <- list()
path <- list()
model <- list()
for(f in list.files(path=path.to.data,pattern="*.csv$",recursive=T,full.names=T)) {
  nom <- gsub(".*/(.*).csv","\\1",f)    
  cat(nom,"\n")
  path[[nom]] <- gsub("(.*)/.*csv","\\1/",f)
  d[[nom]] <- read.table(f,header=T,sep=";",dec=",")

}

all_data = d[["all_data_sexual_selection_Daphnia"]]


Data_male_couple_2011 = d[["Data_male_couple_2011_selection_Daphnia"]]
Data_sperm_2011 = d[["Data_sperm_2011_selection_Daphnia"]]
Data_mean_sperm_size = d[["Data_mean_sperm_size_selection_Daphnia"]]
Sperm_length_lab = d[["Sperm_length_lab_selection_Daphnia"]]

Sexratio_infection_NaturalPop_free= d[["Sexratio_inf_NatPop_free_sexual_selection_Daphnia"]]


SexbiasInfectionNaturalPop=
subset(all_data,!is.na(Infectious_status)) %>%
  group_by(Sampling_date,Population,Mating_status,Sex,Infectious_status)%>%
  summarise (n = n()) %>%
  mutate(freq = n / sum(n))

SexbiasInfectionNaturalPop = subset(SexbiasInfectionNaturalPop, Infectious_status=="1")
SexbiasInfectionNaturalPop = subset(SexbiasInfectionNaturalPop,select=-c(Infectious_status))
colnames(SexbiasInfectionNaturalPop)[ncol(SexbiasInfectionNaturalPop)] = "Infection_rate"
SexbiasInfectionNaturalPop = data.frame(SexbiasInfectionNaturalPop)

SexbiasInfectionNaturalPop_bis = subset(SexbiasInfectionNaturalPop,select=-c(n))

SexbiasInfectionNaturalPop1=
data.frame(SexbiasInfectionNaturalPop_bis %>%
 spread(Sex,Infection_rate))

sex_ratio_free = subset(SexbiasInfectionNaturalPop,Mating_status=="Free") 

SexbiasInfectionNaturalPop2=
 SexbiasInfectionNaturalPop_bis %>%
 spread(Mating_status,Infection_rate)


all_data$Prop_spine = with(all_data,
                      Spine_size/(Body_size+Spine_size)*100)
```

#### Summary data

---


---

### 1 - Mating formation

#### 1.1 - Occurence of males in studied populations

**What’s the proportion of males in the Finnish natural metapopulation? (Figure 1A)**  
We found that the proportion of males in summer was on average around 30%, ranging from 5 to 60 % of males (of all ages). Males can be frequent in the *Daphnia* metapopulation.

```
Name= "Proportion of males"
      
Plot_sex_ratio = 
  ggplot(Sexratio_infection_NaturalPop_free,
         aes(x=Mating_status,y=Male_ratio)) +
labs(caption = "(Figure 1A)")+
  geom_boxplot(size =0.6,outlier.shape=NA,fill=NA)+
  geom_dotplot(alpha=0.3,pch=19, binaxis = "y", stackdir = "center",binwidth = 0.03, show.legend=FALSE)+
  scale_y_continuous(Name,
                     limits=c(0,1),
                     breaks=(seq(0,1,by=0.2)))+
  scale_x_discrete("",
                   labels="All populations")+
  stat_summary(fun.y = mean, geom = "point",colour="red",size=2) +
  theme(axis.title.x = element_text(size=Mediumfont),
        axis.title.y = element_text(size=Mediumfont),
        axis.line.x = element_line(colour="black",size=0.75),
        axis.line.y = element_line(colour="black",size=0.75),
        axis.ticks.x = element_line(size = 0.75),
        axis.ticks.y = element_line(size = 0.75),
        axis.text.x = element_text(size=Mediumfont,colour="black"),
        axis.text.y = element_text(size=Smallfont,colour="black"),
        plot.margin = unit(Margin, "cm"),
        strip.text.x = element_text(size =Smallfont, colour = "black",face="italic"),
        strip.text.y = element_text(size =Smallfont, colour = "black",face="italic"),
        legend.direction = "vertical", 
        legend.box = "horizontal",
        legend.position = "none",
        legend.key.height = unit(0.4, "cm"),
        legend.key.width= unit(0.3, "cm"),
        legend.title = element_text(face="italic",size=Smallfont), 
        legend.key = element_rect(colour = 'white', fill = "white", linetype='dashed'),
        legend.text = element_text(size=Smallfont),
        legend.background = element_rect(fill=NA),
        panel.background = element_blank())+
  guides(shape=guide_legend(ncol=1),
         fill=guide_legend(ncol=1),
         col=guide_legend(ncol=1))

grid.draw(Plot_sex_ratio)
```

```
# svg(filename="F:/Dropbox/0_work/001_Res/0_Proj/006_Sex_select_daphnia_projects/04_presentation/Plot_sex_ratio.svg",
#     width=5,
#     height=4,
#     pointsize=12)
# Plot_sex_ratio
# dev.off()
```

#### 1.2 - Number of males in a mating

**How many males can be found in a same mating? (Figure 1B)**  
We found that the sexual process can occur with one (monandrous) or two males (polyandrous) at the same time. Overall, we rarely found three males in the same sexual process (7 times out of the 968 mating in the study). However, in populations where polyandrous matings were very frequent, the frequency of trio reached 2 (pop. SP1-5) and 3 % (pop. SP1-6).

```
img1 <- readImage("F:/Dropbox/0_work/001_Res/0_Proj/006_Sex_select_daphnia_projects/analysis/dmagna_mating_ventral_1_enhanced.jpg") 
grid.raster(img1)
```

Monandrous mating

```
img1 <- readImage("F:/Dropbox/0_work/001_Res/0_Proj/006_Sex_select_daphnia_projects/analysis/IMG_8401_b.jpg") 
grid.raster(img1)
```

Polyandrous mating (Figure 1B)

**Males are mostly attached to sexual females**  
We found that in 80 % of the cases (382/477) mating females showed the typical morphological changes of the brood pouch associated with the formation of a resting egg case (ephippium), suggesting that they were ready to mate.

```
Broodpouch_mating = with(subset(all_data,Mating_status == "Mating" & Sex=="Female"),
     table(Brood_pouch))

Broodpouch_mating_2 = with(subset(all_data,Mating_status == "Mating" & Sex=="Female"),
     table(Brood_pouch)/(Broodpouch_mating["Asex"]+Broodpouch_mating["Empty"]+Broodpouch_mating["Ephip"]))

Broodpouch_mating = cbind(Broodpouch_mating,Broodpouch_mating_2)


Broodpouch_mating= as.data.frame(Broodpouch_mating)
Broodpouch_mating$Content_brood_pouch = rownames(Broodpouch_mating)
Broodpouch_mating = Broodpouch_mating[,c(3,1,2)]
colnames(Broodpouch_mating) = c("Content_brood_pouch","Counts","Proportion")
Broodpouch_mating[, 1] =c("Asexual eggs","Empty","Ephippium")
Broodpouch_mating = Broodpouch_mating[c(3,1,2),]
Broodpouch_mating[, 3] = round(Broodpouch_mating[, 3],2)
rownames(Broodpouch_mating) = c()


plot_Broodpouch_mating=
ggplot(Broodpouch_mating,aes(x=
Content_brood_pouch,y=Proportion))+
  geom_bar(stat="identity",fill="black",alpha=0.3,col="black",width=0.5)+
labs(caption = "(Supplementary figure)")+
   scale_x_discrete("Content brood pouch\nfrom females mating",
                    limits=c("Ephippium","Asexual eggs","Empty"),
                    labels=c("Ephippium","Asexual","Empty")) +
      scale_y_continuous("Proportion",
                         limits=c(0,1.05),
                         breaks=(seq(0,1,by=0.2)))+
  geom_text(aes(label=paste("(",Counts,")",sep="")), vjust=-0.5, color="black", size=3.5)+
  theme(axis.title.x = element_text(size = Mediumfont,colour="black"),
        axis.title.y = element_blank(),
        axis.line.x = element_line(colour="black",size=0.75),
        axis.line.y = element_line(colour="black",size=0.75),
        axis.ticks.x = element_line(size = 0.75),
        axis.ticks.y = element_line(size = 0.75),
        axis.text.x = element_text(size=Mediumfont,colour="black"),
        axis.text.y = element_text(size=Mediumfont,colour="black"),
        plot.margin = unit(c(0,0.2,0.2,0), "cm"),
       panel.background = element_blank())
plot_Broodpouch_mating
```

#### 1.3 - Effect of body size on mating formation

```
tmp_M_single = subset(all_data,!is.na(Body_size)  & Sex =="Male" & Mating_status=="Free")
Mean_size_single_male_pop=
  tmp_M_single %>%
  group_by(Population)%>%     
  summarise(mean_size_single=mean(Body_size))
Mean_size_single_male_pop = data.frame(Mean_size_single_male_pop)

size_male =subset(all_data,Sex=="Male" & !is.na(Body_size))

size_male$Mean_size = Mean_size_single_male_pop[match(size_male$Population,Mean_size_single_male_pop$Population),2]

size_male$Body_size_centred = size_male$Body_size - size_male$Mean_size

size_male = size_male[,c(1:7,19,32,33)]

tmp_F_single = subset(all_data,!is.na(Body_size)  & Sex =="Female" & Mating_status=="Free")
Mean_size_single_female_pop=
  tmp_F_single %>%
  group_by(Population)%>%     
  summarise(mean_size_single=mean(Body_size))
Mean_size_single_female_pop = data.frame(Mean_size_single_female_pop)

size_female = subset(all_data,Sex=="Female" & !is.na(Body_size)  & Population!="SK-58")

size_female$Mean_size = Mean_size_single_female_pop[match(size_female$Population,Mean_size_single_female_pop$Population),2]

size_female$Body_size_centred = size_female$Body_size - size_female$Mean_size

size_female = size_female[,c(1:7,19,32,33)]

tmp_F_mating = subset(all_data,!is.na(Body_size)  & Sex =="Female" & Mating_status=="Mating")
Mean_size_mating_female_pop=
  tmp_F_mating %>%
  group_by(Population)%>%     
  summarise(mean_size_mating=mean(Body_size))
Mean_size_mating_female_pop = data.frame(Mean_size_mating_female_pop)

Mean_size_mating_female_pop$Mean_size_single = Mean_size_single_female_pop[match(Mean_size_mating_female_pop$Population,Mean_size_single_female_pop$Population),2]

Mean_size_mating_female_pop$Size_difference = Mean_size_mating_female_pop$mean_size_mating -Mean_size_mating_female_pop$Mean_size_single

Mean_size_mating_female_pop$Prop_larger = Mean_size_mating_female_pop$Size_difference/Mean_size_mating_female_pop$mean_size_mating*100

#mean(Mean_size_mating_female_pop$Prop_larger,na.rm=T)

###

tmp_M_mating = subset(all_data,!is.na(Body_size)  & Sex =="Male" & Mating_status=="Mating")
Mean_size_mating_male_pop=
  tmp_M_mating %>%
  group_by(Population)%>%     
  summarise(mean_size_mating=mean(Body_size))
Mean_size_mating_male_pop = data.frame(Mean_size_mating_male_pop)

Mean_size_mating_male_pop$Mean_size_single = Mean_size_single_male_pop[match(Mean_size_mating_male_pop$Population,Mean_size_single_male_pop$Population),2]

Mean_size_mating_male_pop$Size_difference = Mean_size_mating_male_pop$mean_size_mating -Mean_size_mating_male_pop$Mean_size_single

Mean_size_mating_male_pop$Prop_larger = Mean_size_mating_male_pop$Size_difference/Mean_size_mating_male_pop$mean_size_mating*100

#mean(Mean_size_mating_male_pop$Prop_larger,na.rm=T)
```

##### 1.3.1 Body length and sexual process

**Is there an advantage to be large to access to mating? Is there a size-related assortative mating? (Figure 2A)**  
We tested whether individuals larger than the average in their population had a better access to matings. To do so, we centred the body length of each sex by the mean of the individuals of the same sex caught randomly in the population.  
We found that larger individuals of both sexes generally had better access to matings. Females in mating were on average 9.5 % larger than those randomly caught in the population. Males in mating were on average 2.3 % larger than those randomly caught in the population. For females, this is probably due to the fact that it is generally the older females which produce ephippia. For males, this suggests that larger males have a better access to females. We found that, controlling for the averaged length by population, there was an assortative mating regarding the body length (Pearson correlation test: Rho = 0.15, p= 0.0001). The strength of the homogamy (15 %) is lower than the average strength regarding size-related homogamy accross animal taxa (31% according to Jiang, Bolnick, and Kirkpatrick (2013)) and it depends on the population.

```
Assort_mating_Size_grouped=
  ggplot(Size_couple, aes(x = Mean_size_female_centred, y = Mean_size_male_centred)) +
  labs(caption = "(Figure 2A)")+
  #  geom_point(aes(color=factor(Nbr_of_males),shape=factor(Nbr_of_males)))+ 
  geom_count(pch=19,alpha=0.3)+
  scale_size("N", breaks = c(1,5,10,20))+
  scale_x_continuous("Diff. body length mated females\nto the population average (mm)",
                     expand=c(0.01,0.01),
                     limits=c(-0.5, 1.7),
                     breaks=c(seq(-0.5,1.5,by=0.5)))+
  scale_y_continuous("Diff. body length mated males\nto the population average (mm)",
                     limits=c(-0.6, 0.6),
                     breaks=c(-0.6,-0.4,-0.2,0,0.2,0.4,0.6))+
  scale_color_manual("Nbr of males in mating",
                     values = c("black","blue","red"))+
  scale_shape_manual("Nbr of males in mating",
                     values = c(1,4,19))+
  geom_hline(yintercept = 0)+
  geom_vline(xintercept = 0)+
  annotate("text",x=1,y=-0.6,size=3.5,label="Rho= 0.15, p= 0.0001")+
  geom_smooth(method='lm',formula=y~x,size=1,color="black")+
  theme(axis.title.x = element_text(size=Mediumfont+4,colour="black"),
        axis.title.y = element_text(size=Mediumfont+4,colour="black"),
        axis.line.x = element_line(colour="black",size=0.75),
        axis.line.y = element_line(colour="black",size=0.75),
        axis.ticks.x = element_line(size = 0.75),
        axis.ticks.y = element_line(size = 0.75),
        axis.text.x = element_text(size=Mediumfont,colour="black"),
        axis.text.y = element_text(size=Mediumfont,colour="black"),
        plot.margin = unit(Margin, "cm"),  
        legend.direction = "horizontal", 
        legend.box = "vertical",
        legend.position = "bottom",
        legend.key.height = unit(0.4, "cm"),
        legend.key.width= unit(0.3, "cm"),
        legend.title = element_text(face="italic",size=Mediumfont+4), 
        legend.key = element_rect(colour = 'white', fill = "white", linetype='dashed'),
        legend.text = element_text(size=Mediumfont+4),
        legend.background = element_rect(fill=NA),
        panel.background = element_blank())+
  guides(shape=guide_legend(ncol=3),
         fill=guide_legend(ncol=3),
         col=guide_legend(ncol=3))


grid.draw(Assort_mating_Size_grouped)
```

```
# svg(filename="F:/Dropbox/0_work/001_Res/0_Proj/006_Sex_select_daphnia_projects/04_presentation/Assort_mating_Size_grouped.svg",
#     width=5,
#     height=5,
#     pointsize=12)
# Assort_mating_Size_grouped
# dev.off()
```

```
#anova(lm(Body_size_Male~Population, data=Size_couple))
# Male body length depends on the population so we removed the part of the variance in male body length that is explained by the population and correlated those residuals to the female body length.

with(Size_couple,
unlist(format(cor.test(Mean_size_female_centred, Mean_size_male_centred),digit=3)))%>%
  kable(col.names="Pearson correlation") %>%
  kable_styling(bootstrap_options = c("striped", "hover", "condensed"), full_width = F)
```

|  | Pearson correlation |
| --- | --- |
| statistic | 3.9 |
| parameter | 646 |
| p.value | 0.000105 |
| estimate | 0.152 |
| null.value | 0 |
| alternative | two.sided |
| method | Pearson’s product-moment correlation |
| data.name | Mean\_size\_female\_centred and Mean\_size\_male\_centred |
| conf.int | 0.0757, 0.2262 |

##### 1.3.2 Advantage body length (Raw data)

**Are individuals mating larger than individuals single? (Raw data - supp)**

```
Plot_size_mating_F =
  ggplot(subset(all_data,Sex=="Female" & !is.na(Body_size)  & Population!="SK-58"),
         aes(x=Mating_status,y=Body_size)) +
  facet_wrap(.~Population,nrow=2)+
  geom_boxplot(aes(color=Sex),size =0.6,outlier.shape=NA)+
  geom_dotplot(aes(color=Sex),fill=NA, binaxis = "y", stackdir = "center",dotsize=0.4, show.legend=FALSE)+
  labs(caption = "(Supplementary figure)")+
  scale_y_continuous("Body length - Female (mm)",
                     limits=c(1.5,4.3),
                     breaks=(seq(1.5,4,by=0.5)))+
  scale_x_discrete("Mating status",
                   labels=c("Single","Mating"))+
  scale_color_manual(values = c("red"))+
  stat_summary(fun.y = mean, geom = "point",colour="black",size=2) +
        theme(axis.title.x = element_text(size=Mediumfont+6),
        axis.title.y =  element_text(size=Mediumfont+6),
        axis.line.x = element_line(colour="black",size=0.75),
        axis.line.y = element_line(colour="black",size=0.75),
        axis.ticks.x = element_line(size = 0.75),
        axis.ticks.y = element_line(size = 0.75),
        axis.text.x = element_text(size=Mediumfont+4,colour="black"),
        axis.text.y = element_text(size=Mediumfont+4,colour="black"),
        plot.margin = unit(Margin, "cm"),
        legend.position = "none",
        strip.text.x = element_text(size =Mediumfont+6, colour = "black",face="italic"),
        strip.text.y = element_text(size =Mediumfont+6, colour = "black",face="italic"),
        panel.background = element_blank())

grid.draw(Plot_size_mating_F)
```

```
Plot_size_mating_M =
  ggplot(subset(all_data,Sex=="Male" & !is.na(Body_size) & !(Population %in% c("SK-58","SP1")) ),
         aes(x=Mating_status,y=Body_size)) +
  labs(caption = "(Supplementary figure)")+
  facet_wrap(.~Population,nrow=2)+
  geom_boxplot(aes(color=Sex),size =0.6,outlier.shape=NA)+
  geom_dotplot(aes(color=Sex),fill=NA, binaxis = "y", stackdir = "center",dotsize=0.4, show.legend=FALSE)+
  scale_y_continuous("Body length - Male (mm)",
                     limits=c(1.2,2.3),
                     breaks=(seq(1,2.2,by=0.2)))+
  scale_x_discrete("Mating status",
                   labels=c("Single","Mating"))+
  scale_color_manual(values = c("blue"))+
  stat_summary(fun.y = mean, geom = "point",colour="black",size=2) +
        theme(axis.title.x = element_text(size=Mediumfont+6),
        axis.title.y =  element_text(size=Mediumfont+6),
        axis.line.x = element_line(colour="black",size=0.75),
        axis.line.y = element_line(colour="black",size=0.75),
        axis.ticks.x = element_line(size = 0.75),
        axis.ticks.y = element_line(size = 0.75),
        axis.text.x = element_text(size=Mediumfont+4,colour="black"),
        axis.text.y = element_text(size=Mediumfont+4,colour="black"),
        plot.margin = unit(Margin, "cm"),
        legend.position = "none",
        strip.text.x = element_text(size =Mediumfont+6, colour = "black",face="italic"),
        strip.text.y = element_text(size =Mediumfont+6, colour = "black",face="italic"),
        panel.background = element_blank())

grid.draw(Plot_size_mating_M)
```

```
# svg(filename="F:/Dropbox/0_work/001_Res/0_Proj/006_Sex_select_daphnia_projects/04_presentation/Plot_size_mating_F.svg",
#     width=5,
#     height=5,
#     pointsize=12)
# Plot_size_mating_F
# dev.off()
# 
# svg(filename="F:/Dropbox/0_work/001_Res/0_Proj/006_Sex_select_daphnia_projects/04_presentation/Plot_size_mating_M.svg",
#     width=5,
#     height=5,
#     pointsize=12)
# Plot_size_mating_M
# dev.off()
```

```
#print("Full model= lmer(Body_size ~ Mating_status + (1|Population))")

print("For females")
data_temp = subset(all_data,Sex=="Female" & !is.na(Body_size) & Population!="SK-58")

# model_full = lmer(Body_size ~ Mating_status + (1|Population),data_temp) 
# effect_size = (fixef(model_full)[[2]]/fixef(model_full)[[1]]+fixef(model_full)[[2]])*100
# 
#   standard.res_init = (residuals(model_full)/sd(residuals(model_full)))
#   par(mfrow=c(2,2))
#   hist(residuals(model_full)) 
#   skewness(residuals(model_full)) # indeed, moderately positive skewness (0.42)
#   qqnorm(residuals(model_full)) # normality is not good
#   qqline(residuals(model_full))
# 
# ##box-cox transformation
# bc <- powerTransform(model_full,family="bcPower" )
# lambda <- bc$lambda # 0.27
# 
# model_full_transformed <- lmer(log(Body_size)~Mating_status + (1|Population),data_temp) #box-cox calculation gave me an idea of the good transformation but the usual transformation was not good enough
# 
# standard.res = (residuals(model_full_transformed)/sd(residuals(model_full_transformed)))
# par(mfrow=c(2,2))
# plot(data_temp$Mating_status,standard.res,ylab="standardized residuals",xlab="mating status",main="Heteroscedasticity") #no evidence for non linearity ad heteroscedasticity
# hist(residuals(model_full_transformed),main="distribution residuals")
# skewness(residuals(model_full_transformed)) # skewness was reduced by the transformation (-0.11)
# qqnorm(residuals(model_full),main="qqplot before transformation")
# qqline(residuals(model_full))
# qqnorm(residuals(model_full_transformed),main="qqplot after transformation") # normality was still bad
# qqline(residuals(model_full_transformed))


residuals_size_pop_F = residuals(lm(Body_size~ Population,data_temp))
data_temp$residuals_size_pop_F = residuals_size_pop_F


unlist(format(wilcox.test(residuals_size_pop_F ~ Mating_status,data_temp),digit=3))%>%
  kable() %>%
  kable_styling(bootstrap_options = c("striped", "hover", "condensed"), full_width = F)

data_temp=NULL

##################################################################################################################
print("For males")
data_temp = subset(all_data,Sex=="Male" & !is.na(Body_size) & !(Population %in% c("SK-58","SP1")))

#  model_full = lmer(Body_size ~ Mating_status + (1|Population),data_temp) 
#  effect_size = (fixef(model_full)[[2]]/fixef(model_full)[[1]]+fixef(model_full)[[2]])*100
# 
#   standard.res_init = (residuals(model_full)/sd(residuals(model_full)))
#   par(mfrow=c(2,2))
#   hist(residuals(model_full))
#  skewness(residuals(model_full)) # indeed, a small negative skewness (-0.41)
#  qqnorm(residuals(model_full)) # normality is not good
#  qqline(residuals(model_full))
# 
# #box-cox transformation
# bc <- powerTransform(model_full,family="bcPower" )
# lambda <- bc$lambda # 1.23 
# 
#  model_full_transformed <- lmer((Body_size)^2~Mating_status + (1|Population),data_temp) #box-cox calculation gave me an idea of the good transformation but the usual transformation was not good enough
# 
#  standard.res = (residuals(model_full_transformed)/sd(residuals(model_full_transformed)))
#  par(mfrow=c(2,2))
#  plot(data_temp$Mating_status,standard.res,ylab="standardized residuals",xlab="mating status",main="Heteroscedasticity") #no evidence for non linearity ad heteroscedasticity
#  hist(residuals(model_full_transformed),main="distribution residuals")
#  skewness(residuals(model_full_transformed)) # skewness was abolished(-0.14)
#  qqnorm(residuals(model_full),main="qqplot before transformation")
#  qqline(residuals(model_full))
#  qqnorm(residuals(model_full_transformed),main="qqplot after transformation") # normality was improved and is now acceptable
#  qqline(residuals(model_full_transformed))
# 
#  model_full_transformed_Male = model_full_transformed
#  model_reduced_transformed_Male <-  lmer((Body_size^lambda-1)/lambda~ 1 + (1|Population),data_temp)

residuals_size_pop_M = residuals(lm(Body_size~ Population,data_temp))
data_temp$residuals_size_pop_M = residuals_size_pop_M


unlist(format(wilcox.test(residuals_size_pop_M ~ Mating_status,data_temp),digit=3))%>%
  kable() %>%
  kable_styling(bootstrap_options = c("striped", "hover", "condensed"), full_width = F)

data_temp=NULL
```

##### 1.3.3 Assortative mating (Raw data)

**Is there an assortative mating based on body length? (Raw data - supp)**  
(Dot size illustrate the number of pairs sharing the same length)

```
Assort_mating_Size=
  ggplot(Size_couple, aes(x = Body_size_Female, y = Body_size_Male)) +
  labs(caption = "(Supplementary figure)")+
  geom_point(aes(color=factor(Nbr_of_males),shape=factor(Nbr_of_males)))+ 
  geom_count()+
  scale_size("N", breaks = c(1,5,10,20))+
  facet_wrap(.~Population, scales = "free",nrow=4)+
  scale_x_continuous("Body length - females (mm)",
                     expand=c(0.01,0.01))+
  scale_y_continuous("Body length - males (mm)")+
  scale_color_manual("Nbr of males in mating",
                     values = c("black","blue","red"))+
  scale_shape_manual("Nbr of males in mating",
                     values = c(1,4,19))+
  geom_smooth(method='lm',formula=y~x,size=1,color="black")+
  theme(axis.title.x = element_text(size=Mediumfont,colour="black"),
        axis.title.y = element_text(size=Mediumfont,colour="black"),
        axis.line.x = element_line(colour="black",size=0.75),
        axis.line.y = element_line(colour="black",size=0.75),
        axis.ticks.x = element_line(size = 0.75),
        axis.ticks.y = element_line(size = 0.75),
        axis.text.x = element_text(size=Mediumfont,colour="black"),
        axis.text.y = element_text(size=Mediumfont,colour="black"),
        plot.margin = unit(Margin, "cm"),
        strip.text.y = element_text(size =Mediumfont, colour = "black",face="italic"),
        legend.direction = "horizontal", 
        legend.box = "vertical",
        legend.position = "bottom",
        legend.key.height = unit(0.4, "cm"),
        legend.key.width= unit(0.3, "cm"),
        legend.key = element_rect(colour = 'white', fill = "white", linetype='dashed'),
        legend.text = element_text(size=Mediumfont),
        legend.title = element_text(face="italic",size=Mediumfont), 
        legend.background = element_rect(fill=NA),
        panel.background = element_blank())+
  guides(shape=guide_legend(ncol=3),
         fill=guide_legend(ncol=3),
         col=guide_legend(ncol=3))

grid.draw(Assort_mating_Size)
```

```
# svg(filename="F:/Dropbox/0_work/001_Res/0_Proj/006_Sex_select_daphnia_projects/04_presentation/Assort_mating_Size.svg",
#     width=5,
#     height=5,
#     pointsize=12)
# Assort_mating_Size
# dev.off()
```

---

##### 1.3.4 Hitchhicking

**Are females in polyandrous mating carrying smaller males? (Figure 2B)**  
Controlling for the average body length of mating males in each population, we tested whether females with two males carried smaller males than those carrying one male. Males were significantly smaller in polyandrous matings suggesting that smaller males could get access to matings when females are already mating (Wilcoxon test: df=1, W= 83062, p= 0.01). However, on the one hand, the difference is very small (1.806mm in monandrous versus 1.798mm in polyandrous), on the other hand, polyandrous matings per definition include a first male attaching who is expected to be of the same size as males in monandrous matings. This seems to be illustrated by the two groups in the violin plots of males in polyandrous matings.

```
populationWoDiversMatingType = c("Fu-29","Langskar","SK-58","SP1")

data_size_males= subset(all_data,!is.na(Body_size)  & Sex =="Male" & Mating_status=="Mating" & !is.na(Nbr_of_males) & Nbr_of_males!=3 & !(Population %in%populationWoDiversMatingType))

Mean_data_size_mating_males=
  data_size_males %>%
  group_by(Population,Nbr_of_males)%>%     
  summarise(mean_size_mating_males=mean(Body_size))%>%
  spread(key = Nbr_of_males,value = mean_size_mating_males)

Mean_data_size_mating_males = data.frame(Mean_data_size_mating_males)
Mean_data_size_mating_males$Difference = Mean_data_size_mating_males[,2]-Mean_data_size_mating_males[,3]
Mean_data_size_mating_males$Prop = Mean_data_size_mating_males$Difference/Mean_data_size_mating_males[,2]*100

#mean(Mean_data_size_mating_males$Prop,na.rm=T)

residuals_size_pop = residuals(lm(Body_size~ Population,data_size_males))
data_size_males$residuals_size_pop = residuals_size_pop

Size_double_mating_grouped=
  ggplot(data_size_males,aes(x=as.factor(Nbr_of_males),y=residuals_size_pop)) +
  labs(caption = "(Supplementary figure 5)")+
  # geom_boxplot(fill=NA,size =0.6,outlier.shape=NA)+
  geom_violin(draw_quantiles = c(0.25, 0.5, 0.75),colour="darkgrey",adjust=1)+
  geom_dotplot(aes(color= Population),binaxis = "y", stackdir = "center",dotsize=0.2,fill=NA, show.legend=FALSE)+
  scale_y_continuous("Diff. body length to the average of mating males (mm)")+
  scale_x_discrete("Mating type",
                   expand=c(0.1,0.1),
                   limits=c(1,2),
                   labels=c("Monandrous","Polyandrous"))+
  stat_summary(fun.y = mean, geom = "point",colour="red",size=2) +
  annotate("text",x=1.5,y=0.38,label="*",size=7)+
  theme(axis.title.x = element_text(size=Mediumfont+4),
        axis.title.y = element_text(size=Mediumfont+4),
        axis.line.x = element_line(colour="black",size=0.75),
        axis.line.y = element_line(colour="black",size=0.75),
        axis.ticks.x = element_line(size = 0.75),
        axis.ticks.y = element_line(size = 0.75),
        axis.text.x = element_text(size=Mediumfont,colour="black"),
        axis.text.y = element_text(size=Mediumfont,colour="black"),
        plot.margin = unit(Margin, "cm"),
        panel.background = element_blank())

grid.draw(Size_double_mating_grouped)

# svg(filename="F:/Dropbox/0_work/001_Res/0_Proj/006_Sex_select_daphnia_projects/04_presentation/Size_double_mating_grouped.svg",
#     width=5,
#     height=5,
#     pointsize=12)
# Size_double_mating_grouped
# dev.off()
```

```
populationWoDiversMatingType = c("Fu-29","Langskar","SK-58","SP1")

data_size_males= subset(all_data,!is.na(Body_size)  & Sex =="Male" & Mating_status=="Mating" & !is.na(Nbr_of_males) & Nbr_of_males!=3 & !(Population %in%populationWoDiversMatingType))

Mean_data_size_mating_males=
  as.data.frame(data_size_males %>%
  group_by(Population,Nbr_of_males)%>%     
  summarise(mean_size_mating_males=mean(Body_size),sd_size_mating_males=sd(Body_size)))

plot_size_mating_M=
ggplot(Mean_data_size_mating_males, aes(as.factor(Nbr_of_males),mean_size_mating_males, group=Population,color=Population)) +
  labs(caption = "(Figure 2B)")+
  geom_path(size=0.5,position = position_dodge(width=0.4),color="black")+
  geom_point(shape=16,size= 1,position = position_dodge(width=0.4))+
  geom_errorbar(aes(ymax = mean_size_mating_males+sd_size_mating_males, ymin = mean_size_mating_males-sd_size_mating_males),linetype=1,size=1,width=0.5,position = position_dodge(width=0.4))+
  scale_x_discrete("Mating type",
                     expand=c(0.1,0.1),
                   limits=c("1","2"),
                   labels=c("Monandrous", "Polyandrous"))+
  scale_y_continuous("Mean body length - males (mm)",
                     limits=c(1.5, 2.22),
                     breaks=c(c(seq(1.6,2.25,by=0.2))))+
  annotate("text", label="ns", x= 1.5, y=2.1,size=4)+
  scale_color_viridis_d("Population",end = 0.8)+
  theme(
    axis.title.x = element_text(size=Mediumfont),
    axis.title.y = element_text(size=Mediumfont),
    axis.line.x = element_line(colour="black",size=0.75),
    axis.line.y = element_line(colour="black",size=0.75),
    axis.ticks.x = element_line(size = 0.75),
    axis.ticks.y = element_line(size = 0.75),
    axis.text.x = element_text(size=Mediumfont,colour="black"),
    axis.text.y = element_text(size=Mediumfont,colour="black"),
    # plot.margin = unit(Margin, "cm"),
   legend.position = "none",
    panel.background = element_blank())

grid.draw(plot_size_mating_M)
```

```
# svg(filename="F:/Dropbox/0_work/001_Res/0_Proj/006_Sex_select_daphnia_projects/04_presentation/plot_size_mating_M.svg",
#     width=5,
#     height=5,
#     pointsize=12)
# plot_size_mating_M
# dev.off()
```

```
populationWoDiversMatingType = c("Fu-29","Langskar","SK-58","SP1")

data_temp= subset(all_data,!is.na(Body_size)  & 
                    Sex =="Male" & 
                    Mating_status=="Mating" &
                    !is.na(Nbr_of_males) & 
                    Nbr_of_males!=3 & 
                    !(Population %in%populationWoDiversMatingType))
data_temp$Nbr_of_males = factor(data_temp$Nbr_of_males)

data_temp$Population = as.factor(data_temp$Population)


model_mating_type_full = fitme(Body_size~ as.factor(Nbr_of_males) + (1|Population),family=gaussian(link=identity), resid.model= ~ Population, data=data_temp)

model_mating_type_reduced = fitme(Body_size~ 1 + (1|Population),family=gaussian(link=identity), resid.model= ~ Population, data=data_temp)

test=anova(model_mating_type_full,model_mating_type_reduced) 

Res_size_mat_type= data.frame(Variable=as.character("Mating type"),
                              chi2_LR=as.numeric(test$basicLRT$chi2_LR),
                              df=as.numeric(test$basicLRT$df),
                              Pvalue=as.numeric(test$basicLRT$p_value),
                              Effect_mm=as.numeric(model_mating_type_full$fixef[2]))


format(Res_size_mat_type,digit=2)%>%
  kable(col.names = c("", "chi^2 LRT", "df", "p-value", "Effect (mm)")) %>%
  add_header_above(c("fitme(Body_size~ as.factor(Nbr_of_males) + (1|Population),\nfamily=gaussian(link=identity), resid.model= ~ Population)" = 5))%>%
  kable_styling(bootstrap_options = c("striped", "hover", "condensed"), full_width = F)
```

| fitme(Body\_size~ as.factor(Nbr\_of\_males) + (1\|Population), family=gaussian(link=identity), resid.model= ~ Population) | | | | |
| --- | --- | --- | --- | --- |
|  | chi^2 LRT | df | p-value | Effect (mm) |
| Mating type | 2.3 | 1 | 0.13 | -0.011 |

---

**Are females in polyandrous mating carrying smaller males? (Raw data - supp)**

```
populationWoDiversMatingType = c("Fu-29","Langskar","SK-58","SP1")

data_size_males= subset(all_data,!is.na(Body_size)  & Sex =="Male" & !is.na(Nbr_of_males) & Nbr_of_males!=3 & !(Population %in%populationWoDiversMatingType))

Size_double_mating=
  ggplot(data_size_males,aes(x=as.factor(Nbr_of_males),y=Body_size)) +
  labs(caption = "(Supplementary figure)")+
  geom_boxplot(fill=NA,size =0.6,outlier.shape=NA)+
  geom_dotplot(binaxis = "y", stackdir = "center",dotsize= 0.8, fill=NA,color="blue", show.legend=FALSE)+
  facet_wrap(.~Population,ncol=5,scales="free_y")+
  scale_y_continuous("Body length - males (mm)")+
  scale_x_discrete("Mating type",
                   expand=c(0.1,0.1),
                   limits=c(1,2),
                   labels=c("Monandrous","Polyandrous"))+
  stat_summary(fun.y = mean, geom = "point",colour="red",size=1) +
  theme(axis.title.x = element_text(size=12),
        axis.title.y = element_text(size=12),
        axis.line.x = element_line(colour="black",size=0.75),
        axis.line.y = element_line(colour="black",size=0.75),
        axis.ticks.x = element_line(size = 0.75),
        axis.ticks.y = element_line(size = 0.75),
        axis.text.x = element_text(size=12,colour="black"),
        axis.text.y = element_text(size=12,colour="black"),
        plot.margin = unit(Margin, "cm"),
        strip.text.y = element_text(size =12, colour = "black",face="italic"),
        panel.background = element_blank())+
  guides(shape=guide_legend(ncol=1),
         fill=guide_legend(ncol=1),
         col=guide_legend(ncol=1))

grid.draw(Size_double_mating)
```

```
# svg(filename="F:/Dropbox/0_work/001_Res/0_Proj/006_Sex_select_daphnia_projects/04_presentation/Size_double_mating.svg",
#     width=5,
#     height=5,
#     pointsize=12)
# Size_double_mating
# dev.off()

# 
# tmp = subset(all_data,!is.na(Body_size)  & Sex =="Male" & !is.na(Nbr_of_males))
# tmp$Nbr_of_males[grep("1", tmp$Nbr_of_males)] = "Monandrous"
# tmp$Nbr_of_males[grep("2", tmp$Nbr_of_males)] = "Polyandrous"
# tmp$Nbr_of_males = as.factor(tmp$Nbr_of_males)
# 
# test=
# tmp%>%
#          dabest(Nbr_of_males,Body_size,
#                 idx=c("Monandrous","Polyandrous"),
#                 paired=F)
# 
# plot(test,
#      palette = "Dark2",
#      rawplot.ylabel = "Body length - males (mm)",
#      effsize.ylabel = "Mean difference and bootsraped CI",
#      axes.title.fontsize=18,
#      tick.fontsize=16)

populationWoDiversMatingType = c("Fu-29","Langskar","SK-58","SP1")

data_temp= subset(all_data,!is.na(Body_size)  & 
                    Sex =="Male" & 
                    !is.na(Nbr_of_males) & 
                    Nbr_of_males!=3 & 
                    !(Population %in%populationWoDiversMatingType))
data_temp$Nbr_of_males = factor(data_temp$Nbr_of_males)

#model_full = lmer(Body_size~ Nbr_of_males + (1|Population),data_temp)

#standard.res_init = (residuals(model_full)/sd(residuals(model_full)))
# par(mfrow=c(2,2))
# hist(residuals(model_full))
# skewness(residuals(model_full)) #  skewness (-0.26)
# qqnorm(residuals(model_full)) # not normality distributed
# qqline(residuals(model_full))

##box-cox transformation
# bc <- powerTransform(model_full,family="bcPower" )
# lambda <- bc$lambda # 0.68 
# 
# model_full_transformed <- lmer((Body_size^lambda-1)/lambda~Nbr_of_males+ (1|Population),data_temp)
# 
# standard.res = (residuals(model_full_transformed)/sd(residuals(model_full_transformed)))
# par(mfrow=c(2,2))
# plot(data_temp$Nbr_of_males,standard.res,ylab="standardized residuals",xlab="Nbr of males",main="Heteroscedasticity") #no evidence for non linearity and heteroscedasticity
# hist(residuals(model_full_transformed),main="distribution residuals")
# skewness(residuals(model_full_transformed)) # acceptable minor skewness to the left (-0.38)
# qqnorm(residuals(model_full),main="qqplot before transformation")
# qqline(residuals(model_full))
# qqnorm(residuals(model_full_transformed),main="qqplot after transformation") # normality was improved
# qqline(residuals(model_full_transformed))
```

---

##### 1.3.5 Male size and mating duration

**Are larger males able to stay longer on females than smaller males? (Figure 2C)**

```
table_couple = all_data[,c("Sex","Mating_status","ID_mating","Nbr_of_males","Position_detached","Body_size")]
table_couple = subset(table_couple,Mating_status=="Mating" & !is.na(Position_detached) & Nbr_of_males==2 & Position_detached!=0)


list_remove = with(table_couple[which(is.na(table_couple$Body_size)),],
                   unique(ID_mating))
'%!in%' <- function(x,y)!('%in%'(x,y))
table_couple = filter(table_couple, ID_mating %!in% list_remove)

table_couple=
  table_couple %>%
  arrange(ID_mating)

table_couple$sub_ID = rep(c(1,2),nrow(table_couple)/2)

table_couple_spread = 
  table_couple[,-5]%>%
  spread(sub_ID,Body_size)
colnames(table_couple_spread)[5] = "Bodysize_male_1"
colnames(table_couple_spread)[6] = "Bodysize_male_2"


table_couple_spread$Difference_body_length = 
  with(table_couple_spread,
       Bodysize_male_1-Bodysize_male_2
       )

means_diff_bodysize = mean(table_couple_spread$Difference_body_length,na.rm=T)
se <- function(x) sqrt(var(x)/length(x))
se_diff_bodysize = se(table_couple_spread$Difference_body_length)


Distribution_body_difference=
  ggplot(table_couple_spread,aes(x=Difference_body_length)) +
  geom_histogram(colour="black",fill="black",alpha=0.3,binwidth = 0.05)+
  geom_point(x= means_diff_bodysize, y = 18, shape=16,size= 0.6, col="red")+
  geom_errorbarh(aes(xmax = means_diff_bodysize + se_diff_bodysize , xmin =means_diff_bodysize - se_diff_bodysize ,y=18),linetype=1,size=0.6, col="red")+
  labs(caption = "(Figure 2C)")+
  scale_y_continuous("Count",
                     expand=c(0,0.1),
                     limits=c(0,25),
                     breaks=(c(seq(0,25,by=5))))+
  scale_x_continuous("Difference body length\n(first minus second in mm)",
                     limits=c(-0.30,0.40),
                     breaks=(c(seq(-0.2,0.3,by=0.1))))+
  theme(axis.title.x = element_text(size = Mediumfont,colour="black"),
        axis.title.y = element_text(size = Mediumfont,colour="black"),
        axis.line.x = element_line(colour="black",size=0.75),
        axis.line.y = element_line(colour="black",size=0.75),
        axis.ticks.x = element_line(size = 0.75),
        axis.ticks.y = element_line(size = 0.75),
        axis.text.x = element_text(size=Mediumfont,colour="black"),
        axis.text.y = element_text(size=Smallfont,colour="black"),
        plot.margin = unit(c(1.5,0.3,0,0), "cm"),
        panel.background = element_blank())
grid.draw(Distribution_body_difference)
```

```
# svg(filename="F:/Dropbox/0_work/001_Res/0_Proj/006_Sex_select_daphnia_projects/04_presentation/Distribution_body_difference.svg",
#     width=5,
#     height=5,
#     pointsize=12)
# Distribution_body_difference
# dev.off()
```

```
test= t.test(table_couple_spread$Bodysize_male_1,table_couple_spread$Bodysize_male_2, paired=T)
test2= t.test(table_couple_spread$Bodysize_male_1,table_couple_spread$Bodysize_male_2, paired=T, alternative="less")

Res_size_mat_type= data.frame(Variable=as.character("Order detached"),
                              df=as.numeric(test$parameter),
                              chi2_LR=as.numeric(test$statistic),
                              Effect_mm=as.numeric(test$estimate),
                              Pvalue_two_tail=as.numeric(test$p.value),
                              Pvalue_one_tail=as.numeric(test2$p.value))


format(Res_size_mat_type,digit=2)%>%
  kable(col.names = c("","df", "t",  "Effect (mm)", "p-value\n2-tail","p-value\nfirst is smaller")) %>%
  add_header_above(c("paired t-test" = 6))%>%
  kable_styling(bootstrap_options = c("striped", "hover", "condensed"), full_width = F)
```

| paired t-test | | | | | |
| --- | --- | --- | --- | --- | --- |
|  | df | t | Effect (mm) | p-value 2-tail | p-value first is smaller |
| Order detached | 97 | -2.2 | -0.023 | 0.033 | 0.016 |

---

##### 1.3.6 Spine length and sexual process

```
tmp_M_single = subset(all_data,!is.na(Spine_size)  & Sex =="Male" & Mating_status=="Free")

Mean_spine_size_single_male_pop=
  tmp_M_single %>%
  group_by(Population)%>%     
  summarise(mean_prop_spine_size_single=mean(Prop_spine))
Mean_spine_size_single_male_pop = data.frame(Mean_spine_size_single_male_pop)

spine_size_male =subset(all_data,Sex=="Male" & !is.na(Spine_size))

spine_size_male$Mean_prop_spine_size = Mean_spine_size_single_male_pop[match(spine_size_male$Population,Mean_spine_size_single_male_pop$Population),2]

spine_size_male$Prop_spine_size_centred = spine_size_male$Prop_spine - spine_size_male$Mean_prop_spine_size

spine_size_male = spine_size_male[,c(1:7,32:34)]


###
tmp_F_single = subset(all_data,!is.na(Spine_size)  & Sex =="Female" & Mating_status=="Free")

Mean_spine_size_single_female_pop=
  tmp_F_single %>%
  group_by(Population)%>%     
  summarise(mean_prop_spine_size_single=mean(Prop_spine))
Mean_spine_size_single_female_pop = data.frame(Mean_spine_size_single_female_pop)

spine_size_female =subset(all_data,Sex=="Female" & !is.na(Spine_size) & Population!="SK-58")

spine_size_female$Mean_prop_spine_size = Mean_spine_size_single_female_pop[match(spine_size_female$Population,Mean_spine_size_single_female_pop$Population),2]

spine_size_female$Prop_spine_size_centred = spine_size_female$Prop_spine - spine_size_female$Mean_prop_spine_size

spine_size_female = spine_size_female[,c(1:7,32:34)]
```

**Is spine a secondary sexual trait ? (Supplementary figure)**  
We tested whether individuals with larger spine (proportionnally to their body size) than the average in its population were more often found mating. To do so, we centred the spine length relative to body size of each sex caught in mating by the mean of the individuals of the same sex caught randomly in the population.  
We found that spines were generally smaller for mating individuals, suggesting that spine is not a secondary sexual trait.  
Additionnalyly, males with proportionnally large spines where in mating with females with spine proportionnally larger spines than the average.

```
Assort_mating_Prop_Spine_Size_grouped=
  ggplot(Prop_spine_couple, aes(x = Mean_prop_spine_size_female_centred, y = Mean_prop_spine_size_male_centred)) +
  labs(caption = "(Supplementary figure)")+
  geom_count(pch=19,alpha=0.3)+
  scale_size("N", breaks = c(1,2,3,4,5))+
  scale_x_continuous("Diff. prop. spine length mated females\nto the population average",
                      expand=c(0.01,0.01),
                      limits=c(-20, 6),
                      breaks=c(seq(-20,6,by=5)))+
   scale_y_continuous("Diff. prop. spine length mated males\nto the population average",
                      expand=c(0.01,0.01),
                      limits=c(-23, 16),
                      breaks=c(seq(-25,20,by=5)))+
  geom_hline(yintercept = 0)+
  geom_vline(xintercept = 0)+
  geom_smooth(method='lm',formula=y~x,size=1,color="black")+
  theme(axis.title.x = element_text(size=Mediumfont,colour="black"),
        axis.title.y = element_text(size=Mediumfont,colour="black"),
        axis.line.x = element_line(colour="black",size=0.75),
        axis.line.y = element_line(colour="black",size=0.75),
        axis.ticks.x = element_line(size = 0.75),
        axis.ticks.y = element_line(size = 0.75),
        axis.text.x = element_text(size=Mediumfont,colour="black"),
        axis.text.y = element_text(size=Mediumfont,colour="black"),
        plot.margin = unit(Margin, "cm"),  
        legend.direction = "horizontal", 
        legend.box = "vertical",
        legend.position = "bottom",
        legend.key.height = unit(0.4, "cm"),
        legend.key.width= unit(0.3, "cm"),
        legend.title = element_text(face="italic",size=Smallfont), 
        legend.key = element_rect(colour = 'white', fill = "white", linetype='dashed'),
        legend.text = element_text(size=Mediumfont),
        legend.background = element_rect(fill=NA),
        panel.background = element_blank())+
  guides(shape=guide_legend(ncol=3),
         fill=guide_legend(ncol=3),
         col=guide_legend(ncol=3))

grid.draw(Assort_mating_Prop_Spine_Size_grouped)
```

```
# 
# svg(filename="F:/Dropbox/0_work/001_Res/0_Proj/006_Sex_select_daphnia_projects/04_presentation/Assort_mating_Prop_Spine_Size_grouped.svg",
#     width=5,
#     height=5,
#     pointsize=12)
# Assort_mating_Prop_Spine_Size_grouped
# dev.off()
```

```
# both sexes
model_prop_full=  fitme(Spine_size~Body_size + Sex + Mating_status + (1|Population), resid.model= ~ Population,family=gaussian(link=identity), data=subset(all_data, !is.na(Mating_status) & !is.na(Spine_size) & Population!="SK-58"))

model_prop_reduced=  fitme(Spine_size~Body_size + Sex + 1 +(1|Population), resid.model= ~ Population, family=gaussian(link=identity), data=subset(all_data,!is.na(Mating_status) & !is.na(Spine_size) & Population!="SK-58"))

# Chi2_LRT <- 2*(model_prop_full$APHLs[["p_v"]]-model_prop_reduced$APHLs[["p_v"]])
# pchisq(Chi2_LRT,df=1,lower.tail = F)

test = anova(model_prop_full,model_prop_reduced)

# Females
model_prop_full_F=  fitme(Spine_size~Body_size +  Mating_status + (1|Population), resid.model= ~ Population,family=gaussian(link=identity), data=subset(all_data,Sex=="Female" & !is.na(Mating_status) & !is.na(Spine_size) & Population!="SK-58"))

model_prop_reduced_F=  fitme(Spine_size~Body_size + 1 +(1|Population), resid.model= ~ Population, family=gaussian(link=identity), data=subset(all_data,Sex=="Female" & !is.na(Mating_status) & !is.na(Spine_size) & Population!="SK-58"))
test_F = anova(model_prop_full_F,model_prop_reduced_F)

# Males
model_prop_full_M=  fitme(Spine_size~Body_size +  Mating_status + (1|Population), resid.model= ~ Population,family=gaussian(link=identity), data=subset(all_data,Sex=="Male" & !is.na(Mating_status) & !is.na(Spine_size) & Population!="SK-58"))

model_prop_reduced_M=  fitme(Spine_size~Body_size + 1 +(1|Population), resid.model= ~ Population, family=gaussian(link=identity), data=subset(all_data,Sex=="Male" & !is.na(Mating_status) & !is.na(Spine_size) & Population!="SK-58"))
test_M = anova(model_prop_full_M,model_prop_reduced_M)


#######
Res_spine_mat_status= data.frame(Variable=as.character("Mating Status"),
                              chi2_LR=as.numeric(test$basicLRT$chi2_LR),
                              df=as.numeric(test$basicLRT$df),
                              Pvalue=as.numeric(test$basicLRT$p_value),
                              Effect_mm=as.numeric(model_prop_full$fixef[4]))

format(Res_spine_mat_status,digit=2)%>%
  kable(col.names = c("", "chi^2 LRT", "df", "p-value", "Effect Mating")) %>%
  add_header_above(c("fitme(Spine_size~Body_size + Sex + Mating_status + (1|Population),\nresid.model= ~ Population, family=gaussian(link=identity))" = 5))%>%
  kable_styling(bootstrap_options = c("striped", "hover", "condensed"), full_width = F)
```

| fitme(Spine\_size~Body\_size + Sex + Mating\_status + (1\|Population), resid.model= ~ Population, family=gaussian(link=identity)) | | | | |
| --- | --- | --- | --- | --- |
|  | chi^2 LRT | df | p-value | Effect Mating |
| Mating Status | 6.5 | 1 | 0.011 | -0.015 |

---

#### 1.4 - Effect of inbreeding in mating formation

**Is there inbreeding avoidance? (Supplementary figure)**

```
#(see taylor Ecol Evol 2015, very good to understand limits of the approach)
# triadic likelihood (TrioML) was the methods which correlated best with simulated true relatedness in this paper as for us.

data_genotype = subset(all_data,!is.na(Genotype))

data_genotype= data_genotype[,c("Sampling_date", "Population", "Sex", "Infectious_status", "Mating_status", "ID_mating", "Nbr_of_males","Marker_12", "Marker_173", "Marker_208", "Marker_B075", "Genotype")]

data_genotype$ID = 
with( data_genotype, 
      paste(ifelse(as.character(Sex)=="Female","FE","MA"),
            ifelse(as.character(Mating_status)=="Mating","Mt","Fr"),
            ID_mating,
            RIGHT(as.character(rownames(data_genotype)),2),sep="_"))

data_genotype$Genotype = with(data_genotype,
                              paste(Marker_12,Marker_173,Marker_208,Marker_B075))

data_genotype$Genotype = gsub("_"," ",data_genotype$Genotype)

data_genotype$Genotype = gsub("NA","0 0",data_genotype$Genotype)

data_genotype$Genotype = factor(data_genotype$Genotype)

# Export
 data_genotype_sp1 = subset(data_genotype,Population=="SP1-6" | Population=="SP1-5" )
 
write.table(data_genotype_sp1[,c(13,12)],"F:/Dropbox/0_work/001_Res/0_Proj/006_Sex_select_daphnia_projects/Genotyping/Coancestry.20180730/Genotype_data_coancestry_SP1.txt", row.names=FALSE,  col.names=FALSE,sep=" ",quote=F)

data_genotype_sp1_5 = subset(data_genotype,Population=="SP1-5")
data_genotype_sp1_5_free = subset(data_genotype_sp1_5,Mating_status=="Free")
data_genotype_sp1_5_Mating = subset(data_genotype_sp1_5,Mating_status=="Mating")

write.table(data_genotype_sp1_5[,c(13,12)],"F:/Dropbox/0_work/001_Res/0_Proj/006_Sex_select_daphnia_projects/Genotyping/Coancestry.20180730/Genotype_data_coancestry_SP1_5.txt", row.names=FALSE,  sep=" ",col.names=FALSE,quote=F)

data_genotype_sp1_6 = subset(data_genotype,Population=="SP1-6")
 write.table(data_genotype_sp1_6[,c(13,12)],"F:/Dropbox/0_work/001_Res/0_Proj/006_Sex_select_daphnia_projects/Genotyping/Coancestry.20180730/Genotype_data_coancestry_SP1_6.txt", row.names=FALSE,  col.names=FALSE,sep=" ",quote=F)

input_free = readgenotypedata ("F:/Dropbox/0_work/001_Res/0_Proj/006_Sex_select_daphnia_projects/Genotyping/Coancestry.20180730/Genotype_data_coancestry_SP1.txt")

## coancestry coefficient or the coefficient of kinship, h, between two individuals. In Wright's correlation definition, h between two individuals is simply equal to the expected F of their (hypothetical) offspring, and F can be regarded as the coancestry coefficient between the male and female gametes that unite to form an individual (From Wang  JEB 2014)

##Pop SP1-5

input_5 = readgenotypedata ("F:/Dropbox/0_work/001_Res/0_Proj/006_Sex_select_daphnia_projects/Genotyping/Coancestry.20180730/Genotype_data_coancestry_SP1_5.txt")

## Simulation SP1-5
#Simulation
# simdata_SP1_5 = familysim(input_5$freqs,1000)
# input_simdata_SP1_5= readgenotypedata(simdata_SP1_5)
# 
# output_sim_SP1_5 = coancestry (input_simdata_SP1_5$gdata, wang =1,lynchli = 1,lynchrd = 1,quellergt = 1,ritland = 1,dyadml = 1,trioml = 1, allow.inbreeding = T)
# write.table(output_sim_SP1_5$relatedness,"F:/Dropbox/0_work/001_Res/0_Proj/006_Sex_select_daphnia_projects/Genotyping/Coancestry.20180730/Results_genotype_data_coancestry_SP1-5_simu.txt", row.names=FALSE,sep=" ",quote=F)

#output_sim_SP1_5=read.table("F:/Dropbox/0_work/001_Res/0_Proj/006_Sex_select_daphnia_projects/Genotyping/Coancestry.20180730/Results_genotype_data_coancestry_SP1-5_simu.txt")
# toto = cleanuprvals(output_sim_SP1_5$relatedness,1000)
# sim_values = toto[,11]
# label1 = rep("PO",1000)
# label2 = rep("Full",1000)
# label3 = rep("Half",1000)
# label4 = rep("Unrelated",1000)
# labels=c(label1,label2,label3,label4)
# 
# sim_values = data.frame(sim_values)
# sim_values[,2]=as.factor(labels)
# colnames(sim_values)[2]="Relatedness"
# 
# with(sim_values,
# plot(Relatedness,sim_values,ylab="Relatedness value", xlab="Relatedness"))

## Analysis SP1-5

#outfile_5 = coancestry (input_5$gdata, wang =1,lynchli = 1,lynchrd = 1,quellergt = 1,ritland = 1,dyadml = 1,trioml = 1 ,allow.inbreeding = T)
#write.table(outfile_5$relatedness,"F:/Dropbox/0_work/001_Res/0_Proj/006_Sex_select_daphnia_projects/Genotyping/Coancestry.20180730/Results_genotype_data_coancestry_SP1_5.txt", row.names=FALSE,sep=" ",quote=F)

tab_rel_SP1_5 = read.table("F:/Dropbox/0_work/001_Res/0_Proj/006_Sex_select_daphnia_projects/Genotyping/Coancestry.20180730/Results_genotype_data_coancestry_SP1_5.txt",header=T)

tab_rel_SP1_5 = subset(tab_rel_SP1_5,!(group%in%c("FEFE","MAMA")))

tab_rel_SP1_5_mating = tab_rel_SP1_5[!grepl("Fr", tab_rel_SP1_5$ind1.id),] 
tab_rel_SP1_5_mating = tab_rel_SP1_5_mating[!grepl("Fr", tab_rel_SP1_5_mating$ind2.id),] 

tab_rel_SP1_5_mating$Mating_1 = LEFT(as.character(tab_rel_SP1_5_mating$ind1.id),4)
tab_rel_SP1_5_mating$Mating_1 = RIGHT(as.character(tab_rel_SP1_5_mating$Mating_1),7)
tab_rel_SP1_5_mating$Mating_2 = LEFT(as.character(tab_rel_SP1_5_mating$ind2.id),4)
tab_rel_SP1_5_mating$Mating_2 = RIGHT(as.character(tab_rel_SP1_5_mating$Mating_2),7)
tab_rel_SP1_5_mating = subset(tab_rel_SP1_5_mating,Mating_1==Mating_2)
tab_rel_SP1_5_mating$Mating_status="Mating"

tab_rel_SP1_5_random = tab_rel_SP1_5[grepl("FE_Mt", tab_rel_SP1_5$ind1.id)|grepl("FE_Mt", tab_rel_SP1_5$ind2.id),] 

tab_rel_SP1_5_random$couple_tested_1 = paste(tab_rel_SP1_5_random$ind1.id,tab_rel_SP1_5_random$ind2.id,sep="_")
tab_rel_SP1_5_random$couple_tested_2 = paste(tab_rel_SP1_5_random$ind2.id,tab_rel_SP1_5_random$ind1.id,sep="_")

list_SP1_5_random = unique(tab_rel_SP1_5_random$ind1.id)
tab_SP1_5_random = tab_rel_SP1_5_random[-c(1:nrow(tab_rel_SP1_5_random)),]

for( i in 1:length(list_SP1_5_random)){
j = list_SP1_5_random[i]
tmp=subset(tab_rel_SP1_5_random,ind1.id==j)
list_couple_tested_1 = tab_SP1_5_random$couple_tested_1
list_couple_tested_2 = tab_SP1_5_random$couple_tested_2
tab_SP1_5_random = rbind(tab_SP1_5_random,sample_n(subset(tmp,!(couple_tested_1 %in% list_couple_tested_1) | !(couple_tested_2 %in% list_couple_tested_2) ), 1))
}

tab_SP1_5_random$Mating_status="Random"

Sub_tab_SP1_5 = rbind(tab_SP1_5_random[,-c(12,13)],tab_rel_SP1_5_mating[,-c(12,13)])
Sub_tab_SP1_5$Population = "SP1-5"

# par(mfrow=c(2,4))
# plot(trioml~as.factor(Mating_status),Sub_tab_SP1_5)
# plot(wang~as.factor(Mating_status),Sub_tab_SP1_5)
# plot(lynchli~as.factor(Mating_status),Sub_tab_SP1_5)
# plot(lynchrd~as.factor(Mating_status),Sub_tab_SP1_5)
# plot(ritland~as.factor(Mating_status),Sub_tab_SP1_5)
# plot(quellergt~as.factor(Mating_status),Sub_tab_SP1_5)
# plot(dyadml~as.factor(Mating_status),Sub_tab_SP1_5)

# wilcox.test(dyadml~as.factor(Mating_status),Sub_tab_SP1_5)
# 

##Pop SP1-6

input_6 = readgenotypedata ("F:/Dropbox/0_work/001_Res/0_Proj/006_Sex_select_daphnia_projects/Genotyping/Coancestry.20180730/Genotype_data_coancestry_SP1_6.txt")

## Simulation SP1-6
#Simulation
# simdata_SP1_6 = familysim(input_6$freqs,100)
# input_simdata_SP1_6= readgenotypedata(simdata_SP1_6)
# 
# #output_sim_SP1_6 = coancestry (input_simdata_SP1_6$gdata, dyadml = 1,trioml = 1, allow.inbreeding = T)
# #write.table(output_sim_SP1_6$relatedness,"F:/Dropbox/0_work/001_Res/0_Proj/006_Sex_select_daphnia_projects/Genotyping/Coancestry.20180730/Results_genotype_data_coancestry_SP1-6_simu.txt", row.names=FALSE,sep=" ",quote=F)
# 
# output_sim_SP1_6=read.table("F:/Dropbox/0_work/001_Res/0_Proj/006_Sex_select_daphnia_projects/Genotyping/Coancestry.20180730/Results_genotype_data_coancestry_SP1-6_simu.txt")
# toto = cleanuprvals(output_sim_SP1_6$relatedness,100)
# sim_values = toto[,5]
# label1 = rep("PO",100)
# label2 = rep("Full",100)
# label3 = rep("Half",100)
# label4 = rep("Unrelated",100)
# labels=c(label1,label2,label3,label4)
# 
# sim_values = data.frame(sim_values)
# sim_values[,2]=as.factor(labels)
# colnames(sim_values)[2]="Relatedness"
# 
# with(sim_values,
# plot(Relatedness,sim_values,ylab="Relatedness value", xlab="Relatedness"))
# 
## Analysis SP1-6

#outfile_6 = coancestry (input_6$gdata, wang =1,lynchli = 1,lynchrd = 1,quellergt = 1,ritland = 1,dyadml = 1,trioml = 1 ,allow.inbreeding = T)
#write.table(outfile_6$relatedness,"F:/Dropbox/0_work/001_Res/0_Proj/006_Sex_select_daphnia_projects/Genotyping/Coancestry.20180730/Results_genotype_data_coancestry_SP1_6.txt", row.names=FALSE,sep=" ",quote=F)

tab_rel_SP1_6 = read.table("F:/Dropbox/0_work/001_Res/0_Proj/006_Sex_select_daphnia_projects/Genotyping/Coancestry.20180730/Results_genotype_data_coancestry_SP1_6.txt",header=T)

tab_rel_SP1_6 = subset(tab_rel_SP1_6,!(group%in%c("FEFE","MAMA")))

tab_rel_SP1_6_mating = tab_rel_SP1_6[!grepl("Fr", tab_rel_SP1_6$ind1.id),] 
tab_rel_SP1_6_mating = tab_rel_SP1_6_mating[!grepl("Fr", tab_rel_SP1_6_mating$ind2.id),] 

tab_rel_SP1_6_mating$Mating_1 = LEFT(as.character(tab_rel_SP1_6_mating$ind1.id),4)
tab_rel_SP1_6_mating$Mating_1 = RIGHT(as.character(tab_rel_SP1_6_mating$Mating_1),7)
tab_rel_SP1_6_mating$Mating_2 = LEFT(as.character(tab_rel_SP1_6_mating$ind2.id),4)
tab_rel_SP1_6_mating$Mating_2 = RIGHT(as.character(tab_rel_SP1_6_mating$Mating_2),7)
tab_rel_SP1_6_mating = subset(tab_rel_SP1_6_mating,Mating_1==Mating_2)
tab_rel_SP1_6_mating$Mating_status="Mating"

tab_rel_SP1_6_random = tab_rel_SP1_6[grepl("FE_Mt", tab_rel_SP1_6$ind1.id)|grepl("FE_Mt", tab_rel_SP1_6$ind2.id),] 

tab_rel_SP1_6_random$couple_tested_1 = paste(tab_rel_SP1_6_random$ind1.id,tab_rel_SP1_6_random$ind2.id,sep="_")
tab_rel_SP1_6_random$couple_tested_2 = paste(tab_rel_SP1_6_random$ind2.id,tab_rel_SP1_6_random$ind1.id,sep="_")

list_SP1_6_random = unique(tab_rel_SP1_6_random$ind1.id)
tab_SP1_6_random = tab_rel_SP1_6_random[-c(1:nrow(tab_rel_SP1_6_random)),]

for( i in 1:length(list_SP1_6_random)){
j = list_SP1_6_random[i]
tmp=subset(tab_rel_SP1_6_random,ind1.id==j)
list_couple_tested_1 = tab_SP1_6_random$couple_tested_1
list_couple_tested_2 = tab_SP1_6_random$couple_tested_2
tab_SP1_6_random = rbind(tab_SP1_6_random,sample_n(subset(tmp,!(couple_tested_1 %in% list_couple_tested_1) | !(couple_tested_2 %in% list_couple_tested_2) ), 1))
}

tab_SP1_6_random$Mating_status="Random"

Sub_tab_SP1_6 = rbind(tab_SP1_6_random[,-c(12,13)],tab_rel_SP1_6_mating[,-c(12,13)])
Sub_tab_SP1_6$Population = "SP1-6"
```

```
Sub_tab_SP1 = rbind(Sub_tab_SP1_5,Sub_tab_SP1_6)

Plot_relatedness=
  ggplot(Sub_tab_SP1,aes(x=as.factor(Mating_status),y=trioml)) +
  geom_violin(draw_quantiles = c(0.25, 0.5, 0.75),colour="darkgrey",adjust=0.5)+
  geom_dotplot(binaxis = "y", stackdir = "center",dotsize= 0.2, fill=NA,color="black", show.legend=FALSE)+
  labs(caption = "(Supplementary figure)")+
  facet_grid(.~Population,scales="free_y")+
  scale_y_continuous("Relatedness value (trioML index)")+
  scale_x_discrete("Pairs",
                   expand=c(0.1,0.1),
                   limits=c("Random","Mating"),
                   labels=c("Random","mating"))+
  stat_summary(fun.y = mean, geom = "point",colour="red",size=2) +
  theme(axis.title.x = element_text(size=Mediumfont+4),
        axis.title.y = element_text(size=Mediumfont+4),
        axis.line.x = element_line(colour="black",size=0.75),
        axis.line.y = element_line(colour="black",size=0.75),
        axis.ticks.x = element_line(size = 0.75),
        axis.ticks.y = element_line(size = 0.75),
        axis.text.x = element_text(size=Mediumfont,colour="black"),
        axis.text.y = element_text(size=Mediumfont,colour="black"),
        plot.margin = unit(Margin, "cm"),
        strip.text.y = element_text(size =Mediumfont+2, colour = "black",face="italic"),
        panel.background = element_blank())+
  guides(shape=guide_legend(ncol=1),
         fill=guide_legend(ncol=1),
         col=guide_legend(ncol=1))

grid.draw(Plot_relatedness)
```

```
# svg(filename="F:/Dropbox/0_work/001_Res/0_Proj/006_Sex_select_daphnia_projects/04_presentation/Plot_relatedness.svg",
#     width=10,
#     height=4,
#     pointsize=12)
# Plot_relatedness
# dev.off()
```

```
round(rbind(input_5$freqs$locus1,input_5$freqs$locus2,input_5$freqs$locus3,input_5$freqs$locus4),digit=3)%>%
  kable(col.names = c("Alleles","Frequencies")) %>%
    kable_styling(bootstrap_options = c("striped", "hover", "condensed"), full_width = F)%>%
  add_header_above(c("Pop SP1-5" = 2))%>%
  pack_rows("Locus A", 1, 1) %>%
  pack_rows("Locus B", 5, 5)%>%
  pack_rows("Locus C", 8, 8)%>%
  pack_rows("Locus D", 11, 11)
```

| Pop SP1-5 | |
| --- | --- |
| Alleles | Frequencies |
| **Locus A** | |
| 244 | 0.002 |
| 246 | 0.081 |
| 250 | 0.367 |
| 255 | 0.549 |
| **Locus B** | |
| 438 | 0.048 |
| 457 | 0.125 |
| 492 | 0.827 |
| **Locus C** | |
| 355 | 0.600 |
| 362 | 0.002 |
| 387 | 0.398 |
| **Locus D** | |
| 181 | 0.004 |
| 183 | 0.023 |
| 186 | 0.029 |
| 195 | 0.745 |
| 201 | 0.200 |

```
round(rbind(input_6$freqs$locus1,input_6$freqs$locus2,input_6$freqs$locus3,input_6$freqs$locus4),digit=3)%>%
  kable(col.names = c("Alleles","Frequencies")) %>%
    kable_styling(bootstrap_options = c("striped", "hover", "condensed"), full_width = F)%>%
  add_header_above(c("Pop SP1-6" = 2))%>%
  pack_rows("Locus A", 1, 1) %>%
  pack_rows("Locus B", 7, 7)%>%
  pack_rows("Locus C", 10, 10)%>%
  pack_rows("Locus D", 12, 12)
```

| Pop SP1-6 | |
| --- | --- |
| Alleles | Frequencies |
| **Locus A** | |
| 236 | 0.002 |
| 241 | 0.002 |
| 244 | 0.004 |
| 246 | 0.107 |
| 250 | 0.307 |
| 255 | 0.578 |
| **Locus B** | |
| 438 | 0.044 |
| 457 | 0.128 |
| 492 | 0.828 |
| **Locus C** | |
| 355 | 0.556 |
| 387 | 0.444 |
| **Locus D** | |
| 183 | 0.015 |
| 186 | 0.029 |
| 195 | 0.797 |
| 201 | 0.158 |

```
unlist(format(wilcox.test(trioml~as.factor(Mating_status), data=Sub_tab_SP1_5),digit=3))%>%
  kable(col.names = "Pop. SP1-5") %>%
  kable_styling(bootstrap_options = c("striped", "hover", "condensed"), full_width = F)
```

|  | Pop. SP1-5 |
| --- | --- |
| statistic | 17740 |
| parameter | NULL |
| p.value | 0.232 |
| null.value | 0 |
| alternative | two.sided |
| method | Wilcoxon rank sum test with continuity correction |
| data.name | trioml by as.factor(Mating\_status) |

```
unlist(format(wilcox.test(trioml~as.factor(Mating_status), data=Sub_tab_SP1_6),digit=3))%>%
  kable(col.names = "Pop. SP1-6") %>%
  kable_styling(bootstrap_options = c("striped", "hover", "condensed"), full_width = F)
```

|  | Pop. SP1-6 |
| --- | --- |
| statistic | 15446 |
| parameter | NULL |
| p.value | 0.892 |
| null.value | 0 |
| alternative | two.sided |
| method | Wilcoxon rank sum test with continuity correction |
| data.name | trioml by as.factor(Mating\_status) |

#### 1.5 - Effect of *H. tvaerminnensis* infection on mating

##### 1.5.1 Prevalence in studied populations

**What’s the prevalence of H. t. in natural metapopulation? (Supplementary figure)**  
We found that the prevalence of H.t. overall in summer was in average around 40%, ranging from 0 to 100% of individuals (both sexes) infected. Thus, H.t is frequent in the Finnish metapopulation at the moment of our study and in agreement with Ebert, Hottinger, and Pajunen (2001) and Lass and Ebert (2006).

```
Plot_prevalence_free = 
ggplot(sex_ratio_free,
         aes(x=Mating_status,y=Infection_rate)) +
  labs(caption = "(Supplementary figure)")+
  geom_boxplot(size =0.6,outlier.shape=NA,fill=NA)+
  geom_dotplot(alpha=0.3, binaxis = "y", stackdir = "center",dotsize=1, show.legend=FALSE)+
  scale_y_continuous("Proportion of infected",
                     limits=c(0,1),
                     breaks=(seq(0,1,by=0.2)))+
  scale_x_discrete("",
                   labels="All populations")+
  stat_summary(fun.y = mean, geom = "point",colour="red",size=2) +
  theme(axis.title.x = element_text(size=Mediumfont+4),
        axis.title.y = element_text(size=Mediumfont+4),
        axis.line.x = element_line(colour="black",size=0.75),
        axis.line.y = element_line(colour="black",size=0.75),
        axis.ticks.x = element_line(size = 0.75),
        axis.ticks.y = element_line(size = 0.75),
        axis.text.x = element_text(size=Mediumfont,colour="black"),
        axis.text.y = element_text(size=Mediumfont,colour="black"),
        plot.margin = unit(Margin, "cm"),
        legend.position = "none",
        panel.background = element_blank())

grid.draw(Plot_prevalence_free)
```

```
# svg(filename="F:/Dropbox/0_work/001_Res/0_Proj/006_Sex_select_daphnia_projects/04_presentation/Plot_prevalence_free_population.svg",
#     width=3,
#     height=3,
#     pointsize=12)
# Plot_prevalence_free
# dev.off()
```

---

##### 1.5.2 Prevalence and male proportion

**Does prevalence correlate with male proportion in natural populations? (Figure 3A)**  
We found that the prevalence of H.t. in females single does not correlate with the proportion of males in the population.  
Roth et al. (2008) suggest that infected females produce more males but, in our study, the proportion of males could be high when the prevalence was low and the other way around, could be low when the prevalence was high. This suggests that even if individual infected females produced more males, the production of males was compensated at the population level. This is completely in accordance with Booksmythe et al. (2018) showing that population of *Daphnia magna* are able to adjust the production of males depending on the current sex-ratio.

```
nb_pop = length(levels(Sexratio_infection_NaturalPop_free$Population))

mean_sexratio = mean(Sexratio_infection_NaturalPop_free$Male_ratio,na.rm=T)
Sexratio_infection_NaturalPop_free$Sex_ratio =with( Sexratio_infection_NaturalPop_free,
Male_ratio/(1-Male_ratio))

tmp=Sexratio_infection_NaturalPop_free
Sexratio_infection_NaturalPop_free$Sampling_date <- as.character(Sexratio_infection_NaturalPop_free$Sampling_date)
Sexratio_infection_NaturalPop_free$Sampling_date[grep("2003",Sexratio_infection_NaturalPop_free$Sampling_date)]="2003"
Sexratio_infection_NaturalPop_free$Sampling_date[grep("2010",Sexratio_infection_NaturalPop_free$Sampling_date)]="2010"
Sexratio_infection_NaturalPop_free$Sampling_date= as.factor(Sexratio_infection_NaturalPop_free$Sampling_date)

Plot_prevalence_free = 
  ggplot(Sexratio_infection_NaturalPop_free,
         aes(x = Prevalence_Female, y = Male_ratio)) +
  geom_point(aes(size= Population_size,color=Sampling_date),alpha=0.3,shape=19)+
  labs(caption = "(Figure 3A)")+ 
  scale_x_continuous("Prevalence in single females",
                     limits=c(0, 1),
                     breaks=c(0,0.2,0.4,0.6,0.8,1))+
  scale_y_continuous("Proportion of males",
                     limits=c(0, 1),
                     breaks=c(seq(0,1,by=0.2)))+
  scale_color_viridis_d("Sampling year",end=0.8) +
  scale_size("Sampling size (N)", breaks = c(15,25,50,100))+
  geom_hline(yintercept=1,colour=grey(0.5),linetype=3)+
  theme(aspect.ratio=1,
        axis.title.x = element_text(size=Mediumfont,colour="black"),
        axis.title.y = element_text(size=Mediumfont,colour="black"),
        axis.line.x = element_line(colour="black",size=0.75),
        axis.line.y = element_line(colour="black",size=0.75),
        axis.ticks.x = element_line(size = 0.75),
        axis.ticks.y = element_line(size = 0.75),
        axis.text.x = element_text(size=Mediumfont,colour="black"),
        axis.text.y = element_text(size=Mediumfont,colour="black"),
        plot.margin = unit(Margin, "cm"),
        strip.text.x = element_text(size =Smallfont, colour = "black",face="italic"),
        strip.text.y = element_text(size =Smallfont, colour = "black",face="italic"),
        legend.direction = "vertical", 
        legend.box = "horizontal",
        legend.position = "top",
        legend.key.height = unit(0.1, "cm"),
        legend.key.width= unit(0.1, "cm"),
        legend.title = element_text(face="italic",size=Smallfont), 
        legend.key = element_rect(colour = 'white', fill = "white", linetype='dashed'),
        legend.text = element_text(size=Smallfont,margin = margin(t = 1)),
        legend.spacing.y = unit(0.2, 'cm'),        
        legend.spacing.x = unit(0.2, 'cm'),
        legend.background = element_rect(fill=NA),
        legend.box.margin=margin(-20,-10,-20,-10),
        panel.background = element_blank())+
  guides(col = guide_legend(order=1,
                            label.position = "bottom",
                           label.vjust=-1,
                            title.position = "top", 
                            title.vjust = -1,
                           override.aes = list(size=4),
                            ncol=3),
         size=guide_legend(order=2,
                           label.position = "bottom",
                           label.vjust=1,
                           title.position = "top", 
                           title.vjust = -1,
                           ncol=4))
  
grid.draw(Plot_prevalence_free)
```

```
# svg(filename="F:/Dropbox/0_work/001_Res/0_Proj/006_Sex_select_daphnia_projects/04_presentation/Plot_prevalence_free.svg",
#     width=3,
#     height=3,
#     pointsize=12)
# Plot_prevalence_free
# dev.off()
```

```
model_Cor_full=  fitme(Sex_ratio~  Population_size +  Prevalence_Female , family=gaussian(link=identity), data=Sexratio_infection_NaturalPop_free)

model_Cor_reduced=  fitme(Sex_ratio~  Population_size +  1 , family=gaussian(link=identity), data=Sexratio_infection_NaturalPop_free)

#Chi2_LRT <- 2*(model_Cor_full$APHLs[["p_v"]]-model_Cor_reduced$APHLs[["p_v"]])
#pchisq(Chi2_LRT,df=1,lower.tail = F)

test = anova(model_Cor_full,model_Cor_reduced)

Res_Cor_Sexratio_infection= data.frame(Variable=as.character("Prevalence in females"),
                              chi2_LR=as.numeric(test$basicLRT$chi2_LR),
                              df=as.numeric(test$basicLRT$df),
                              Pvalue=as.numeric(test$basicLRT$p_value),
                              Effect_mm=as.numeric(model_Cor_full$fixef[3]))

format(Res_Cor_Sexratio_infection,digit=2)%>%
  kable(col.names = c("", "chi^2 LRT", "df", "p-value", "Effect")) %>%
  add_header_above(c("fitme(Sex_ratio~ Population_size +  Prevalence_Female , family=gaussian(link=identity))" = 5))%>%
  kable_styling(bootstrap_options = c("striped", "hover", "condensed"), full_width = F)
```

| fitme(Sex\_ratio~ Population\_size + Prevalence\_Female , family=gaussian(link=identity)) | | | | |
| --- | --- | --- | --- | --- |
|  | chi^2 LRT | df | p-value | Effect |
| Prevalence in females | 0.53 | 1 | 0.47 | 0.13 |

##### 1.5.3 Effect of sex and mating status

**Are both sexes equally infected by H. t. in natural populations? Does it depend on the mating status? (Figure 3B)**  
We found that, prevalence can be different between sexes but it depends on the mating status (binomial distribution, interaction *Sex* x *Mating status*: chi2\_LRT = 17.9, df=1, p= 0.00002). On average, the prevalence is lower in males than in females in the population (as suggested in Roth et al. (2008)) but the tendency seems reversed in mating, although this result depends strongly on the population.

```
Mating.status <- c("Single", "Mating")
names(Mating.status) <- c("Free", "Mating")

SexbiasInfectionNaturalPop$group=with(SexbiasInfectionNaturalPop,paste(Mating_status,Population,Sex,sep="_"))


paired_mean_diff <- dabest(subset(SexbiasInfectionNaturalPop,  Mating_status=="Free" & !(Population %in% c("G-45","SK-58","SP1-5"))), 
                             Sex, Infection_rate,
                             idx = c("Female","Male"),
                             paired = T,id.col=Population,
                             ci = 95, reps = 5000)

toto= as.data.frame(paired_mean_diff$result)
boot_diff_sex = unlist(toto$bootstraps)# get the bootstrap values
tata= as.data.frame(boot_diff_sex)


mean_diff = toto$difference
Boot_low = toto$pct_ci_low # The lower limits of the percentile bootstrap confidence interval.
Boot_up = toto$pct_ci_high # The upper limits of the percentile bootstrap confidence interval.

Bootstrap_plot= cbind(mean_diff,Boot_low,Boot_up)
Bootstrap_plot = as.data.frame(Bootstrap_plot)


Sexbias_infection_single_diff=
ggplot(tata,aes(x=boot_diff_sex)) +
  labs(caption = "(Figure 3Bi)")+
  geom_histogram(colour="black",fill="black",alpha=0.3,binwidth = 0.005)+
  geom_segment(data=Bootstrap_plot, aes(y=100,yend=100, x=Boot_low, xend=Boot_up),color="red") +
  geom_point(data=Bootstrap_plot, aes(y=100, x=mean_diff),color="red",size=2) +
  scale_y_continuous("bootstrap")+
  scale_x_continuous("Mean difference",
                     limits=c(-0.21,0.305),
                     breaks=c(seq(-0.20,0.3,by=0.1)))+
  theme(axis.title.x = element_text(size=Mediumfont),
        axis.title.y = element_text(size=Mediumfont),
        axis.line.x = element_line(colour="black",size=0.75),
        axis.line.y = element_line(colour="black",size=0.75),
        axis.ticks.x = element_line(size = 0.75),
        axis.ticks.y = element_line(size = 0.75),
        axis.text.x = element_text(size=Mediumfont,colour="black"),
        axis.text.y = element_text(size=Mediumfont,colour="black"),
        plot.margin = unit(Margin, "cm"),
        legend.position = "none",
        strip.text.x = element_text(size =Mediumfont, colour = "black",face="italic"),
        strip.text.y = element_text(size =Mediumfont, colour = "black",face="italic"),
        strip.background = element_rect(color="white",fill=grey(0.90)),
        panel.background = element_blank())


Sexbias_infection_single=
ggplot(subset(SexbiasInfectionNaturalPop,  Mating_status=="Free" & !(Population %in% c("G-45","SK-58","SP1-5"))), 
       aes(Sex, Infection_rate, group=Population,colour = Sex)) +
  ggtitle("Single")+
  geom_path(alpha = 0.5,size=0.7,color="black",position = position_dodge(width=0.2))+
  geom_point(shape=1,aes(size= n),position = position_dodge(width=0.2))+
  scale_size("N", breaks = c(1,10,25))+
  geom_hline(yintercept = 0.5,colour=grey(0.5),linetype=3)+  
 scale_colour_manual("",
                     guide=FALSE,
                      limits=c("Female","Male"),
                      values=c("red","blue"),
                      labels=c("Female","Male"))+
  scale_x_discrete("",
                   expand=c(0.1,0.1),
                     limits=c("Female","Male"),
                     breaks=c("Female","Male"),
                   labels=c("Female","Male"))+
  scale_y_continuous("Prevalence",
                     limits=c(0, 1.1),
                     breaks=c(0,0.2,0.4,0.6,0.8,1))+
  theme(axis.title.x = element_text(size=Mediumfont),
        axis.title.y = element_text(size=Mediumfont),
        axis.line.x = element_line(colour="black",size=0.75),
        axis.line.y = element_line(colour="black",size=0.75),
        axis.ticks.x = element_line(size = 0.75),
        axis.ticks.y = element_line(size = 0.75),
        axis.text.x = element_text(size=Mediumfont,colour="black",angle=0),
        axis.text.y = element_text(size=Mediumfont,colour="black"),
        plot.margin = unit(Margin, "cm"),
        plot.title = element_text(hjust = 0.5),
        strip.text.x = element_text(size =Mediumfont, colour = "black",face="italic"),
        strip.text.y = element_text(size =Mediumfont, colour = "black",face="italic"),
        strip.background = element_rect(color="white",fill=grey(0.90)),
        legend.direction = "horizontal", 
        legend.box = "horizontal",
        legend.position = "top",
        panel.background = element_blank())


p1= Sexbias_infection_single
p2= Sexbias_infection_single_diff

gg_Fig_Sexbias_infection_single=
as_ggplot(arrangeGrob(p1,p2, ncol = 1, nrow=2,heights=c(1,0.4)))

gg_Fig_Sexbias_infection_single
```

```
# svg(filename="F:/Dropbox/0_work/001_Res/0_Proj/006_Sex_select_daphnia_projects/04_presentation/gg_Fig_Sexbias_infection_single.svg",
#     width=3,
#     height=6,
#     pointsize=12)
# gg_Fig_Sexbias_infection_single
# dev.off()
```

```
Mating.status <- c("Single", "Mating")
names(Mating.status) <- c("Free", "Mating")

SexbiasInfectionNaturalPop$group=with(SexbiasInfectionNaturalPop,paste(Mating_status,Population,Sex,sep="_"))

paired_mean_diff <- dabest(subset(SexbiasInfectionNaturalPop, !(group%in%c("Mating_N-43_Male","Mating_N_86_Male")) & Mating_status=="Mating"), 
                             Sex, Infection_rate,
                             idx = c("Female","Male"),
                             paired = T,id.col=Population,
                             ci = 95, reps = 5000)

toto= as.data.frame(paired_mean_diff$result)
boot_diff_sex = unlist(toto$bootstraps)# get the bootstrap values
tata= as.data.frame(boot_diff_sex)


mean_diff = toto$difference
Boot_low = toto$pct_ci_low # The lower limits of the percentile bootstrap confidence interval.
Boot_up = toto$pct_ci_high # The upper limits of the percentile bootstrap confidence interval.

Bootstrap_plot= cbind(mean_diff,Boot_low,Boot_up)
Bootstrap_plot = as.data.frame(Bootstrap_plot)


Sexbias_infection_mating_diff=
ggplot(tata,aes(x=boot_diff_sex)) +
  labs(caption = "(Figure 3Bii)")+
  geom_histogram(colour="black",fill="black",alpha=0.3,binwidth = 0.005)+
  geom_segment(data=Bootstrap_plot, aes(y=50,yend=50, x=Boot_low, xend=Boot_up),color="red") +
  geom_point(data=Bootstrap_plot, aes(y=50, x=mean_diff),color="red",size=2) +
  scale_y_continuous("bootstrap")+
  scale_x_continuous("Mean difference",
                     limits=c(-0.21,0.305),
                     breaks=c(seq(-0.20,0.3,by=0.1)))+
  theme(axis.title.x = element_text(size=Mediumfont),
        axis.title.y = element_text(size=Mediumfont),
        axis.line.x = element_line(colour="black",size=0.75),
        axis.line.y = element_line(colour="black",size=0.75),
        axis.ticks.x = element_line(size = 0.75),
        axis.ticks.y = element_line(size = 0.75),
        axis.text.x = element_text(size=Mediumfont,colour="black"),
        axis.text.y = element_text(size=Mediumfont,colour="black"),
        plot.margin = unit(Margin, "cm"),
        legend.position = "none",
        strip.text.x = element_text(size =Mediumfont, colour = "black",face="italic"),
        strip.text.y = element_text(size =Mediumfont, colour = "black",face="italic"),
        strip.background = element_rect(color="white",fill=grey(0.90)),
        panel.background = element_blank())


Sexbias_infection_mating=
ggplot(subset(SexbiasInfectionNaturalPop, !(group%in%c("Mating_N-43_Male","Mating_N_86_Male")) & Mating_status=="Mating"), 
       aes(Sex, Infection_rate, group=Population,colour = Sex)) +
  geom_path(alpha = 0.5,size=1,color="black",position = position_dodge(width=0.2))+
  geom_point(shape=1,aes(size= n),position = position_dodge(width=0.2))+
  ggtitle("Mating")+
  scale_size("N", breaks = c(1,10,25,50))+
  geom_hline(yintercept = 0.5,colour=grey(0.5),linetype=3)+  
 scale_colour_manual("",
                     guide=FALSE,
                      limits=c("Female","Male"),
                      values=c("red","blue"),
                      labels=c("Females", "Males"))+
  scale_x_discrete("",
                   expand=c(0.1,0.1),
                     limits=c("Female","Male"),
                     breaks=c("Female","Male"),
                   labels=c("Females", "Males"))+
  scale_y_continuous("Prevalence",
                     limits=c(0, 1.1),
                     breaks=c(0,0.2,0.4,0.6,0.8,1))+
#  geom_text(data = Results_SexbiasInfection, aes(label = Test,x= 1.5, y = 1.1),colour="black",size=4)+
  theme(axis.title.x = element_text(size=Mediumfont),
        axis.title.y = element_text(size=Mediumfont),
        axis.line.x = element_line(colour="black",size=0.75),
        axis.line.y = element_line(colour="black",size=0.75),
        axis.ticks.x = element_line(size = 0.75),
        axis.ticks.y = element_line(size = 0.75),
        axis.text.x = element_text(size=Mediumfont,colour="black",angle=0),
        axis.text.y = element_text(size=Mediumfont,colour="black"),
        plot.margin = unit(Margin, "cm"),
        plot.title = element_text(hjust = 0.5),
        strip.text.x = element_text(size =Mediumfont, colour = "black",face="italic"),
        strip.text.y = element_text(size =Mediumfont, colour = "black",face="italic"),
        strip.background = element_rect(color="white",fill=grey(0.90)),
        legend.direction = "horizontal", 
        legend.box = "horizontal",
        legend.position = "top",
        panel.background = element_blank())

p1= Sexbias_infection_mating
p2= Sexbias_infection_mating_diff

gg_Fig_Sexbias_infection_mating=
as_ggplot(arrangeGrob(p1,p2, ncol = 1, nrow=2,heights=c(1,0.4)))

gg_Fig_Sexbias_infection_mating
```

```
# svg(filename="F:/Dropbox/0_work/001_Res/0_Proj/006_Sex_select_daphnia_projects/04_presentation/gg_Fig_Sexbias_infection_mating.svg",
#     width=3,
#     height=6,
#     pointsize=12)
# gg_Fig_Sexbias_infection_mating
# dev.off()
```

```
data_temp = subset(all_data, !is.na(Infectious_status) & Mating_status=="Free")

model_prevalence_Sex_single_full = fitme(Infectious_status~ Sex + (1|Population),family=binomial(link="logit"), data=data_temp)

model_prevalence_Sex_single_reduced = fitme(Infectious_status~ 1 + (1|Population),family=binomial(link="logit"), data=data_temp)

test=anova(model_prevalence_Sex_single_full,model_prevalence_Sex_single_reduced) 


Res_Prevalence_sex_single= data.frame(Variable=as.character("Dif. female to male"),
                              chi2_LR=as.numeric(test$basicLRT$chi2_LR),
                              df=as.numeric(test$basicLRT$df),
                              Pvalue=as.numeric(test$basicLRT$p_value),
                              Effect=as.numeric(exp(model_prevalence_Sex_single_full$fixef[2])))


format(Res_Prevalence_sex_single,digit=2)%>%
  kable(col.names = c("", "chi^2 LRT", "df", "p-value", "Odd ratio")) %>%
  add_header_above(c("fitme(Infectious_status~ Sex + (1|Population),\nfamily=binomial(link=logit)" = 5))%>%
  kable_styling(bootstrap_options = c("striped", "hover", "condensed"), full_width = F)
```

| fitme(Infectious\_status~ Sex + (1\|Population), family=binomial(link=logit) | | | | |
| --- | --- | --- | --- | --- |
|  | chi^2 LRT | df | p-value | Odd ratio |
| Dif. female to male | 15 | 1 | 0.00012 | 0.64 |

```
data_temp = subset(all_data, !is.na(Infectious_status) & Mating_status=="Mating")

model_prevalence_Sex_Mating_full = fitme(Infectious_status~ Sex + (1|Population),family=binomial(link="logit"), data=data_temp)

model_prevalence_Sex_Mating_reduced = fitme(Infectious_status~ 1 + (1|Population),family=binomial(link="logit"), data=data_temp)

test=anova(model_prevalence_Sex_Mating_full,model_prevalence_Sex_Mating_reduced) 

Res_Prevalence_sex_Mating= data.frame(Variable=as.character("Dif. female to male"),
                              chi2_LR=as.numeric(test$basicLRT$chi2_LR),
                              df=as.numeric(test$basicLRT$df),
                              Pvalue=as.numeric(test$basicLRT$p_value),
                              Effect=as.numeric(exp(model_prevalence_Sex_Mating_full$fixef[2])))


format(Res_Prevalence_sex_Mating,digit=2)%>%
  kable(col.names = c("", "chi^2 LRT", "df", "p-value", "Odd ratio")) %>%
  add_header_above(c("fitme(Infectious_status~ Sex + (1|Population),\nfamily=binomial(link=logit)" = 5))%>%
  kable_styling(bootstrap_options = c("striped", "hover", "condensed"), full_width = F)
```

| fitme(Infectious\_status~ Sex + (1\|Population), family=binomial(link=logit) | | | | |
| --- | --- | --- | --- | --- |
|  | chi^2 LRT | df | p-value | Odd ratio |
| Dif. female to male | 4.7 | 1 | 0.03 | 1.4 |

**Are individuals in mating less infected than those single? (Figure 3C)**

```
tmp = subset(SexbiasInfectionNaturalPop, Mating_status=="Mating" & Sex=="Male")
tmp$Population =factor(tmp$Population)
tmp2 = subset(SexbiasInfectionNaturalPop, Mating_status=="Free" & Sex=="Male")
tmp2$Population =factor(tmp2$Population)

list_pop_male = levels(tmp2$Population)[which(levels(tmp2$Population) %in% levels(tmp$Population))]

Mating.status <- c("Single", "Mating")
names(Mating.status) <- c("Free", "Mating")

SexbiasInfectionNaturalPop$group=with(SexbiasInfectionNaturalPop,paste(Mating_status,Population,Sex,sep="_"))

Mating_bias_infection_Male= 
  ggplot(subset(SexbiasInfectionNaturalPop, Population %in% list_pop_male & Sex=="Male"), 
       aes(Mating_status, Infection_rate, group=Population)) +
  ggtitle("Males")+
  geom_path(alpha = 0.5,size=1,color="black",position = position_dodge(width=0.2))+
  geom_point(shape=1,color = "blue",aes(size= n),position = position_dodge(width=0.2))+
  scale_size("N", breaks = c(1,10,25,50))+
  geom_hline(yintercept = 0.5,colour=grey(0.5),linetype=3)+  
  scale_x_discrete("",
                   expand=c(0.1,0.1),
                     limits=c("Free","Mating"),
                     breaks=c("Free","Mating"),
                   labels=c("Single", "Mating"))+
  scale_y_continuous("Prevalence in males",
                     limits=c(0, 1.1),
                     breaks=c(0,0.2,0.4,0.6,0.8,1))+
  theme(axis.title.x = element_text(size=Mediumfont),
        axis.title.y = element_text(size=Mediumfont),
        axis.line.x = element_line(colour="black",size=0.75),
        axis.line.y = element_line(colour="black",size=0.75),
        axis.ticks.x = element_line(size = 0.75),
        axis.ticks.y = element_line(size = 0.75),
        axis.text.x = element_text(size=Mediumfont,colour="black",angle=0),
        axis.text.y = element_text(size=Mediumfont,colour="black"),
        plot.margin = unit(Margin, "cm"),
        plot.title = element_text(hjust = 0.5),
        strip.text.x = element_text(size =Mediumfont, colour = "black",face="italic"),
        strip.text.y = element_text(size =Mediumfont, colour = "black",face="italic"),
        strip.background = element_rect(color="white",fill=grey(0.90)),
        legend.direction = "horizontal", 
        legend.box = "horizontal",
        legend.position = "top",
        panel.background = element_blank())


paired_mean_diff <- dabest(subset(SexbiasInfectionNaturalPop, Population %in% list_pop_male & Sex=="Male"), 
                             Mating_status, Infection_rate,
                             idx = c("Free","Mating"),
                             paired = T,id.col=Population,
                             ci = 95, reps = 5000)

toto= as.data.frame(paired_mean_diff$result)
boot_diff_mating = unlist(toto$bootstraps)# get the bootstrap values
tata= as.data.frame(boot_diff_mating)


mean_diff = toto$difference
Boot_low = toto$pct_ci_low # The lower limits of the percentile bootstrap confidence interval.
Boot_up = toto$pct_ci_high # The upper limits of the percentile bootstrap confidence interval.

Bootstrap_plot= cbind(mean_diff,Boot_low,Boot_up)
Bootstrap_plot = as.data.frame(Bootstrap_plot)


Sexbias_infection_Male_diff=
ggplot(tata,aes(x=boot_diff_mating)) +
  geom_histogram(colour="black",fill="black",alpha=0.3,binwidth = 0.005)+
  geom_segment(data=Bootstrap_plot, aes(y=100,yend=100, x=Boot_low, xend=Boot_up),color="red") +
  geom_point(data=Bootstrap_plot, aes(y=100, x=mean_diff),color="red",size=2) +
  labs(caption = "(Figure 3Ci)")+
  scale_y_continuous("bootstrap")+
  scale_x_continuous("Mean difference",
                     limits=c(-0.21,0.305),
                     breaks=c(seq(-0.20,0.3,by=0.1)))+
  theme(axis.title.x = element_text(size=Mediumfont),
        axis.title.y = element_text(size=Mediumfont),
        axis.line.x = element_line(colour="black",size=0.75),
        axis.line.y = element_line(colour="black",size=0.75),
        axis.ticks.x = element_line(size = 0.75),
        axis.ticks.y = element_line(size = 0.75),
        axis.text.x = element_text(size=Mediumfont,colour="black"),
        axis.text.y = element_text(size=Mediumfont,colour="black"),
        plot.margin = unit(Margin, "cm"),
        legend.position = "none",
        strip.text.x = element_text(size =Mediumfont, colour = "black",face="italic"),
        strip.text.y = element_text(size =Mediumfont, colour = "black",face="italic"),
        strip.background = element_rect(color="white",fill=grey(0.90)),
        panel.background = element_blank())


p1= Mating_bias_infection_Male
p2= Sexbias_infection_Male_diff

gg_Fig_Mating_bias_infection_Male=
as_ggplot(arrangeGrob(p1,p2, ncol = 1, nrow=2,heights=c(1,0.4)))

gg_Fig_Mating_bias_infection_Male
```

```
# svg(filename="F:/Dropbox/0_work/001_Res/0_Proj/006_Sex_select_daphnia_projects/04_presentation/gg_Fig_Mating_bias_infection_Male.svg",
#     width=3,
#     height=6,
#     pointsize=12)
# gg_Fig_Mating_bias_infection_Male
# dev.off()
```

```
tmp = subset(SexbiasInfectionNaturalPop, Mating_status=="Mating" & Sex=="Female")
tmp$Population =factor(tmp$Population)
tmp2 = subset(SexbiasInfectionNaturalPop, Mating_status=="Free" & Sex=="Female")
tmp2$Population =factor(tmp2$Population)

list_pop_female = levels(tmp2$Population)[which(levels(tmp2$Population) %in% levels(tmp$Population))]

Mating.status <- c("Single", "Mating")
names(Mating.status) <- c("Free", "Mating")

SexbiasInfectionNaturalPop$group=with(SexbiasInfectionNaturalPop,paste(Mating_status,Population,Sex,sep="_"))

Mating_bias_infection_Female= 
  ggplot(subset(SexbiasInfectionNaturalPop, Population %in% list_pop_female & Sex=="Female"), 
       aes(Mating_status, Infection_rate, group=Population)) +
  ggtitle("Females")+
  geom_path(alpha = 0.5,size=1,color="black",position = position_dodge(width=0.2))+
  geom_point(shape=1,color = "red",aes(size= n),position = position_dodge(width=0.2))+
  scale_size("N", breaks = c(1,5,10,25))+
  geom_hline(yintercept = 0.5,colour=grey(0.5),linetype=3)+  
  scale_x_discrete("",
                   expand=c(0.1,0.1),
                     limits=c("Free","Mating"),
                     breaks=c("Free","Mating"),
                   labels=c("Single", "Mating"))+
  scale_y_continuous("Prevalence in females",
                     limits=c(0, 1.1),
                     breaks=c(0,0.2,0.4,0.6,0.8,1))+
  theme(axis.title.x = element_text(size=Mediumfont),
        axis.title.y = element_text(size=Mediumfont),
        axis.line.x = element_line(colour="black",size=0.75),
        axis.line.y = element_line(colour="black",size=0.75),
        axis.ticks.x = element_line(size = 0.75),
        axis.ticks.y = element_line(size = 0.75),
        axis.text.x = element_text(size=Mediumfont,colour="black",angle=0),
        axis.text.y = element_text(size=Mediumfont,colour="black"),
        plot.margin = unit(Margin, "cm"),
        plot.title = element_text(hjust = 0.5),
        strip.text.x = element_text(size =Mediumfont, colour = "black",face="italic"),
        strip.text.y = element_text(size =Mediumfont, colour = "black",face="italic"),
        strip.background = element_rect(color="white",fill=grey(0.90)),
        legend.direction = "horizontal", 
        legend.box = "horizontal",
        legend.position = "top",
        panel.background = element_blank())

paired_mean_diff <- dabest(subset(SexbiasInfectionNaturalPop, Population %in% list_pop_female & Sex=="Female"), 
                             Mating_status, Infection_rate,
                             idx = c("Free","Mating"),
                             paired = T,id.col=Population,
                             ci = 95, reps = 5000)

toto= as.data.frame(paired_mean_diff$result)
boot_diff_mating = unlist(toto$bootstraps)# get the bootstrap values
tata= as.data.frame(boot_diff_mating)


mean_diff = toto$difference
Boot_low = toto$pct_ci_low # The lower limits of the percentile bootstrap confidence interval.
Boot_up = toto$pct_ci_high # The upper limits of the percentile bootstrap confidence interval.

Bootstrap_plot= cbind(mean_diff,Boot_low,Boot_up)
Bootstrap_plot = as.data.frame(Bootstrap_plot)


Sexbias_infection_Female_diff=
ggplot(tata,aes(x=boot_diff_mating)) +
  geom_histogram(colour="black",fill="black",alpha=0.3,binwidth = 0.005)+
  geom_segment(data=Bootstrap_plot, aes(y=100,yend=100, x=Boot_low, xend=Boot_up),color="red") +
  geom_point(data=Bootstrap_plot, aes(y=100, x=mean_diff),color="red",size=2) +
  labs(caption = "(Figure 3Cii)")+
  scale_y_continuous("bootstrap")+
  scale_x_continuous("Mean difference",
                     limits=c(-0.21,0.305),
                     breaks=c(seq(-0.20,0.3,by=0.1)))+
  theme(axis.title.x = element_text(size=Mediumfont),
        axis.title.y = element_text(size=Mediumfont),
        axis.line.x = element_line(colour="black",size=0.75),
        axis.line.y = element_line(colour="black",size=0.75),
        axis.ticks.x = element_line(size = 0.75),
        axis.ticks.y = element_line(size = 0.75),
        axis.text.x = element_text(size=Mediumfont,colour="black"),
        axis.text.y = element_text(size=Mediumfont,colour="black"),
        plot.margin = unit(Margin, "cm"),
        legend.position = "none",
        strip.text.x = element_text(size =Mediumfont, colour = "black",face="italic"),
        strip.text.y = element_text(size =Mediumfont, colour = "black",face="italic"),
        strip.background = element_rect(color="white",fill=grey(0.90)),
        panel.background = element_blank())


p1= Mating_bias_infection_Female
p2= Sexbias_infection_Female_diff

gg_Fig_Mating_bias_infection_Female=
as_ggplot(arrangeGrob(p1,p2, ncol = 1, nrow=2,heights=c(1,0.4)))

gg_Fig_Mating_bias_infection_Female
```

```
# svg(filename="F:/Dropbox/0_work/001_Res/0_Proj/006_Sex_select_daphnia_projects/04_presentation/gg_Fig_Mating_bias_infection_Female.svg",
#     width=3,
#     height=6,
#     pointsize=12)
# gg_Fig_Mating_bias_infection_Female
# dev.off()
```

```
data_temp = subset(all_data, !is.na(Infectious_status)& !is.na(Mating_status) & Sex=="Male")

model_prevalence_MatingStat_Male_full = fitme(Infectious_status ~ Mating_status + (1|Population), family=binomial(link="logit"), rand.family=gaussian(link="identity"), data=data_temp)

model_prevalence_MatingStat_Male_reduced = fitme(Infectious_status ~ 1 + (1|Population), family=binomial(link="logit"), rand.family=gaussian(link="identity"), data=data_temp)

test=anova(model_prevalence_MatingStat_Male_full,model_prevalence_MatingStat_Male_reduced) 


Res_Prevalence_sex_single= data.frame(Variable=as.character("Dif. Single to Mating"),
                              chi2_LR=as.numeric(test$basicLRT$chi2_LR),
                              df=as.numeric(test$basicLRT$df),
                              Pvalue=as.numeric(test$basicLRT$p_value),
                              Effect=as.numeric(exp(model_prevalence_MatingStat_Male_full$fixef[2])))


format(Res_Prevalence_sex_single,digit=2)%>%
  kable(col.names = c("", "chi^2 LRT", "df", "p-value", "Odd ratio")) %>%
  add_header_above(c("fitme(Infectious_status~ Mating status + (1|Population),\nfamily=binomial(link=logit), rand.family=gaussian(link=identity)" = 5))%>%
  kable_styling(bootstrap_options = c("striped", "hover", "condensed"), full_width = F)
```

| fitme(Infectious\_status~ Mating status + (1\|Population), family=binomial(link=logit), rand.family=gaussian(link=identity) | | | | |
| --- | --- | --- | --- | --- |
|  | chi^2 LRT | df | p-value | Odd ratio |
| Dif. Single to Mating | 17 | 1 | 4.8e-05 | 1.8 |

```
data_temp = subset(all_data, !is.na(Infectious_status) & !is.na(Mating_status) & Sex=="Female")

model_prevalence_MatingStat_Female_full = fitme(Infectious_status ~ Mating_status + (1|Population), family=binomial(link="logit"), data=data_temp)

model_prevalence_MatingStat_Female_reduced = fitme(Infectious_status ~ 1 + (1|Population), family=binomial(link="logit"), data=data_temp)

test=anova(model_prevalence_MatingStat_Female_full,model_prevalence_MatingStat_Female_reduced) 


Res_Prevalence_sex_single= data.frame(Variable=as.character("Dif. Single to Mating"),
                              chi2_LR=as.numeric(test$basicLRT$chi2_LR),
                              df=as.numeric(test$basicLRT$df),
                              Pvalue=as.numeric(test$basicLRT$p_value),
                              Effect=as.numeric(exp(model_prevalence_MatingStat_Female_full$fixef[2])))


format(Res_Prevalence_sex_single,digit=2)%>%
  kable(col.names = c("", "chi^2 LRT", "df", "p-value", "Odd ratio")) %>%
  add_header_above(c("fitme(Infectious_status~ Mating status + (1|Population),\nfamily=binomial(link=logit)" = 5))%>%
  kable_styling(bootstrap_options = c("striped", "hover", "condensed"), full_width = F)
```

| fitme(Infectious\_status~ Mating status + (1\|Population), family=binomial(link=logit) | | | | |
| --- | --- | --- | --- | --- |
|  | chi^2 LRT | df | p-value | Odd ratio |
| Dif. Single to Mating | 0.49 | 1 | 0.49 | 0.89 |

##### 1.5.4 Infection-related assortative mating

**Is there an assortative mating with respect to infection status? (Figure 3D)**

```
Data_mating_infection = all_data[,c(1:9)]
Data_mating_infection = subset(Data_mating_infection,Mating_status=="Mating" & !is.na(Nbr_of_males) & !is.na(Infectious_status))

list_all_IDmating = unique(Data_mating_infection$ID_mating)

tab_infected_males = data.frame(ID_mating=character(),
                  Nbr_infected_males= numeric(),
                  stringsAsFactors=FALSE)

for (i in 1:length(list_all_IDmating) ){
  IDmating = as.character(list_all_IDmating[i])
  tab_infected_males[i,1] = IDmating
  tab_infected_males[i,2] = with(subset(Data_mating_infection,ID_mating==IDmating & Sex=="Male"),
                     sum(Infectious_status))
}


Data_mating_infection$Nbr_infected_males = tab_infected_males[match(Data_mating_infection$ID_mating,tab_infected_males$ID_mating),2]

Data_mating_infection$Nbr_uninfected_males = Data_mating_infection$Nbr_of_males - Data_mating_infection$Nbr_infected_males

Data_mating_infection$Prop_infected = Data_mating_infection$Nbr_infected_males/Data_mating_infection$Nbr_of_males


Av_inf_pop=
  subset(Data_mating_infection,Sex=="Female")%>%
  group_by(Population,Infectious_status)%>%
  summarise(mean_prop = mean(Prop_infected),n=n())

Av_inf_pop = as.data.frame(Av_inf_pop)
Av_inf_pop$Population = factor(Av_inf_pop$Population) 

Av_inf_pop = subset(Av_inf_pop,!(Population%in%c("N-43","N_86","BR1_39"))) #because one condition is missing and Br1_39 is totally infected.

Plot_assortative_mating_inf=
  ggplot(Av_inf_pop, aes(x=as.factor(Infectious_status), y= mean_prop, group=Population)) +
  labs(caption = "(Figure 3D)")+
  geom_point(aes(size=n),shape=19,alpha=0.3,position= position_dodge(width=0.1))+
  geom_path(size=0.5,position= position_dodge(width=0.1))+
  scale_size("N females", breaks = c(1,5,10,15,20,40))+
  scale_x_discrete("Infection status female",
                   expand=c(0.1,0.1),
                   limits=c("0","1"),
                   labels=c("Uninfected", "Infected"))+
  scale_y_continuous("Prevalence in males",
                     limits=c(0, 1),
                     breaks=c(c(seq(0,1,by=0.25))))+  
  theme(
    axis.title.x = element_text(size=Mediumfont+6),
    axis.title.y = element_text(size=Mediumfont++6),
    axis.line.x = element_line(colour="black",size=0.75),
    axis.line.y = element_line(colour="black",size=0.75),
    axis.ticks.x = element_line(size = 0.75),
    axis.ticks.y = element_line(size = 0.75),
    axis.text.x = element_text(size=Mediumfont+4,colour="black"),
    axis.text.y = element_text(size=Mediumfont+4,colour="black"),
    strip.background = element_rect(color="white",fill=grey(0.90)),
        legend.direction = "horizontal", 
        legend.box = "horizontal",
    legend.position = "top",
    legend.key.height = unit(0.4, "cm"),
    legend.key.width= unit(0.3, "cm"),
    legend.title = element_text(face="italic",size=Mediumfont+2),
    legend.key = element_rect(colour = 'white', fill = "white", linetype='dashed'),
    legend.text = element_text(size=Mediumfont+2),
    legend.background = element_rect(fill=NA),
    panel.background = element_blank())


paired_mean_diff <- dabest(Av_inf_pop, 
                            Infectious_status, mean_prop,
                             idx = c("0","1"),
                             paired = T,id.col=Population,
                             ci = 95, reps = 5000)

toto= as.data.frame(paired_mean_diff$result)
boot_diff_assort_parasite = unlist(toto$bootstraps)# get the bootstrap values
tata= as.data.frame(boot_diff_assort_parasite)


mean_diff = toto$difference
Boot_low = toto$pct_ci_low # The lower limits of the percentile bootstrap confidence interval.
Boot_up = toto$pct_ci_high # The upper limits of the percentile bootstrap confidence interval.

Bootstrap_plot= cbind(mean_diff,Boot_low,Boot_up)
Bootstrap_plot = as.data.frame(Bootstrap_plot)


Plot_assortative_mating_inf_diff=
ggplot(tata,aes(x=boot_diff_assort_parasite)) +
  geom_histogram(colour="black",fill="black",alpha=0.3,binwidth = 0.005)+
  geom_segment(data=Bootstrap_plot, aes(y=100,yend=100, x=Boot_low, xend=Boot_up),color="red") +
  geom_point(data=Bootstrap_plot, aes(y=100, x=mean_diff),color="red",size=2) +
  geom_vline(xintercept = 0,colour="red",linetype=3,size=1)+
  annotate("text",label="*",size=5,x=1.5,y=1.05)+
  scale_y_continuous(" ")+
  scale_x_continuous("Mean differences\n(Inf. - Uninf.)",
                     limits=c(-0.3,0.31),
                     breaks=c(seq(-0.3,0.3,by=0.15)))+
  theme(axis.title.x = element_text(size=Mediumfont),
        axis.title.y = element_text(size=Mediumfont),
        axis.line.x = element_line(colour="black",size=0.75),
        axis.line.y = element_line(colour="black",size=0.75),
        axis.ticks.x = element_line(size = 0.75),
        axis.ticks.y = element_line(size = 0.75),
        axis.text.x = element_text(size=Mediumfont,colour="black"),
        axis.text.y = element_text(size=Mediumfont,colour="black"),
        plot.margin = unit(c(0,0.5,0,0), "cm"),
        legend.position = "none",
        strip.text.x = element_text(size =Mediumfont, colour = "black",face="italic"),
        strip.text.y = element_text(size =Mediumfont, colour = "black",face="italic"),
        strip.background = element_rect(color="white",fill=grey(0.90)),
        panel.background = element_blank())


p1= Plot_assortative_mating_inf
p2= Plot_assortative_mating_inf_diff

gg_Fig_Plot_assortative_mating_inf=
as_ggplot(arrangeGrob(p1,p2, ncol = 1, nrow=2,heights=c(1,0.4)))


grid.draw(gg_Fig_Plot_assortative_mating_inf)
```

```
# svg(filename="F:/Dropbox/0_work/001_Res/0_Proj/006_Sex_select_daphnia_projects/04_presentation/Plot_assortative_mating_inf.svg",
#     width=5,
#     height=5,
#     pointsize=12)
# Plot_assortative_mating_inf
# dev.off()
```

```
model_assort_inf_full = fitme(cbind(Nbr_infected_males,Nbr_uninfected_males)~ Infectious_status +(1|Population/ID_mating),family=binomial(link=logit), subset(Data_mating_infection,Sex=="Female" & !is.na(Infectious_status) & !(Population%in%c("N-43","N_86","BR1_39"))))

model_assort_inf_reduced = fitme(cbind(Nbr_infected_males,Nbr_uninfected_males)~ 1 +(1|Population/ID_mating),family=binomial(link=logit), subset(Data_mating_infection,Sex=="Female" & !is.na(Infectious_status) & !(Population%in%c("N-43","N_86","BR1_39"))))

test= anova(model_assort_inf_full,model_assort_inf_reduced)

Res_assortMating_infection= data.frame(Variable=as.character("Infection status females"),
                              df=as.numeric(test$basicLRT$df),
                              chi2_LR=as.numeric(test$basicLRT$chi2_LR),
                              Pvalue=as.numeric(test$basicLRT$p_value),
                              Effect=as.numeric(exp(model_assort_inf_full$fixef[2])))


format(Res_assortMating_infection,digit=2)%>%
  kable(col.names = c("", "df", "chi^2 LRT", "p-value", "Odd ratio")) %>%
  add_header_above(c("fitme(cbind(Nbr_inf_M,Nbr_uninf_M)~ Inf_stat_F +(1|Pop/IDmating),family= binomial), rand.family= gaussian (link= identity)" = 5))%>%
  kable_styling(bootstrap_options = c("striped", "hover", "condensed"), full_width = F)
```

| fitme(cbind(Nbr\_inf\_M,Nbr\_uninf\_M)~ Inf\_stat\_F +(1\|Pop/IDmating),family= binomial), rand.family= gaussian (link= identity) | | | | |
| --- | --- | --- | --- | --- |
|  | df | chi^2 LRT | p-value | Odd ratio |
| Infection status females | 1 | 5.2 | 0.023 | 1.8 |

### 2 - Mating behavior

#### 2.1 - Time before the male detached

**How long does the sexual process last? (Figure 4)**

Polyandrous matings lasted longer than monandrous matings due to the second male remaining attached for longer (Wilcoxon rank sum test: W= 4192, p=0.0002). Monandrous matings lasted as long as the first detachment in polyandrous matings (Wilcoxon rank sum test: W= 3676, p=0.99).

```
Name=expression(paste("Time before\ndetachment (",log[2]," (min))",sep=""))
data_time_attached = subset(all_data, !is.na(Time_male_attached_min))

data_time_attached = data_time_attached[,c("ID_mating","Nbr_of_males","Position_detached","Time_male_attached_min")]
data_time_attached$X = as.factor(ifelse(data_time_attached$Nbr_of_males==1,"Single",
                              ifelse(data_time_attached$Nbr_of_males==2 & data_time_attached$Position_detached==1, "First",
                              ifelse(data_time_attached$Nbr_of_males==2 & data_time_attached$Position_detached==2, "Second","First"))))


means_time_mating <- aggregate(Time_male_attached_min ~  X, data_time_attached, mean)
se <- function(x) sqrt(var(x)/length(x))
se_time_mating <- aggregate(Time_male_attached_min ~  X, data_time_attached, se)
colnames(means_time_mating)[2]="Time_male_attached_min"
means_time_mating$Log_Time_male_attached_min =  round(log2(means_time_mating$`Time_male_attached_min`),2)
means_time_mating$Time_male_attached_min_text = paste(round(means_time_mating$Time_male_attached_min,0), " min",sep="")

Plot_length_mating_single_vs_double = 
  ggplot(data_time_attached,
         aes(x=X,y=Time_male_attached_min)) +
  labs(caption = "(Figure 4)")+
  geom_boxplot(size =0.6,outlier.shape=NA)+
  geom_dotplot(alpha=0.3, binaxis = "y", stackdir = "center",binwidth = 0.03, show.legend=FALSE)+
  scale_y_log10("Time before detachment (min)",
               limits=c(1,255),
               breaks=c(1,5,10,20,30,60,120,240)) +
  scale_x_discrete("",
                   limits=c("Single","First","Second"),
                   labels=c("Monandrous","First","Second"))+
  geom_point(data = means_time_mating, aes(x = X, y= Time_male_attached_min),colour="red",size=2)+
  geom_text(data = means_time_mating, aes(label = Time_male_attached_min_text, y =I(2^Log_Time_male_attached_min+7)), colour="red",size=4)+
  annotate("text", x=2, y=252, label= "ns",size=3.5, fontface = "bold") +
  annotate("text", x=3, y=252, label= "***",size=4, fontface = "bold") +
  theme(axis.title.x = element_text(size=Mediumfont+4),
        axis.title.y = element_text(size=Mediumfont+4),
        axis.line.x = element_line(colour="black",size=0.75),
        axis.line.y = element_line(colour="black",size=0.75),
        axis.ticks.x = element_line(size = 0.75),
        axis.ticks.y = element_line(size = 0.75),
        axis.text.x = element_text(size=Mediumfont+4,colour="black"),
        axis.text.y = element_text(size=Mediumfont+4,colour="black"),
        plot.margin = unit(Margin, "cm"),
        legend.position = "none",
        panel.background = element_blank())

grid.draw(Plot_length_mating_single_vs_double)
```

```
# svg(filename="F:/Dropbox/0_work/001_Res/0_Proj/006_Sex_select_daphnia_projects/04_presentation/Plot_length_mating_single_vs_double.svg",
#     width=5,
#     height=4,
#     pointsize=12)
# Plot_length_mating_single_vs_double
# dev.off()
```

```
unlist(format(wilcox.test(Time_male_attached_min~X, data=subset(data_time_attached,X!="First")),digit=3))%>%
  kable(col.names = "Monandrous vs Second") %>%
  kable_styling(bootstrap_options = c("striped", "hover", "condensed"), full_width = F)
```

|  | Monandrous vs Second |
| --- | --- |
| statistic | 4192 |
| parameter | NULL |
| p.value | 0.00021 |
| null.value | 0 |
| alternative | two.sided |
| method | Wilcoxon rank sum test with continuity correction |
| data.name | Time\_male\_attached\_min by X |

```
unlist(format(wilcox.test(Time_male_attached_min~X, data=subset(data_time_attached,X!="Second")),digit=3))%>%
  kable(col.names = "Monandrous vs First") %>%
  kable_styling(bootstrap_options = c("striped", "hover", "condensed"), full_width = F)
```

|  | Monandrous vs First |
| --- | --- |
| statistic | 3676 |
| parameter | NULL |
| p.value | 0.99 |
| null.value | 0 |
| alternative | two.sided |
| method | Wilcoxon rank sum test with continuity correction |
| data.name | Time\_male\_attached\_min by X |

#### 2.2 - Egg laying behavior

**When are females laying in their ephippium? (Figure 3)**

```
time_laying= subset(all_data,!is.na(Time_of_laying_after_last_male_min)&Time_of_laying_after_last_male_min!=""&Time_of_laying_after_last_male_min!="na",select=Time_of_laying_after_last_male_min)

time_laying$Time_of_laying_after_last_male_min = gsub("<","",time_laying$Time_of_laying_after_last_male_min)

time_laying$Time_of_laying_after_last_male_min = gsub(">","",time_laying$Time_of_laying_after_last_male_min)

time_laying$Time_of_laying_after_last_male_min = gsub("\\*","",time_laying$Time_of_laying_after_last_male_min)
time_laying$Female="Female"

time_laying$Time_of_laying_after_last_male_min = as.numeric(time_laying$Time_of_laying_after_last_male_min)


median_laying = median(time_laying$Time_of_laying_after_last_male_min)

Names= expression(paste("Time before laying (",log[10]," (min))",sep=""))

Distribution_time_laying=
  ggplot(time_laying,aes(x=as.numeric(time_laying$Time_of_laying_after_last_male_min))) +
labs(caption = "(Figure 4)")+
  geom_histogram(colour="black",fill="black",alpha=0.3,binwidth = 0.2)+
      scale_y_continuous("Count",
                         limits=c(0,23),
                         breaks=(seq(0,20,by=5)))+
   scale_x_log10("Time before detachment (min)",
                         breaks=c(1,2,3,5,10,20,60,120)) +
  geom_vline(aes(xintercept=median_laying),
            color="red", linetype="dashed", size=0.6)+
 theme(axis.title.x = element_text(size=Mediumfont),
        axis.title.y = element_text(size=Mediumfont),
        axis.line.x = element_line(colour="black",size=0.75),
        axis.line.y = element_line(colour="black",size=0.75),
        axis.ticks.x = element_line(size = 0.75),
        axis.ticks.y = element_line(size = 0.75),
        axis.text.x = element_text(size=Mediumfont,colour="black"),
        axis.text.y = element_text(size=Mediumfont,colour="black"),
        plot.margin = unit(Margin, "cm"),
        legend.position = "none",
        panel.background = element_blank())

grid.draw(Distribution_time_laying)
```

```
# svg(filename="F:/Dropbox/0_work/001_Res/0_Proj/006_Sex_select_daphnia_projects/04_presentation/Distribution_time_laying.svg",
#     width=5,
#     height=4,
#     pointsize=12)
# Distribution_time_laying
# dev.off()
```

**Do females in mating carry ephippia to be fertilized?**

```
Table_nbr_eggs_ephip = with(subset(all_data,Sex=="Female"),
                            as.data.frame(table(Nbr_eggs_in_ephippia)))
colnames(Table_nbr_eggs_ephip)[1] = "Number_of_eggs"
colnames(Table_nbr_eggs_ephip)[2] = "Counts"
rownames(Table_nbr_eggs_ephip) = c()
Table_nbr_eggs_ephip[,1]=c("Zero","One","Two")

Table_nbr_eggs_ephip$Proportion = c(Table_nbr_eggs_ephip[1,2]/(Table_nbr_eggs_ephip[2,2]+Table_nbr_eggs_ephip[1,2]+Table_nbr_eggs_ephip[3,2]),Table_nbr_eggs_ephip[2,2]/(Table_nbr_eggs_ephip[2,2]+Table_nbr_eggs_ephip[1,2]+Table_nbr_eggs_ephip[3,2]),Table_nbr_eggs_ephip[3,2]/(Table_nbr_eggs_ephip[2,2]+Table_nbr_eggs_ephip[1,2]+Table_nbr_eggs_ephip[3,2]))

Table_nbr_eggs_ephip$Proportion = round(Table_nbr_eggs_ephip$Proportion,digit=2)

plot_Table_nbr_eggs_ephip=
ggplot(Table_nbr_eggs_ephip,aes(x=
Number_of_eggs,y=Proportion))+
  geom_bar(stat="identity",fill="black",alpha=0.3,col="black",width=0.5)+
labs(caption = "(Figure 5A)")+
   scale_x_discrete("Nbr of sexual eggs\nin ephippium",
                    limits=c("Zero","One","Two"),
                    labels=c("0","1","2")) +
      scale_y_continuous("Proportion",
                         limits=c(0,1.05),
                         breaks=(seq(0,1,by=0.2)))+
  geom_text(aes(label=paste("(",Counts,")",sep="")), vjust=-0.5, color="black")+
  theme(axis.title.x = element_text(size = Mediumfont,colour="black"),
        axis.title.y = element_blank(),
        axis.line.x = element_line(colour="black",size=0.75),
        axis.line.y = element_line(colour="black",size=0.75),
        axis.ticks.x = element_line(size = 0.75),
        axis.ticks.y = element_line(size = 0.75),
        axis.text.x = element_text(size=Mediumfont,colour="black"),
        axis.text.y = element_text(size=Mediumfont,colour="black"),
        plot.margin = unit(c(0,0.2,0.2,0), "cm"),
       panel.background = element_blank())
plot_Table_nbr_eggs_ephip
```

```
Moment_laying = with(subset(all_data,Sex=="Female"),
                     as.data.frame(table(Laying_after_male_left_yes_no)))
Moment_laying = subset(Moment_laying,Laying_after_male_left_yes_no!="")
colnames(Moment_laying)[1] = "Female_laying_eggs_after_male_left"
colnames(Moment_laying)[2] = "Counts"
rownames(Moment_laying) = c()
Moment_laying[,1]=c("No","Yes")

Moment_laying$Proportion = c(Moment_laying[1,2]/(Moment_laying[2,2]+Moment_laying[1,2]),Moment_laying[2,2]/(Moment_laying[2,2]+Moment_laying[1,2]))

Moment_laying$Proportion = round(Moment_laying$Proportion,digit=2)

plot_Moment_laying=
ggplot(Moment_laying,aes(x=
Female_laying_eggs_after_male_left,y=Proportion))+
  geom_bar(stat="identity",fill="black",alpha=0.3,col="black",width=0.5)+
labs(caption = "(Figure 5B)")+
   scale_x_discrete("Females laying\neggs after male left",
                    limits=c("Yes","No"),
                    labels=c("True","False")) +
      scale_y_continuous("Proportion",
                         limits=c(0,1.05),
                         breaks=(seq(0,1,by=0.2)))+
  geom_text(aes(label=paste("(",Counts,")",sep="")), vjust=-0.5, color="black", size=3.5)+
  theme(axis.title.x = element_text(size = Mediumfont,colour="black"),
        axis.title.y = element_text(size = Mediumfont,colour="black"),
        axis.line.x = element_line(colour="black",size=0.75),
        axis.line.y = element_line(colour="black",size=0.75),
        axis.ticks.x = element_line(size = 0.75),
        axis.ticks.y = element_line(size = 0.75),
        axis.text.x = element_text(size=Mediumfont,colour="black"),
        axis.text.y = element_text(size=Mediumfont,colour="black"),
        plot.margin = unit(c(0,0.2,0.2,0), "cm"),
       panel.background = element_blank())
plot_Moment_laying
```

### 3 - Sperm morphology

#### 3.1 Sperm length

**Is there a difference in sperm length between males in the same mating? (Figure 6A)**  
Using 46 polyandrous mating from which we had sperm length for more than 30 sperms from the ejaculate of each males, we found that males from a same mating generally differ in average sperm length. More than 50% of our couples differed in average of more than 0.77 µm (i.e. 8.6 % larger than the averaged sperm length). (dotted red line represents the median) However, the order of detachment did not predict the average sperm size of the individuals. The second males had sperms on average of the same size than the first male.

[note: Urs is supposed to sequence, in October, few eggs to look if both eggs are fertilized by the same male or not.]

```
Data_sperm_diff = subset(all_data,!is.na(Total_sperm_length_µ_mean))
Data_sperm_diff= Data_sperm_diff[,c(1:9,22)]

Data_sperm_diff = 
 as.data.frame( Data_sperm_diff %>%
  group_by(ID_mating) %>%
    spread(Position_detached,Total_sperm_length_µ_mean))

colnames(Data_sperm_diff)[9]="Sperm_length_male_1"
colnames(Data_sperm_diff)[10]="Sperm_length_male_2"

Data_sperm_diff$Difference_sperm_length = 
  with(Data_sperm_diff,
       Sperm_length_male_1-Sperm_length_male_2
       )

mean_sperm_polyandrous = with(subset(all_data,!is.na(Total_sperm_length_µ_mean)),mean(Total_sperm_length_µ_mean))
relative_difference = median(Data_sperm_diff$Difference_sperm_length)/mean_sperm_polyandrous*100

Distribution_Sperm_difference=
  ggplot(Data_sperm_diff,aes(x=Difference_sperm_length)) +
  labs(caption = "(Figure 6A)")+
  geom_histogram(colour="black",fill="black",alpha=0.3,binwidth = 0.3)+
  scale_y_continuous("Count",
                     expand=c(0,0.1),
                     limits=c(0,6),
                     breaks=(c(seq(0,6,by=1))))+
  scale_x_continuous("Difference mean sperm length (µm)",
                     breaks=(seq(-3,3,by=0.5)))+
    # geom_vline(aes(xintercept=median(Data_sperm_diff$Difference_sperm_length)),
            # color="red", linetype="dashed", size=0.6)+
    geom_vline(aes(xintercept=0),
            color="red", linetype="dashed", size=0.6)+
  theme(axis.title.x = element_text(size=Mediumfont),
        axis.title.y = element_text(size=Mediumfont),
        axis.line.x = element_line(colour="black",size=0.75),
        axis.line.y = element_line(colour="black",size=0.75),
        axis.ticks.x = element_line(size = 0.75),
        axis.ticks.y = element_line(size = 0.75),
        axis.text.x = element_text(size=Mediumfont,colour="black"),
        axis.text.y = element_text(size=Mediumfont,colour="black"),
        plot.margin = unit(Margin, "cm"),
        legend.position = "none",
        strip.text.x = element_text(size =Mediumfont, colour = "black",face="italic"),
        strip.text.y = element_text(size =Mediumfont, colour = "black",face="italic"),
        strip.background = element_rect(color="white",fill=grey(0.90)),
        panel.background = element_blank())

grid.draw(Distribution_Sperm_difference)
```

```
# svg(filename="F:/Dropbox/0_work/001_Res/0_Proj/006_Sex_select_daphnia_projects/04_presentation/Distribution_Sperm_difference.svg",
#     width=5,
#     height=4,
#     pointsize=12)
# Distribution_Sperm_difference
# dev.off()
```

```
Full_model= fitme(Total_sperm_length~ ID_unique_male + (1|ID_couple), family=Gamma("log"),rand.family=gaussian("identity"), data=Data_sperm_2011)

model_reduced= fitme(Total_sperm_length~ 1 + (1|ID_couple), family=Gamma("log"),rand.family=gaussian("identity"), data=Data_sperm_2011)

test=anova(Full_model,model_reduced) 

Res_sperm_couple= data.frame(Variable=as.character("Order effect"),
                              chi2_LR=as.numeric(test$basicLRT$chi2_LR),
                              df=as.numeric(test$basicLRT$df),
                              Pvalue=as.numeric(test$basicLRT$p_value),
                              Effect=as.numeric(exp(Full_model$fixef[2])))

format(Res_sperm_couple,digit=2)%>%
  kable(col.names = c("", "chi^2 LRT", "df", "p-value", "Effect (ratio)")) %>%
  add_header_above(c("fitme(Total_sperm_length ~ Position_detached + (1|ID_mating), family=Gamma(log),rand.family=Gamma(log))" = 5))%>%
  kable_styling(bootstrap_options = c("striped", "hover", "condensed"), full_width = F)
```

| fitme(Total\_sperm\_length ~ Position\_detached + (1\|ID\_mating), family=Gamma(log),rand.family=Gamma(log)) | | | | |
| --- | --- | --- | --- | --- |
|  | chi^2 LRT | df | p-value | Effect (ratio) |
| Order effect | 3 | 1 | 0.082 | 1 |

---

#### 3.2 Genetic variation of sperm length

**Is there genetic variation for sperm length? (Figure 6B)**  
Sperm length in ejaculate of several males belonging to a same laboratory clone (4 different clones from Armenia, Cyprus, Germany and Russia). We were able to detect a strong difference between means of ejaculates.

```
mean_sperm_lab = aggregate(Sperm_length ~  as.factor(Clone), Sperm_length_lab, mean)
mean_sperm_lab$Sperm_length_mean = format(as.numeric(mean_sperm_lab$Sperm_length),digit=3)

sd_sperm_lab = aggregate(Sperm_length ~  as.factor(Clone), Sperm_length_lab, sd)
sd_sperm_lab$Sperm_length_sd = format(as.numeric(sd_sperm_lab$Sperm_length),digit=3)

sperm_lab = left_join(mean_sperm_lab[,c(1,3)],sd_sperm_lab[,c(1,3)])
sperm_lab$IDClone_ID = c("AMAR_4","CY-PA-1_4","Iinb1_4","RU-KOR-1_4")
sperm_lab$IDClone_ID_sd = c("AMAR_3","CY-PA-1_3","Iinb1_3","RU-KOR-1_3")

Plot_Sperm_length_lab=
  ggplot(Sperm_length_lab,aes(x=as.factor(Clone_ID),y=Sperm_length)) +
  geom_boxplot(aes(color=Clone),size =0.6,outlier.shape=NA)+
  geom_dotplot( aes(fill=Clone),alpha=0.3, binaxis = "y", stackdir = "center",dotsize=0.4, show.legend=FALSE)+
  labs(caption = "(Figure 6B)")+
  scale_y_continuous("Sperm length (µm)",
                     limits=c(0,33),
                     breaks=(seq(0,32,by=4)))+
  scale_x_discrete("Males")+
  scale_color_manual(name="Clones",
                     labels=c("AM-AR","CY-PA-1","DE-Iinb1","RU-KOR-1"),
                       values=c("#009E73","#E69F00","#0072B2", "#CC79A7" ))+
  scale_fill_manual(name="Clones",
                     labels=c("AM-AR","CY-PA-1","DE-Iinb1","RU-KOR-1"),
                       values=c("#009E73","#E69F00","#0072B2", "#CC79A7"))+
  stat_summary(fun.y = mean, geom = "point",colour="red",size=1) +
  geom_text(data=sperm_lab,aes(x=IDClone_ID,y=31,
                               label=paste("mean= ",Sperm_length_mean,sep="")),colour="red",size=3)+
  geom_text(data=sperm_lab,aes(x=IDClone_ID_sd,y=28,
                               label=paste("sd= ",Sperm_length_sd,sep="")),colour="red",size=3)+
  theme(axis.title.x = element_text(size = Mediumfont,colour="black"),
        axis.title.y = element_text(size = Mediumfont,colour="black"),
        axis.line.x = element_line(colour="black",size=0.75),
        axis.line.y = element_line(colour="black",size=0.75),
        axis.ticks.x = element_line(size = 0.75),
        axis.ticks.y = element_line(size = 0.75),
        axis.text.x = element_text(size=Mediumfont,colour="white"),
        axis.text.y = element_text(size=Smallfont,colour="black"),
        plot.margin = unit(Margin, "cm"),
       panel.background = element_blank(),
        legend.direction = "horizontal", 
        legend.box = "vertical",
        legend.position = "top",
        legend.key.height = unit(0.4, "cm"),
        legend.key.width= unit(0.3, "cm"),
        legend.title = element_text(face="italic",size=Mediumfont), 
        legend.key = element_rect(colour = 'white', fill = "white", linetype='dashed'),
        legend.text = element_text(size=Mediumfont),
        legend.background = element_rect(fill=NA))

grid.draw(Plot_Sperm_length_lab)
```

```
# svg(filename="F:/Dropbox/0_work/001_Res/0_Proj/006_Sex_select_daphnia_projects/04_presentation/Plot_Sperm_length_lab.svg",
#     width=5,
#     height=4,
#     pointsize=12)
# Plot_Sperm_length_lab
# dev.off()
```

```
model_full= fitme(Sperm_length~Clone + (1|Clone/ID),resid.model= ~ Clone, family=Gamma("log"),rand.family=Gamma("log") ,data=Sperm_length_lab)

model_1= fitme(Sperm_length~1 + (1|Clone/ID),resid.model= ~ Clone, family=Gamma("log"),rand.family=Gamma("log") ,data=Sperm_length_lab)

test=anova(model_full,model_1) #variance between clones is larger than within as there is a clone effect

Res_sperm_lab= data.frame(Variable=as.character("Clone effect"),
                              df=as.numeric(test$basicLRT$df),
                              chi2_LR=as.numeric(test$basicLRT$chi2_LR),
                              Pvalue=as.numeric(test$basicLRT$p_value))

format(Res_sperm_lab,digit=2)%>%
  kable(col.names = c("",  "df","chi^2 LRT", "p-value")) %>%
  add_header_above(c("fitme(Sperm_length~Clone + (1|Clone/ID),\nresid.model= ~ Clone, family=Gamma(log),rand.family=Gamma(log))" = 4))%>%
  kable_styling(bootstrap_options = c("striped", "hover", "condensed"), full_width = F)
```

| fitme(Sperm\_length~Clone + (1\|Clone/ID), resid.model= ~ Clone, family=Gamma(log),rand.family=Gamma(log)) | | | |
| --- | --- | --- | --- |
|  | df | chi^2 LRT | p-value |
| Clone effect | 3 | 17 | 6e-04 |

```
Variable = c("Clone","","","")
Var_log_inter_clone = c(format(model_1$lambda[2],digit=3),"","","")
Clones=c("AMAR","CY-PA-1","Iinb1","RU-KOR-1")
Var_log_intra_clone = format(c(model_full$phi.object$fixef[1],model_full$phi.object$fixef[2]+model_full$phi.object$fixef[1],model_full$phi.object$fixef[3]+model_full$phi.object$fixef[1],model_full$phi.object$fixef[4]+model_full$phi.object$fixef[1]),digit=3)

tab_var_sperm_lab = cbind(Variable,Var_log_inter_clone,Clones,Var_log_intra_clone)

tab_var_sperm_lab%>%
  kable(row.names = F,col.names = c("",  "Variance log - inter","Clones","Variance log - intra"))  %>%
  add_header_above(c(" " = 4))%>%
  kable_styling(bootstrap_options = c("striped", "hover", "condensed"), full_width = F)
```

---

#### 3.3 Description of sperm length variation

```
img2 <- readImage("F:/Dropbox/0_work/001_Res/0_Proj/006_Sex_select_daphnia_projects/illustration_sperm_Dmagna.jpg") 
grid.raster(img2)
```

Ejaculate of a *Daphnia magna* male (Figure 4D inlet)

---

**What is the sperm length variation in an ejaculate? (Figure 6C)**

**What is the sperm length distribution? (Figure 6D)**  
We found that sperm length distribution in an ejaculate was following a Gamma distribution. Based on the AIC to evaluate the quality of fit of the model to explain our data (accumulation of likelihood computed withing individuals ejaculates), the Gamma distribution (AIC=12096.67) was better to explain sperm length than a simple Gaussian (AIC= 12218.56) or a combination of two Gaussian (AIC= 12613.08). This suggests that the scenario with binary phenotypes with different functions is less parcimonious than a scenario with one category of sperm in which the length vary unequally around the mean (red dashed line). The qqplot also shows that a gamma distribution is more likely.

```
 mean_sperm = aggregate(Total_sperm_length_µ_mean ~  1, Data_male_couple_2011, mean)
 sd_sperm = aggregate(Total_sperm_length_µ_sd ~  1, Data_male_couple_2011, mean)

mean_sperm_tot=mean(Data_sperm_2011$Total_sperm_length,na.rm=T)
sd_sperm_tot=sd(Data_sperm_2011$Total_sperm_length,na.rm=T)

####

sperm_size_order=Data_mean_sperm_size[order(Data_mean_sperm_size$Total_sperm_length_µ_median),]
rank_males = sperm_size_order$ID_unique_male

Ranked_Sperm_size=
  ggplot(Data_sperm_2011,aes(x=as.factor(ID_unique_male),y=Total_sperm_length)) +
  labs(caption = "(Figure 6C)")+
  geom_boxplot(aes(color=Population),size =0.6,outlier.shape=NA)+
  geom_dotplot( aes(fill=Population),alpha=0.3, binaxis = "y", stackdir = "center",binwidth = 0.1, show.legend=FALSE)+
  scale_y_continuous("Sperm length",
                     limits=c(0,20),
                     breaks=(seq(0,20,by=4)))+
  scale_x_discrete("Males",
                   limits=rank_males)+
  annotate("text",label="mean all sperm length= 9.04µm \u00B1 2.2 \nmean of standard deviation per ejaculate = 1.9",x=20, y=2,size=4,
             hjust = 0)+
  stat_summary(fun.y = median, geom = "point",colour="red",size=0.8) +
  geom_hline(aes(yintercept=mean(Total_sperm_length)),color="black", linetype="dashed", size=0.6)+
  theme(axis.title.x = element_text(size=Mediumfont),
        axis.title.y = element_text(size=Mediumfont),
        axis.line.x = element_line(colour="black",size=0.75),
        axis.line.y = element_line(colour="black",size=0.75),
        axis.ticks.x = element_line(size = 0.75),
        axis.ticks.y = element_line(size = 0.75),
        axis.text.x = element_text(size=Mediumfont,colour="white"),
        axis.text.y = element_text(size=Mediumfont,colour="black"),
        plot.margin = unit(Margin, "cm"),
        strip.text.x = element_text(size =Mediumfont, colour = "black",face="italic"),
        strip.text.y = element_text(size =Mediumfont, colour = "black",face="italic"),
        legend.direction = "vertical", 
        legend.box = "horizontal",
        legend.position = "none",
        legend.key.height = unit(0.4, "cm"),
        legend.key.width= unit(0.3, "cm"),
        legend.title = element_text(face="italic",size=Smallfont), 
        legend.key = element_rect(colour = 'white', fill = "white", linetype='dashed'),
        legend.text = element_text(size=Smallfont),
        legend.background = element_rect(fill=NA),
        panel.background = element_blank())+
  guides(shape=guide_legend(ncol=1),
         fill=guide_legend(ncol=1),
         col=guide_legend(ncol=1))

# svg(filename="F:/Dropbox/0_work/001_Res/0_Proj/006_Sex_select_daphnia_projects/04_presentation/Ranked_Sperm_size.svg",
#     width=5,
#     height=4,
#     pointsize=12)
# Ranked_Sperm_size
# dev.off()
```

```
fit.gamma <- fitdist(Data_sperm_2011$Total_sperm_length, "gamma", lower = c(0, 0))
fit.normal <- fitdist(Data_sperm_2011$Total_sperm_length, "norm")

#par(mfrow = c(2, 2))
plot.legend <- c( "Gamma","Gaussian")
#denscomp(list( fit.gamma,fit.normal), fitcol = c("red", "blue"), legendtext = plot.legend)
#qqcomp(list(fit.gamma,fit.normal), fitcol = c("red", "blue"), legendtext = plot.legend)
#cdfcomp(list(fit.gamma,fit.normal), fitcol = c("red", "blue"), legendtext = plot.legend)
#ppcomp(list(fit.gamma,fit.normal), fitcol = c("red", "blue"), legendtext = plot.legend)

list_unique_males= unique(Data_sperm_2011$ID_unique_male)
Tab_AIC_sperm = data.frame(IDmale=list_unique_males,GoF_Gamma=NA, GoF_Gausssian=NA, AIC_Gamma=NA, AIC_Gaussian=NA)
for (i in 1:length(list_unique_males)){
ID.male= list_unique_males [i]

fit.params_indiv.gamma <- with(subset(Data_sperm_2011,ID_unique_male==ID.male),
  fitdist(Total_sperm_length, "gamma", lower = c(0, 0)))
fit.params_indiv.norm<- with(subset(Data_sperm_2011,ID_unique_male==ID.male),
  fitdist(Total_sperm_length, "norm", lower = c(0, 0)))

GoF = gofstat(list(fit.params_indiv.gamma, fit.params_indiv.norm))

Tab_AIC_sperm$ID_male[i] = ID.male
Tab_AIC_sperm$AIC_Gamma[i] = GoF$aic[1]
Tab_AIC_sperm$AIC_Gaussian[i] = GoF$aic[2]
}

#sum(Tab_AIC_sperm$AIC_Gamma) -sum(Tab_AIC_sperm$AIC_Gaussian) # AIC Gamma - AIC Gaussian and AIC Gamma is better. Sum represents the sum of the AIC per individual

mix_mod_gauss = densityMclust(Data_sperm_2011$Total_sperm_length,modelNames="V")
AIC_mixModGaus = 2*logLik(mix_mod_gauss)+2*5 # n= 5 because 2 variances, 2 means and 1 proportion

#sum(Tab_AIC_sperm$AIC_Gamma) - (-AIC_mixModGaus[1]) # AIC gamma is better than mixte gaussian

# which(Tab_AIC_sperm$AIC_Gamma - Tab_AIC_sperm$AIC_Gaussian=-2 )
# length(which(Tab_AIC_sperm$AIC_Gamma<=Tab_AIC_sperm$AIC_Gaussian))
```

```
Distribution_Sperm_size=
  ggplot(Data_sperm_2011,aes(x=Total_sperm_length,y=..density..)) +
labs(caption = "(Figure 6D)")+
  geom_histogram(alpha=0.3,binwidth = 0.2,color="black")+
  #facet_grid(Population~.)+
     geom_line(aes(Data_sperm_2011$Total_sperm_length, y=dgamma(Data_sperm_2011$Total_sperm_length,fit.gamma$estimate["shape"], fit.gamma$estimate["rate"])), color="red", size = 1) + 
     geom_line(aes(Data_sperm_2011$Total_sperm_length, y=dnorm(Data_sperm_2011$Total_sperm_length,mean(Data_sperm_2011$Total_sperm_length), sd(Data_sperm_2011$Total_sperm_length))), color="blue", size = 1) + 
  annotate("text",x=6,y=0.18,label="Gamma",col="red")+
  annotate("text",x=12.5,y=0.14,label="Gaussian",col="blue")+
  scale_x_continuous("Sperm length",
                     breaks=(seq(0,20,by=2)))+
  geom_hline(aes(yintercept=0),
             color="black", linetype="solid", size=0.4)+
  geom_vline(aes(xintercept=mean(Total_sperm_length)),
             color="red", linetype="dashed", size=0.6)+
  theme(axis.title.x = element_text(size=Mediumfont+4),
        axis.title.y = element_text(size=Mediumfont+4),
        axis.line.x = element_line(colour="black",size=0.75),
        axis.line.y = element_line(colour="black",size=0.75),
        axis.ticks.x = element_line(size = 0.75),
        axis.ticks.y = element_line(size = 0.75),
        axis.text.x = element_text(size=Mediumfont,colour="black"),
        axis.text.y = element_text(size=Mediumfont,colour="black"),
        plot.margin = unit(Margin, "cm"),
        legend.position = "right",
        strip.text.x = element_text(size =Mediumfont+4, colour = "black",face="italic"),
        strip.text.y = element_text(size =Mediumfont+4, colour = "black",face="italic"),
        strip.background = element_rect(color="white",fill=grey(0.90)),
        panel.background = element_blank())

#grid.draw(Distribution_Sperm_size)

# svg(filename="F:/Dropbox/0_work/001_Res/0_Proj/006_Sex_select_daphnia_projects/04_presentation/Distribution_Sperm_size.svg",
#     width=5,
#     height=4,
#     pointsize=12)
# Distribution_Sperm_size
# dev.off()

qqcomp(list(fit.gamma,fit.normal), fitcol = c("red", "blue"), legendtext = plot.legend)
```

```
qqcomp_spermlength = grab_grob()
```

```
qqcomp_spermlength = as.ggplot(qqcomp_spermlength)
```

```
first_row = plot_grid(Ranked_Sperm_size)
second_row = plot_grid(Distribution_Sperm_size, qqcomp_spermlength,  nrow = 1)
plot_grid(first_row, second_row, labels=c('', ''), ncol=1)
```

---


---


---

### References

Booksmythe, Isobel, Nina Gerber, Dieter Ebert, and Hanna Kokko. 2018. “Daphnia females adjust sex allocation in response to current sex ratio and density.” Edited by Lutz Becks. *Ecology Letters* 21 (5). John Wiley & Sons, Ltd (10.1111): 629–37. doi:10.1111/ele.12929.

Ebert, Dieter, J W Hottinger, and V I Pajunen. 2001. “Temporal and spatial dynamics of parasite richness in a \(\backslash\)emph{Daphnia} metapopulation.” *Ecology* 82 (12): 3417–34. doi:10.1890/0012-9658(2001)082[3417:TASDOP]2.0.CO;2.

Jiang, Yuexin, Daniel I Bolnick, and Mark Kirkpatrick. 2013. “Assortative Mating in Animals.” *The American Naturalist* 181 (6): E125–E138. doi:10.1086/670160.

Lass, Sandra, and Dieter Ebert. 2006. “Apparent seasonality of parasite dynamics: Analysis of cyclic prevalence patterns.” *Proceedings of the Royal Society B: Biological Sciences* 273 (1583): 199–206. doi:10.1098/rspb.2005.3310.

Roth, Olivia, Dieter Ebert, Dita B. Vizoso, Annette Bieger, Sandra Lass, Bieger Annette, and Sandra Lass. 2008. “Male-biased sex-ratio distortion caused by <i>Octosporea bayeri</i>, a vertically and horizontally-transmitted parasite of <i>Daphnia magna</i>.” *International Journal for Parasitology* 38 (8-9). Pergamon: 969–79. doi:10.1016/j.ijpara.2007.11.009.
